# Probing cellular arrhythmogenesis using the O’Hara-Rudy model of the undiseased human ventricular cardiomyocyte

**DOI:** 10.1101/2020.11.15.384032

**Authors:** Gianluca Selvaggio, Wan Hongbin, Robert A. Pearlstein

## Abstract

The ventricular action potential (AP) is subserved by an interdependent system of voltage-gated ion channels and pumps that both alter and respond (directly or indirectly) to the dynamic transmembrane potential (Δ*ψ_m_*(*t*)) via voltage-dependent state transitions governing inward and outward ion currents. The native dynamic inward-outward current balance is subject to disruption caused by acquired or inherited loss or gain of function in one or more ion channels or pumps. Building on our previous work, we used a modified version of the O’Hara-Rudy (ORd) model of the undiseased human ventricular cardiomyocyte to study the pro-arrhythmic effects of three types of arrhythmia-inducing perturbations in midmyocytes (M cells):

1. Blockade of the human ether-a-go-go related gene (hERG) K^+^ channel introduced via a Markov state binding model.
2. Mutation-induced voltage shifts in hERG channel gating, resulting in faster inactivation or slowed recovery of both phosphorylated and non-phosphorylated forms of the channel (known as LQT2 syndrome).
3. Mutation-induced voltage shifts in Na_v_1.5 gating, resulting in slowed late inactivation of the phosphorylated and non-phosphorylated forms of the channel (known as LQT3 syndrome).

We studied the relationships between ion current anomalies and AP morphology as a function of cycle length (CL) and perturbation type/level. The results are summarized as follows:

1. AP duration (APD) is governed directly by Kir2.1 activation (I_K1_), which is delayed when repolarization is slowed by abnormal net inward tipping of the dynamic inward-outward current balance (reflected in decreased *d*(Δ*ψ_m_*(*t*))/*dt* during the late AP repolarization phase). In the case of hERG blockade by non-trappable compounds, the perturbation level consists of the dynamic fractional occupancy of the channel, which is governed by blocker k_on_ relative to the rate of channel opening, pharmacokinetic exposure, and k_off_ (in that order).
2. Arrhythmia progresses from prolonged paced APs → atypical APs (spontaneous and paced) → self-sustaining oscillations. Abrupt transitions between these regimes occur at CL- and perturbation-specific thresholds (denoted as T_1_, T_2_, and T_3_, respectively), whereas intra-regime progression proceeds in a graded fashion toward the subsequent threshold. APD and *d*(Δ*ψ_m_*(*t*))/*dt* during the late repolarization phase varied significantly across the 200 APs of our simulations near the T_1_ threshold at CL = 1/35 min, reflecting increasing instability of the AP generation system.
3. Arrhythmic APs exhibit highly variable cycle-to-cycle morphologies, depending on the perturbation level, type, and phasing between the underlying ion channel states and pacing cycle.
4. Atypical APs may be triggered by typical or atypical depolarizations prior to the T_3_ threshold, depending on perturbation type/level and phasing relative to CL:

a. APD/CL resides outside of the Goldilocks zone:

i. APD/CL → 1 at shorter CL and/or longer APD, resulting in pro-arrhythmic “collisions” between successive paced APs (AP_i_ and AP_j_) within a given cardiomyocyte. We studied this scenario at 60 and 80 beats per minute (BPM), equating to CL = 1/60 and 1/80 min.
ii. APD/CL < 1 at longer CL results in <u>spontaneous</u> atypical depolarizations within prolonged paced APs at elevated takeoff Δψ_m_(t) and increased channel phosphorylation levels. We studied this scenario at CL = 1/35 min.
b. APD and *d*(Δ*ψ_m_*(*t*))/*dt* during the late repolarization phase become increasingly variable over successive APs on approach to the T_1_ threshold, which is the possible source of short-long-short sequences observed in the ECG preceding torsades de pointes arrhythmia (TdP).
5. All atypical depolarizations are solely Ca_v_1.2 (I_Ca,L_)-driven (Δ*ψ_m_*(*t*) falls within the Na_v_1.5 <u>inactivation</u> window), whereas typical depolarizations are Na_v_1.5 (I_Na_) + I_Ca,L_-driven. Atypical depolarization versus typical repolarization occurrences are determined by the faster of Ca_v_1.2 and Kir2.1 (I_K1_) activation (where I_K1_ becomes increasingly dampened as the minimum Δ*ψ_m_*(*t*) drifts above the Kir2.1 activation window).
6. Ca_v_1.2 inactivation gates reset to the open position (accompanied by recovery) synchronously with channel closing under control conditions, generating a small ICa,L window current in the process. This current grows toward a depolarizing spike when the lag time between recovery and closing grows above a threshold level.
7. APs undergo damped oscillatory Ca_v_1.2 recovery/re-inactivation cycles above the T_3_ threshold, which are refreshed by subsequent pacing signals (nodal or reentrant in origin).

## Introduction

As for all cellular systems, the AP generation systems of excitable cells, including cardiomyocytes, operate within the non-equilibrium/non-linear dynamics regime [1]. Stable APs depend on tight dynamic counter-balancing between inward and outward ion currents, such that Δ*ψ_m_*(*t*) neither under-nor overshoots into the proarrhythmic range. Atypical APs, referred to as early afterdepolarizations (EADs), result from inward-outward ion current imbalances caused by acquired or inherited loss or gain of function in outwardly or inwardly conducting voltage-gated ion channels (notably hERG, KCNQ1, and Kir2.1 versus Na_v_1.5 and Ca_v_1.2) or pumps. EADs may progress further to self-sustaining cellular and organ level tachyarrhythmias (denoted as CAs and OAs, respectively), including TdP [2–5]. The molecular causes of CA and OA include:

1. Prolongation of AP duration (APD) in inherited long QT syndromes (manifesting as chronic QT prolongation in the ECG) caused by mutations in hERG (LQT2), Na_v_1.5 (LQT3), or Ca_v_1.2 channels (LQT8).
2. Loss of outward Kir2.1 current (I_K1_) [6], accompanied by gain of ryanodine receptor (RyR_2_)-mediated Ca^2+^ release [7] in heart failure, resulting in delayed afterdepolarizations (DADs) and ventricular fibrillation [2,3].
3. Gains in Ca_v_1.2 current (I_Ca,L_), late Na_v_1.5 current (I_Na(late)_), or the Ca^2+^ transient, combined with reduced repolarizing K^+^ currents, resulting in APD prolongation and EADs in human hypertrophic cardiomyopathy (HCM) [8,9].
4. Loss of I_K1_ and increased Ca^2+^ transient, resulting in rabbit tachyarrhythmia (suggesting a synergistic relationship between these factors) [10].

Understanding the detailed mechanisms by which atypical APs arise is critical for drug safety assessment and the development of therapies targeted at LQT syndromes. Ion channel and electrical dysfunction have been well studied experimentally in the *in vitro, ex vivo,* and *in vivo* settings using electrophysiology (e.g. patch clamp) and biochemical approaches, monophasic action potential (AP) recordings in isolated hearts and tissue preparations, and electrocardiography, respectively. However, studies at the cellular level have been hampered by the lack of stable human myocardial cell lines that fully recapitulate normal and abnormal APs under *in vivo*-like conditions, resulting in the need for high quality *in silico* surrogates such as the O’Hara-Rudy (ORd) simulation of the undiseased human ventricular (endo-, epi-, and midmyo) cardiomyocyte [11]. In our previous work, we used a modified version of the ORd model (denoted as ORd/hERG Markov [12]) to simulate the dynamic state distribution of the hERG potassium channel, and characterize the relationships between blocker binding kinetics (BK), dynamic hERG blocker occupancy, and EADs [12] (noting that CA is an emergent, rather than programmed, behavior of this model). We later used ORd/hERG Markov to study the mechanisms of positive and negative hERG channel potentiation [13]. Here, we again use this model to investigate the detailed mechanisms of arrhythmogenesis in M cells under conditions of 1) hERG channel blockade; 2) LQT2 syndrome, which we simulated by speeding and slowing hERG inactivation and recovery, respectively; and 3) LQT3 syndrome, which we simulated by slowing inactivation of late Na_v_1.5 current (denoted as I_Na(late)_). We set about to characterize the arrhythmogenesis process as a function of perturbation type and level, focusing on:

1. The chronic reversible sub-arrhythmic regime, as exists in the at-risk patient population, versus the acute arrhythmic regime (the point of no return).
2. Perturbation-induced alterations in ion currents, voltage, Ca^2+^, and other arrhythmogenic factors.
3. Perturbation severity-response relationships, including hERG blocker dose-response, and ion channel gain or loss of inactivation function-response relationships.

## Materials and methods

We previously generated the ORd/hERG Markov model via substitution of the original Hodgkin-Huxley-based hERG formulation with a published Markov formulation ([14] and Figure 3 of [12]). All ORd/hERG Markov simulations were performed in MATLAB™ 2017b (The MathWorks Inc., Natick, MA). The major components of the AP-generation system considered in this work are shown in Figure 1 (adapted from [11]).

**Figure 1.**
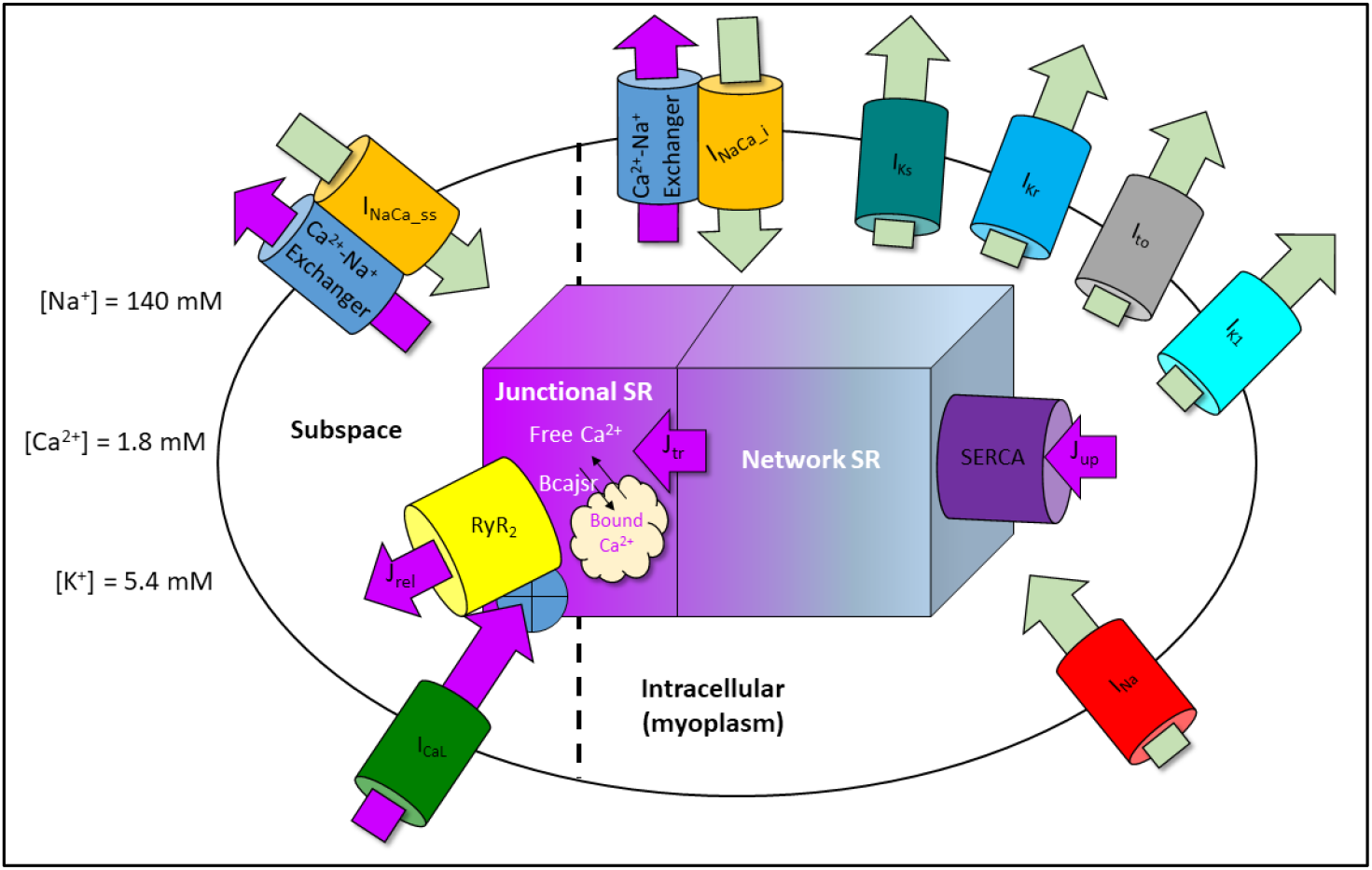
The major channels/pumps/currents considered in this work. Ca^2+^, Na^+^, and K^+^, which are maintained in dynamic non-equilibrium distributions by ion pumps/exchangers, translocate down their electrochemical gradients (toward equilibrium) in response to ion channel activation. Inward and outward transmembrane Ca^2+^ and Na^+^ fluxes occur via voltage-gated channels and intracellular and subspace NCX exchangers (in which the outward Ca^2+^ flux is driven by the transmembrane Na^+^ gradient). Free Ca^2+^ entry/exit to/from the sarcoplasmic reticulum (SR) is driven by the trans-SR concentration gradient. The rates of entry, intra-SR translocation, and exit (J_up_, J_tr_, and J_rel_) are governed by free energy barriers.

### hERG blockade

As in our previous work [12], we used a <u>non-trappable</u> hERG blocker exhibiting k_on_ 10^8^ M^-1^s^-1^, k_off_ = 2.0 s^-1^, and IC_50_ = 20 nM to probe the arrhythmogenesis mechanism in M cells (chosen based on the lower hERG density and concomitantly greater sensitivity of these cells to perturbations compared with their epi-and endocardial counterparts [12]). We calculated the constant fractional occupancy of <u>trappable</u> blockers based on the open + inactivated + closed state populations, and the dynamic fractional occupancy of <u>non-trappable</u> blockers from the <u>occupied</u> open + inactivated states relative to the <u>total</u> open + inactivated state populations. We then used the putative pro-arrhythmic criteria derived from our study to characterize hERG blocker dose-response relationships for several BK profiles at cycle length (CL) = 1/35, 1/60, and 1/80 min. The hERG IC_50_ was maintained in all cases at 1.0 μM, the typical upper practical limit of hERG potency visa-vis the Redfern safety index (SI) [15] (noting that the upper therapeutic free C_max_ in humans at this IC_50_ is constrained by the Redfern SI to ~33 nM).

We developed an automated algorithm to identify the BK and CL-dependent hERG blocker concentrations at which APs transition from the pre-arrhythmic regime (denoted as the PT_1_ threshold) to the arrhythmic regime (denoted as the T_1_ threshold), which is approached asymptotically as a function of fractional decrements in blocker concentration (loosely analogous to the equivalence point in acid-base titrations). The algorithm consists of ORd/hERG Markov simulations performed over a series of blocker concentrations, beginning with a user-specified initial guess (C_i_ < PT_1_), which was then incremented in steps (denoted as s) of 0.1 · C_i_ until the T_1_ threshold is crossed, at which point the concentration was reset to the prior sub-T_1_ concentration, the step size reduced by s/4.0, and the process repeated. We set the convergence criterion to s < 0.001, combined with the absence of atypical depolarizations during each simulation.

### LQT2

We simulated pro-arrhythmia and arrhythmia associated with loss of the hERG current (denoted as I_Kr_) by speeding hERG inactivation and slowing recovery from inactivation, as follows:

**Inactivated → open state transition (denoted as *α_i_* in the hERG Markov model) (Figure 2A):**

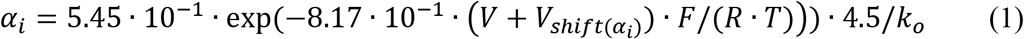

where *V* is the transmembrane voltage, *V*_*shift*(*α_i_*)_ is a voltage shift factor used to slow this state transition, F is the Faraday constant, R is the gas constant, T is the temperature, and *k_o_* is the extracellular K^+^ concentration.

**Figure 2.**
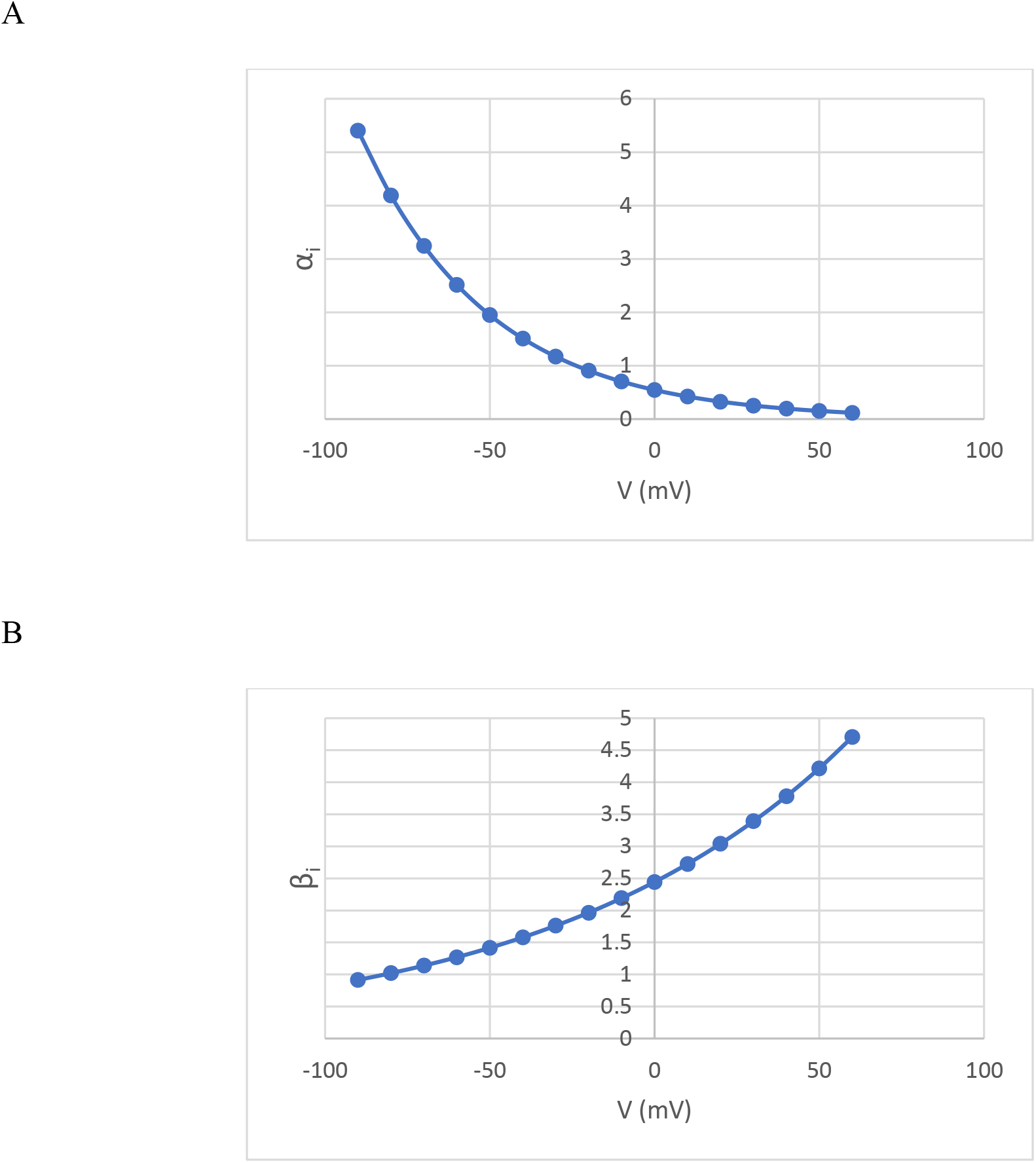
(A) Plot of the voltage dependence of the hERG open → inactivated state transition (*α_i_*) with *V*_*shift*(*α_i_*)_ = 0. The channel population is strongly inactivated during depolarization. (B). Plot of the voltage dependence of the hERG inactivated → open state transition (recovery from inactivation) (*β_i_*) with *V*_Shift(*β_i_*)_ = 0. The channel population undergoes rapid recovery during repolarization (plots of these state transitions are provided in Figure 1 of [12]).

**Open → inactivated state transition (denoted as *β_i_* in the hERG Markov model) (Figure 2B):**

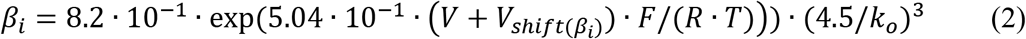

where *V*_*shift*(*β_i_*)_ is a voltage shift factor used to speed this state transition. We then studied arrhythmogenesis as a function of increasing *V*_*shift*(*α_i_*)_ and *V*_*Shift*(*β_i_*)_.

### LQT3

We simulated pro-arrhythmia and arrhythmia associated with increased I_Na(late)_ by slowing the late inactivation gate of Na_v_1.5 (denoted as *h_L_*), which is calculated in the ORd and ORd hERG/Markov models via integration of:

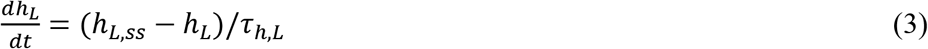

where *h_L,ss_* is the voltage-dependent term (Figure 3A), and *τ_h,L_* is the gating time step = 200 ms. A voltage shift term (*V_shift_* ≤ 0) introduced to *h_L,ss_* was used to slow inactivation, as follows (Figure 3B):

**Figure 3.**
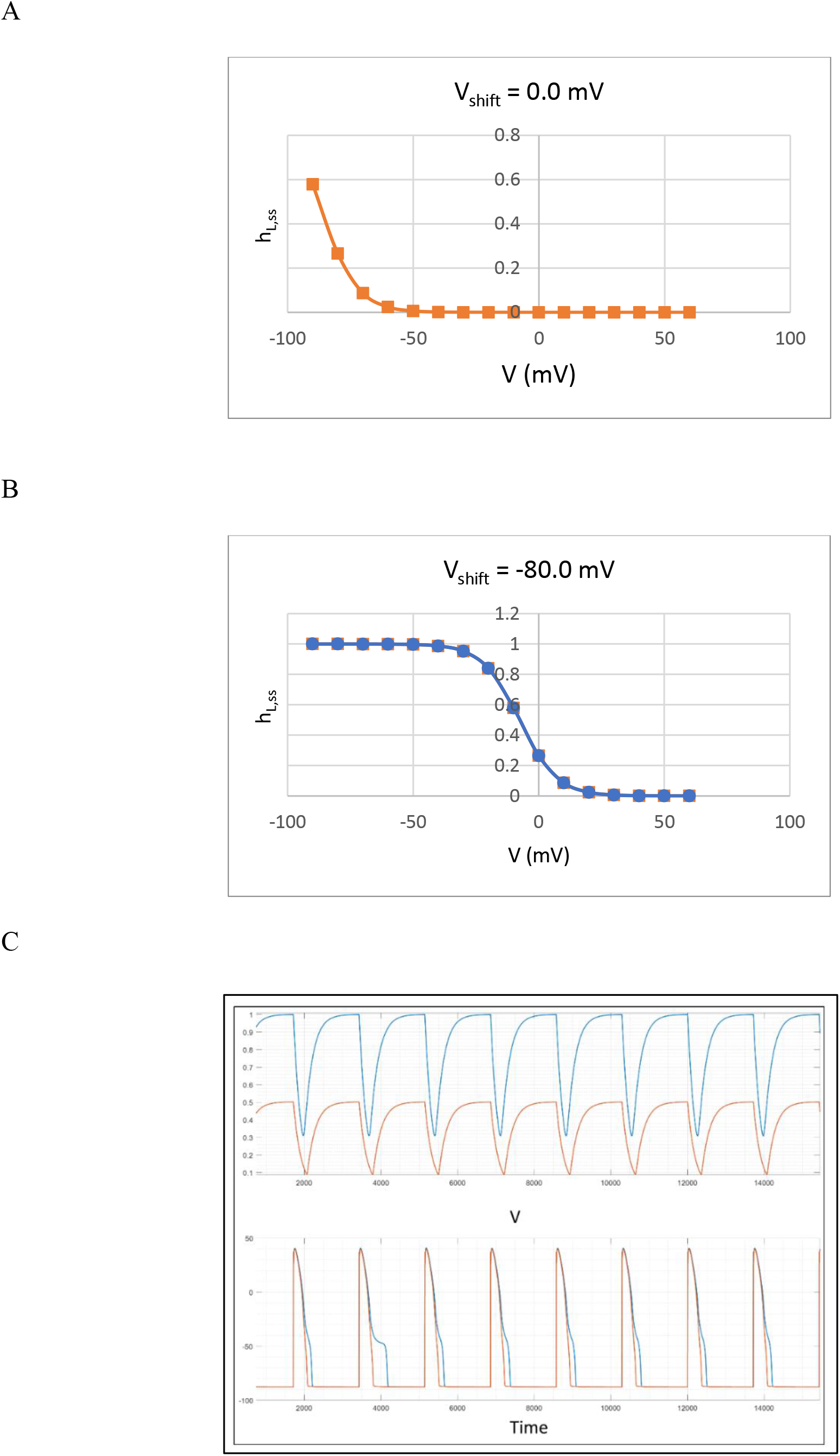
(A) Plot of *h_L,ss_* for the wild type non-phosphorylated Na_v_1.5 population. Late inactivation occurs at voltages ≽ −40 mV, reaching ~50% maximal inactivation at −90 mV (noting I_Na(late)_ results from partial inactivation of the channel population). (B) Plot of *h_L,ss_* for the mutant non-phosphorylated channel population, exemplified by an arbitrary *V_shift_* = −80 mV, which results in late Na_v_1.5 inactivation at voltages ≽ +30 mV, reaching zero inactivation prior to closing at voltages ≼ −50 mV. (C) Upper panel: The simulated *h_L_* gate for wild type (*V_shift_* = 0.0) and mutant (exemplified with *V_shift_* = −74.1893 mV) channels (orange and light blue tracings, respectively), noting a decrease in overall channel inactivation throughout the AP. Lower panel: APs corresponding to control and perturbed conditions (orange and light blue tracings, respectively).

Non-phosphorylated:

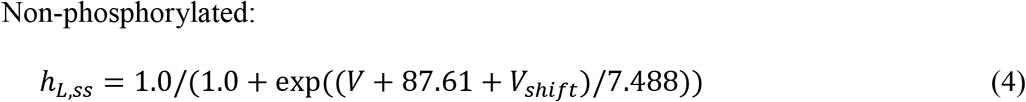

Phosphorylated:

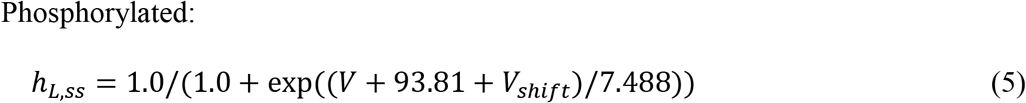

We then studied arrhythmogenesis as a function of increasing |*V_shift_*| (the modified equation 3 is exemplified in Figure 3C for *V_shift_* = −74.1893 mV).

### Ion current knockouts/knockdowns

I_Kr_ and I_K1_ (activated during phases 0 and 3-4 [16], denoted as I_K1(early)_ and I_K1(late)_, respectively) knockouts were generated by fixing their values at 0. CaMKII knockdown was generated by fixing the K_M_ (KmCaMK) to 10.0 (default = 0.15).

### Isolation of individual contributions to I_Ca,L_ and I_Na_

Ca_v_1.2 and Na_v_1.5 currents were calculated separately for each of their individual activation and inactivation mechanisms based on the following expressions taken from [17]:

#### Calculation of I_Ca,L_

**Activation:**

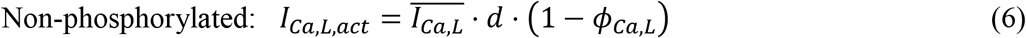

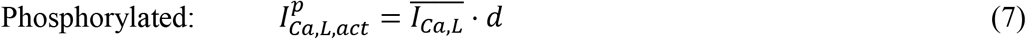

**Inactivation:**

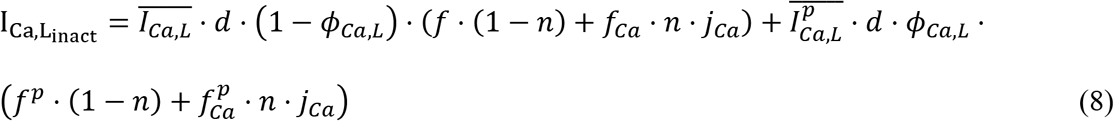

where 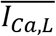 and 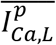 are the maximum currents through non-phosphorylated and phosphorylated Ca_v_1.2, respectively; *d* is the rate of channel activation; *n* and (1 – *n*) denote the fraction of channels undergoing Ca^2+^- (CDI) and voltage-dependent inactivation (VDI), respectively; (1 – *Φ_Ca,L_*) and *φ_Ca,L_* represent the fraction of non-phosphorylated and CaMKII-phosphorylated channels, respectively; *j_Ca_* denotes the recovery rate from Ca^2+^-dependent inactivation; and *f* and *f^p^*, and *f_Ca_* and 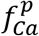 denote the rates of VDI and CDI for non-phosphorylated and phosphorylated channels, respectively. Ca_v_1.2 channels exhibit fast and slow dynamics for both VDI and CDI, consisting of the following contributions:

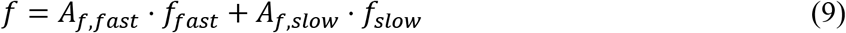

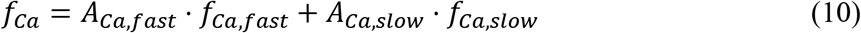

where *A_f,fast_* (= 0.6) is the fraction of channels undergoing fast VDI, *A_f,stow_* = 1 — *A_f,fast_* is the fraction of channels undergoing slow VDI, *A_Ca,fast_* (= 0.3 + (0.6/(1.0 + exp(V — 10)/10)) is the fraction of channels undergoing fast CDI, and *A_Ca,slow_* = 1 — *A_Ca,fast_* is the fraction of channels undergoing slow CDI. I_Ca,L_ is thus separable into k currents (I_CaL_inact,k__) governed by slow/non-phosphorylated (*I_V,s_, I_VCa,s_*), fast/non-phosphorylated (*I_V,f_, I_VCa,f_*), slow/phosphorylated 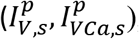, and fast/phosphorylated 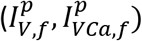 CDI and VDI contributions (eight in total). As we show later in this work, anomalous depolarizations are mediated largely by 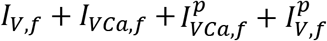 (denoted as I_Ca,L(late)_).

#### Calculation of peak I_Na_ (denoted as I_Na(early)_)

**Activation:**

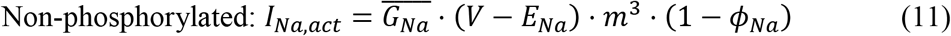

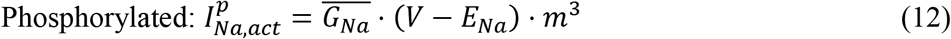

**Inactivation:**

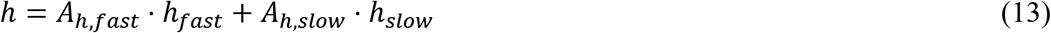

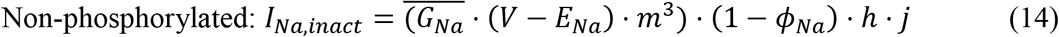

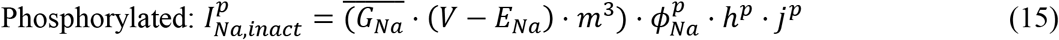

#### Calculation of I_Na(late)_

**Activation:**

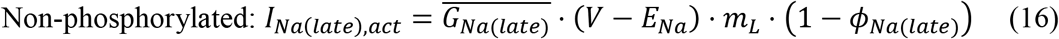

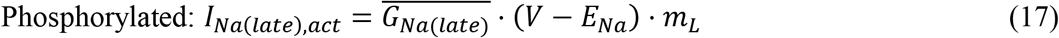

**Inactivation:**

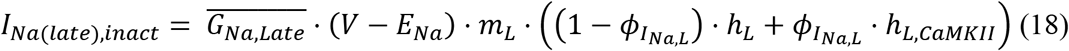

where 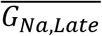 is the maximum conductance of the single channel, *m_L_* is the activation rate, *h_L_* and *h_L,CaMKII_* are the rates of Na_v_1.5 channel inactivation in the non-phosphorylated and phosphorylated forms, respectively, and *E_Na_* is the sodium reversal potential.

#### Calculation of APD and *d*(Δ*ψ_m_*(*t*))/*dt* at AP phase 3

The start time of each AP (phase 0) was determined by parsing the voltage at three successive time points (t_i_, t_i+1_, t_i+2_), such that V(t_i+2_) > V(t_i+i_) > V(t_i_) and V(t_i+2_) > 2 mV. The end time of each AP (phase 4) was determined by parsing the voltage at three successive time points, such that V(t_i+2_) < V(t_i+1_) < V(t_i_) and V(t_i+2_) < 2 mV. The duration of each AP was calculated by subtracting the start time from the end time.

*d*(Δ*ψ_m_*(*t*))/*dt*) at phase 3 (the late repolarization phase [16]) of each AP was calculated by fitting a line to five successive V(t) points centered on −30 mV using the polyfit function of MATLAB (which returns the slope and intercept). We then extended this line to two additional points at V(t_i-3_) and V(t_i+3_) using the polyfit function, which was overlaid on the corresponding AP using the plot function of MATLAB.

#### Calculation of the area under the curve (AUC) for 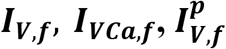 and 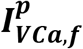

The area under the curve (AUC) was used as a metric of the current-time magnitudes conducted by the four fast VDI and CDI states relative to control. AUCs were calculated for each current using the trapz function of MATLAB (skipping the first 150 time points of each respective AP to isolate the late from the peak contribution).

## Results

Although anomalous I_Ca,L_ is the known direct cause of atypical depolarizations in CA, the specific mechanisms by which this current is generated are poorly understood. Our previous explanation is clearly oversimplified and incomplete [12], as are those of others [4]. In this work, we demonstrate that increasing disruption of the native dynamic inward-outward current balance leads to progressive AP alterations across three major perturbation thresholds (denoted as T_1_, T_2_, and T_3_). The ion current anomalies driving such alterations differ between hERG blockade/dysfunction and gain of late Na_v_1.5 function, and at short versus long pacing CLs. AP alterations progress in a graded fashion as a function of perturbation severity, entering the chaotic oscillatory regime at T_3_ (the severity of which is likewise graded). We attempted to de-convolute the effects of hERG and late Na_v_1.5 perturbations on the AP and underlying ion currents, as follows:

1. Ca_v_1.2 channels close in the wake of inactivation gate resetting/opening in preparation for the subsequent AP, resulting in <u>transient recovery</u> of inactivated sub-populations.
2. Perturbations promoting delayed Ca_v_1.2 closing lead to anomalous retention of the fast-VDI and CDI states, recovery from which evokes an anomalous I_Ca,L(late)_ window current at the T_1_ tipping point (comprised principally of *I_VCa,f_* and *I_V,f_*).
3. Atypical I_Ca,L_ spikes (~2-fold larger than control) arise when *I_VCa,f_, I_V,f_* and subspace Ca^2+^ grow to the tipping point at which CDI and VDI states recover.
4. Decreased slope of the AP (i.e. *d*(Δ*ψ_m_*(*t*))/*dt*) during phase 3 (centered at ~ —30 mV) under perturbed conditions results in slowed ion channel gating, including slowed opening of the fast CDI gate, slowed I_K1_ activation, and prolonged APD.
5. A CL- and perturbation-dependent “race” between spontaneous (non-paced) atypical I_Ca,L-_ driven depolarization, collision with the subsequent paced AP, and I_K1_-driven repolarization ensues.
6. Collisions depend on APD relative to CL (i.e. APD/CL), whereas spontaneous depolarizations depend on *d*(Δ*ψ_m_*(*t*))/*dt* and non-uniformity thereof across the AP wave train.
7. Increased minimum resting Δ*ψ_m_*(*t*), which is mirrored by impaired Kir2.1 activation and I_K1_ knockdown during atypical I_Ca,L_-driven depolarizations.
8. Knockdown of I_Na(early)_, I_Na(late)_, and I_to_ at the takeoff potential, which mirror the level of I_K1_ knockdown.

### I_Ca,L_ under control conditions (APD/CL << 1)

As is well appreciated, I_Ca,L_ plays a prominant role in arrhythmogenesis. I_Ca,L_ and other cardiac ion currents can be partitioned into early and late components coinciding with the depolarization and late repolarization phases (0 and 3 [16]) of the AP, respectively. I_Ca,L(early)_ is triggered by d gate opening in response to paced Ca_v_1.2 activation (Figure 4A), transitioning to I_Ca,L(late)_ as the channel population distributes among the active and fast and slow phosphorylated and non-phosphorylated VDI and CDI states (Figure 4B). CDI inactivation (which is likewise voltage-gated) is stimulated by elevated subspace Ca^2+^ during AP phases 0 to 2. All CDI and VDI inactivation gates (fcaf/fcas and ff/fs, respectively) open prior to depolarization, close at gate-specific rates during repolarization, and then reopen prior to channel closing in preparation for the subsequent AP. Small window currents are generated during inactivation gate re-opening prior to channel closing, the largest of which consist of *I_VCa,f_* and *I_V,s_* (Figure 4B). *I_VCa,f_* grows during AP prolongation, and therefore constitutes an Achilles’ Heel of the Ca_v_1.2 channel gating machinery (noting that *I_V,s_* remains similar to control under pro-arrhythmic conditions).

**Figure 4.**
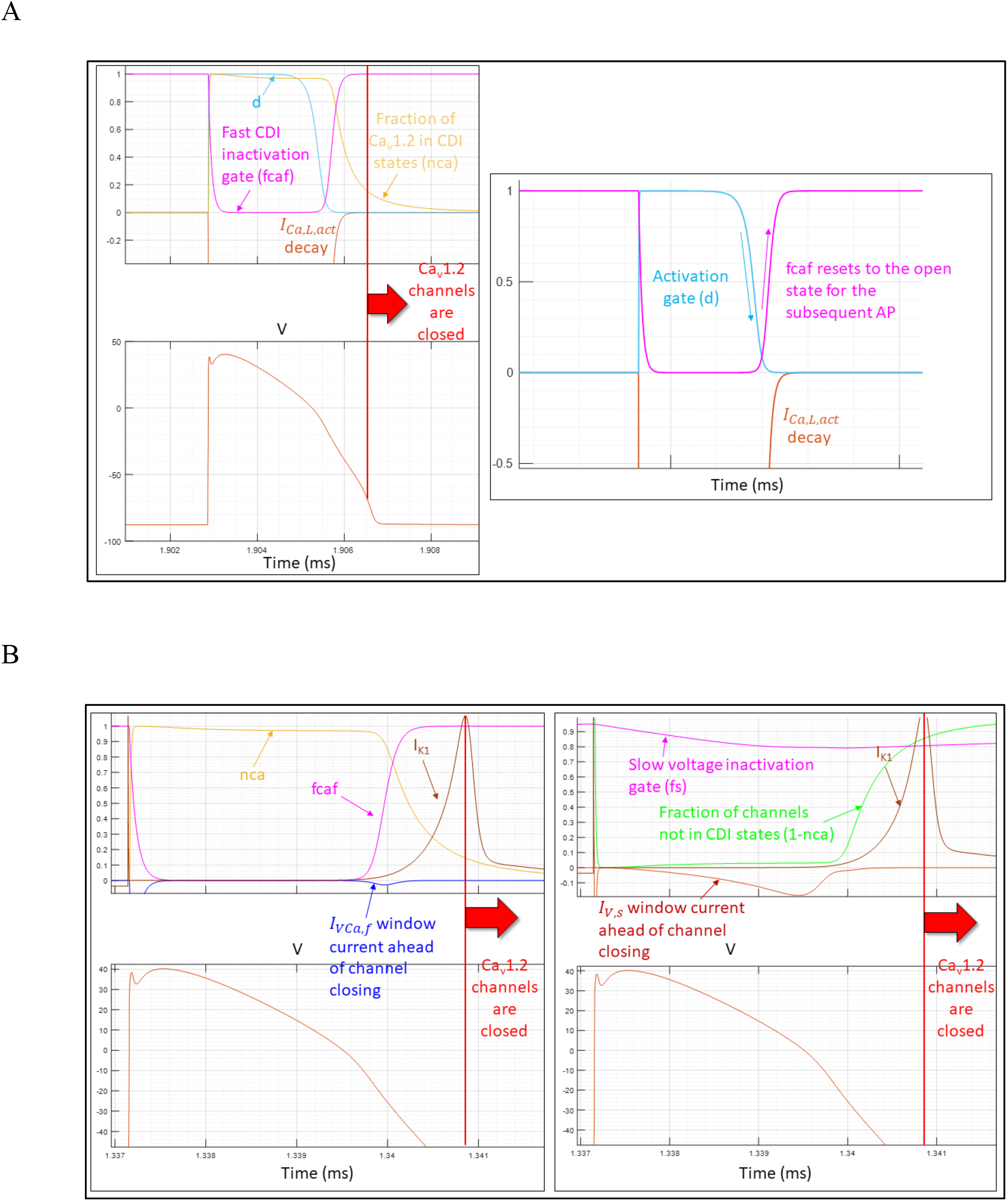
(A) The Ca_v_1.2 activation gate (denoted as d) opens in response to a paced depolarization, and closes during AP phase 3. Left panel: Registration between the AP (lower plot, red tracing), d gate (upper plot, cyan), and *I_Ca,L, act_* (upper plot, auburn tracing) simulated at a CL = 1/35 min. The d gate (light blue tracing) opens during depolarization (noting that the downward leg of d in the model consists of decreasing activation, rather than closing per se). The channels inactivate during repolarization via the slow (fcas) and fast (fcaf) CDI gates, and the slow (fs) and fast nonphosphorylated (ff) and phosphoryated (ffp) VDI gates, giving rise to 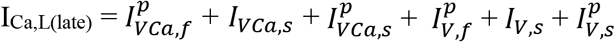. The fraction of channels in the CDI states (ncas) (gold tracing) is shown, togther with the d and fcaf gates (cyan and magenta tracings, respectively). Right panel: The inactivation gates re-open (reset) in preparation for the subsequent AP prior to closing (shown for fcaf). (B) Left panel: I_Ca,L_ window current generated by slight recovery into *I_VCa,f_* (via the fcaf gate). Right panel: I_Ca,L_ window current generated by slight recovery into *I_V,s_* (via the fs gate).

### The major stages of hERG dysfunction-induced arrhythmogenesis

The putative stages of arrhythmogenesis caused by hERG blockade and gain of inactivation/loss of recovery function consist of:

1. **Pro-arrhythmic/sub-acute** (< T_1_): Prolongation of paced APs in the Δ*ψ_m_*(*t*) ≥ –50 mV range, resulting in abnormal buildup of I_Ca,L_ toward the depolarization threshold (as outlined above). The resting voltage always extends into the Kir2.1 activation window.
2. **Pre-T_1_** (PT_1_): The highly unstable regime immediately prior to the depolarization threshold, which serves as a common reference for comparing pro-arrhythmic propensity (noting that this regime exists within a tiny perturbation range that is unlikely encountered in the clinical setting).
3. **Pre-arrhythmic/sub-acute** (≥ T_1_): At long CL (i.e. APD/CL < 1), I_Ca,L_ builds to the spontaneous (non-paced) depolarizing level prior to collisions between successive APs (denoted as AP_i_ and AP_j_). At short CL (i.e. APD/CL → 1), I_Ca,L_ builds to the depolarizing level during prolonged APi, which then “collides” with the subsequent <u>paced</u> AP (Figures 5 and 6A) to form AP_j_. The resting voltage always extends into the respective windows of hERG, KCNQ1, and Kir2.1 activation.
4. **Pre-arrhythmic/severe sub-acute** (≥ T_2_). AP_j_ collides with the subsequent paced AP (denoted as AP_k_), resulting in bifurcation of AP_j_ into AP_j_’ and AP_j_” (Figures 5 and 6A). The resting voltage always extends into the respective windows of hERG, KCNQ1, and Kir2.1 activation (Figure 6B).
5. **Arrhythmic/acute** (≥ T_3_): APs transition to non-paced oscillatory cycles (Figure 6A). The resting voltage always extends into the respective windows of hERG, KCNQ1, and Kir2.1 activation (Figure 6B).
6. **Highly arrhythmic/severe acute** (>> T_3_): Non-paced oscillatory cycles are either extinguished infrequently or not at all (Figure 6B). The minimum repolarized voltage rarely (if ever) extends into the Kir2.1 activation window.

**Figure 5.**
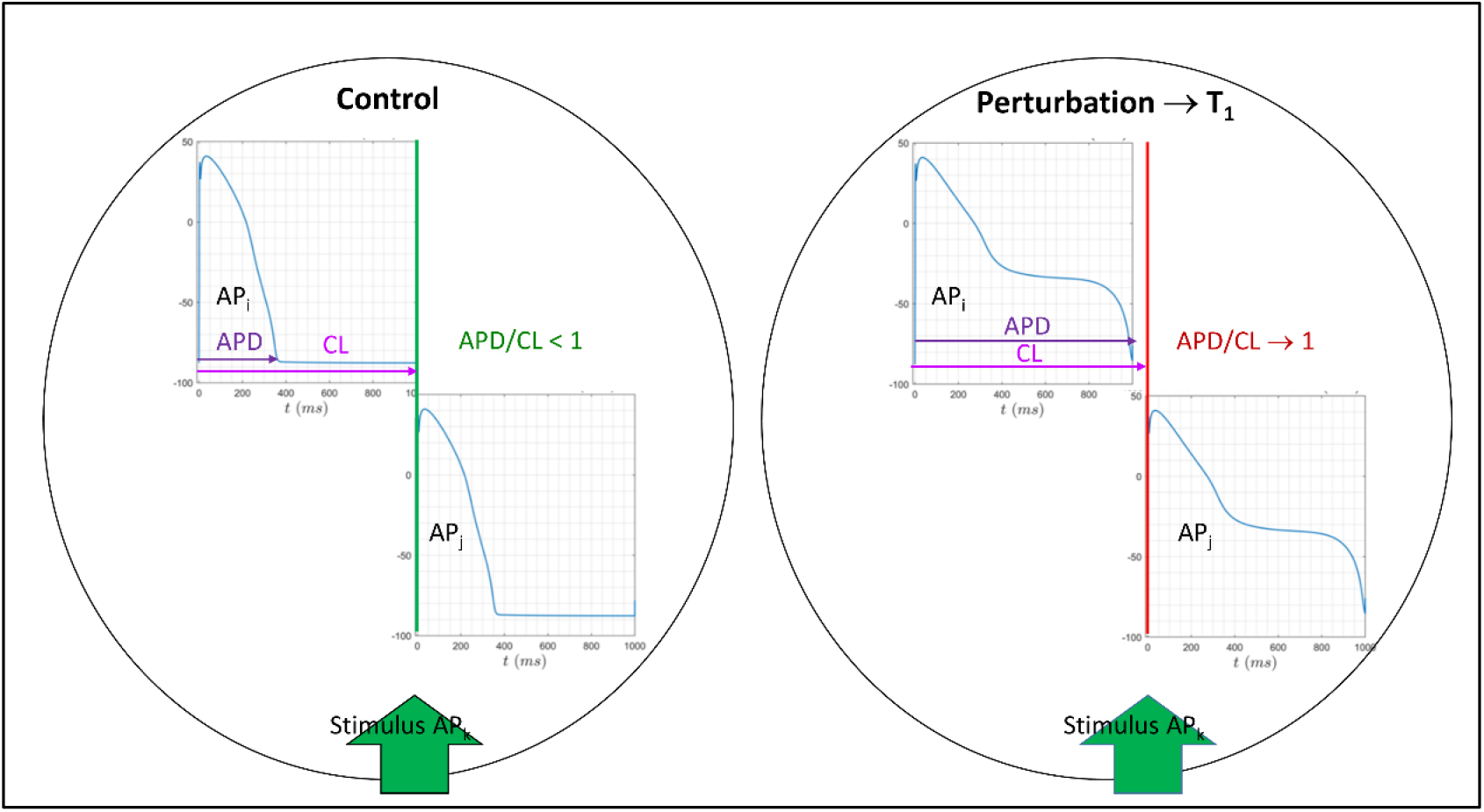
The effect of AP prolongation on the loss of restitution time, denoted in this work as 1 – APD/CL) between two successive APs (AP_i_ and AP_j_). AP_i_ and AP_j_ “collide” when APD/CL = 1, which triggers anomalous (though paced) I_Ca,L_-driven depolarization of AP_j_.

**Figure 6.**
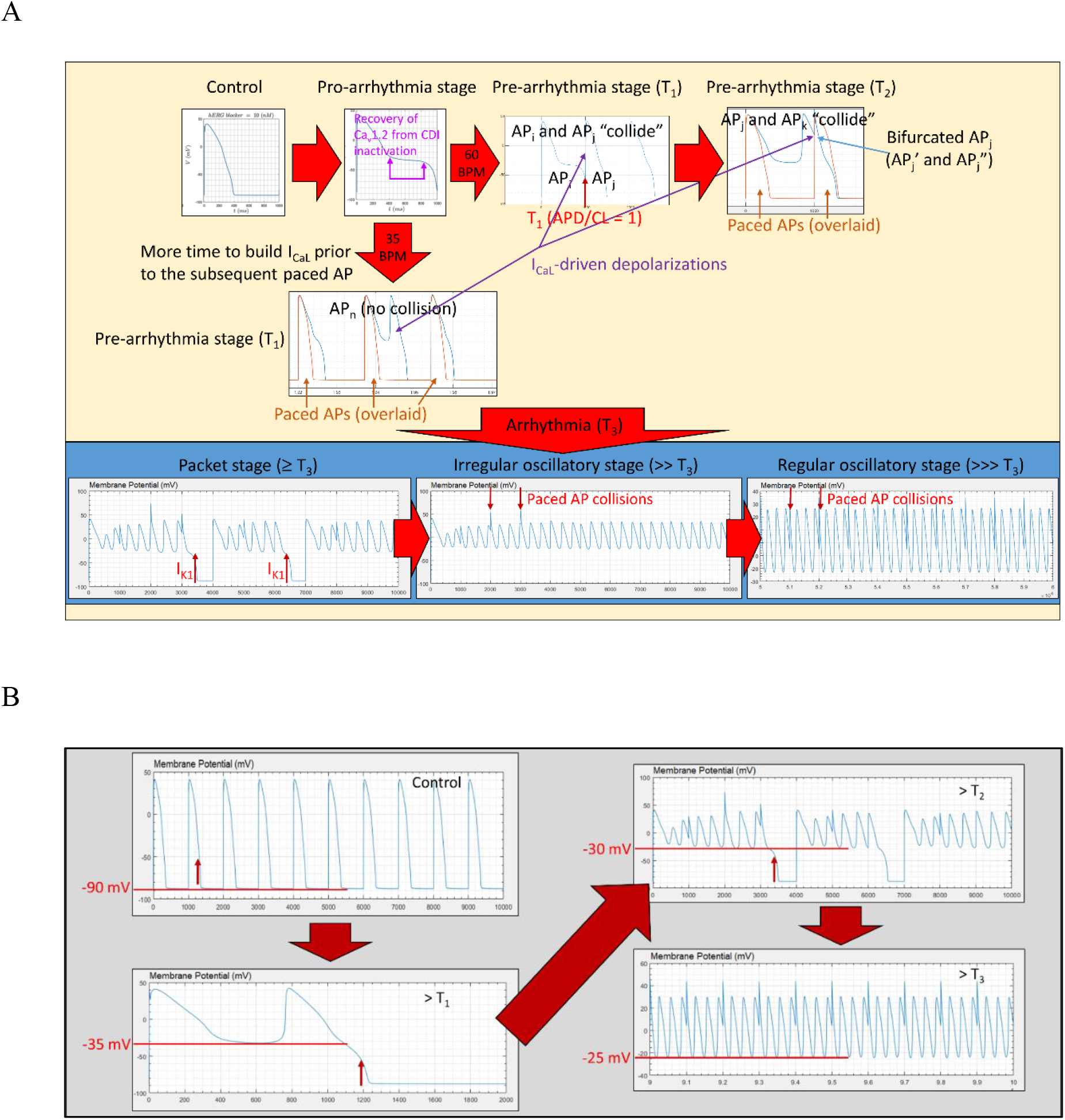
(A) The putative stages of arrhythmogenesis driven by hERG blockade or gain of hERG inactivation/loss of recovery function. Progression within and between each stage is graded and non-linear with respect to increasing levels of hERG blockade/dysfunction. Progressively greater APD lengthening occurs at ever finer increments of increasing blocker concentration on approach to T_1_. The prolonged AP-generation system is primed for anomalous AP_j_-forming depolarization at the T_1_ threshold, when: 1) APD/CL → 1 (resulting in collisions between successive APs); or 2) APD/CL < 1 and the slope of *d*(Δ*ψ_m_*(*t*))/*dt* = 0 mV (typically at longer CL). AP_j_ is primed for additional anomalous depolarizations upon collision with a subsequent AP (denoted as AP_k_), forming AP_j_’ and AP_j_”. (B) Kir2.1 activation follows the graded increase in Δ*ψ_m_*(*t*), which ranges between ~ –90 mV (control) and ~ –25 mV.

The AP-generation system responds to increasing levels of perturbation, as follows:

1. Slow quasi-linear growth in APD, accompanied by decreasing *d*(Δ*ψ_m_*(*t*))/*dt* within the shoulder region. T_1_ is approached more rapidly for trappable versus non-trappable blockers exhibiting slower k_on_, consistent with our earlier argument against a one-size-fits-all safety margin [12] (further addressed in [18]).
2. Large swings in successive APD (Figure 7) (short-long-short QT sequences are known to precede TdP). Such behavior, reflecting high instability of the system, is attributable to oscillations in *d*(Δ*ψ_m_*(*t*))/*dt*, APD, and ion channel activity on approach to the T_1_ tipping point.
3. Abrupt crossing of the T_1_ tipping point.

**Figure 7.**
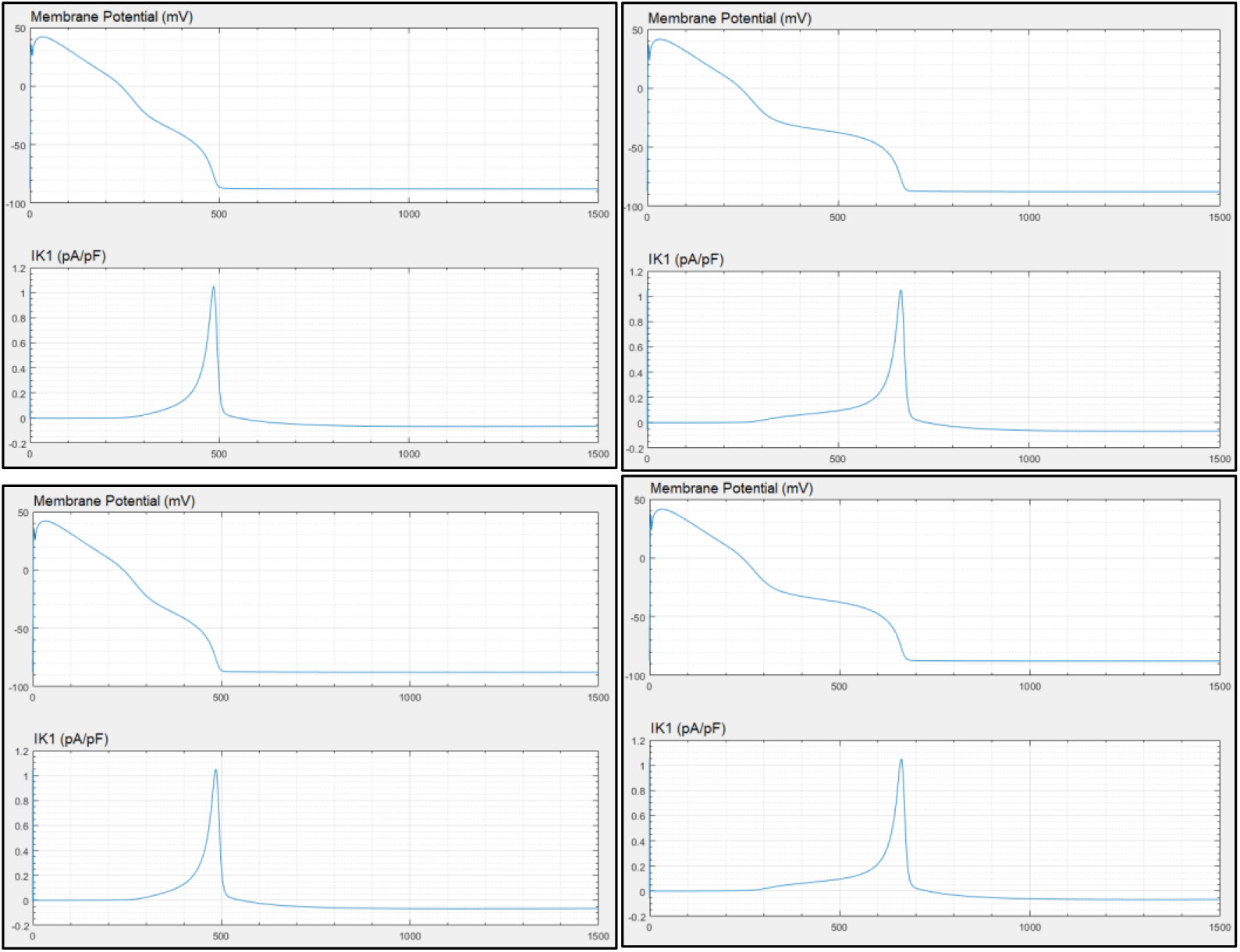
Short-long-short-long durations of four successive APs (top left to right, bottom left to right) caused by hERG blockade at the edge of the T_1_ tipping point (blocker attributes shown in bold in Table 1) at CL = 1/80 min, resulting from fluctuations in *d*(Δ*ψ_m_*(*t*))/*dt* at the edge of the Kir2.1 activation window. Such behavior reflects high instability of the system.

We studied AP morphology as a function of hERG blocker concentration, titrated in increasingly fine increments toward the AP_i_-AP_j_ (APD/CL = 1) and AP_j_-AP_k_ collision thresholds (T_1_ and T_2_ = 51.95 and 51.9511 nM, respectively) (Figure 8).

**Figure 8.**
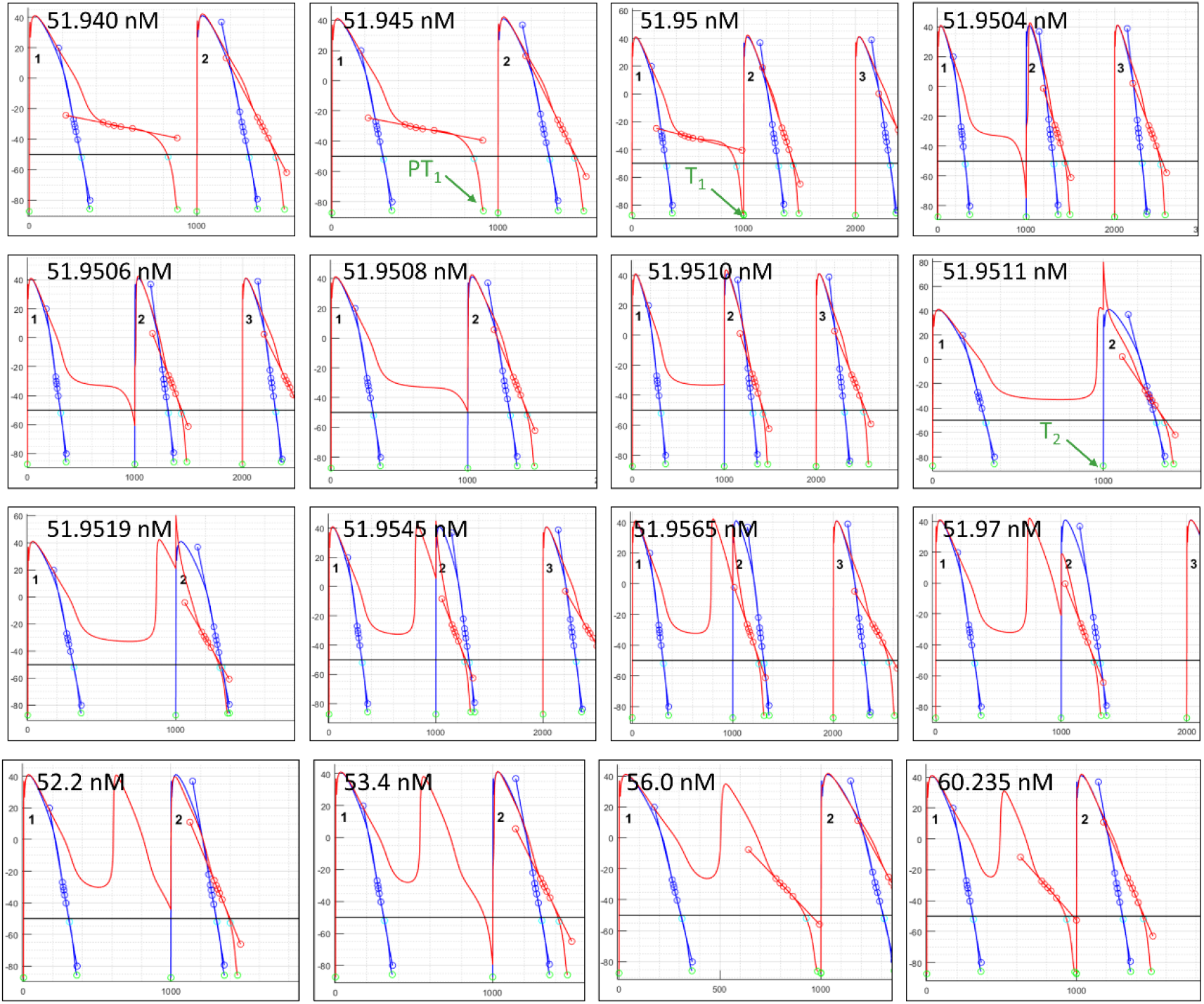
Snapshots of hERG blockade effects on APs (including *d*(Δ*ψ_m_*(*t*))/*dt* shown as tangent lines) at the T_1_ and T_2_ thresholds (simulated at CL = 1/60 min), demonstrating graded responses of the AP waveform to incremental changes in perturbation severity (hERG blocker concentrations as noted). Collisions between successive paced APs (AP_i_-AP_j_ and AP_j_-AP_k_) result in anomalous depolarizations.

Dynamically counter-balanced systems are subject to oscillations when Yin and Yang are coupled in reciprocal out-of-phase relationships (as in the AP generation system), in which inward currents activate outward currents, and vice versa through their opposing effects on Δ*ψ_m_*(*t*). The AP is terminated under control conditions when the current balance is tipped strongly in the hyperpolarized direction by I_K1_ activation (Figure 9A). However, the I_K1_ activation window (Δ*ψ_m_*(*t*) ≾ −50 mV) (Figure 9B) is inaccessible below a threshold of I_Kr_ + I_Ks_ (noting that I_K1_ knockdown in nodal cells and iPSC-derived cardiomyocytes results in sustained AP oscillations). Total knockout of I_Kr_ results in ringing oscillations in the AP waveform (Figure 10).

**Figure 9.**
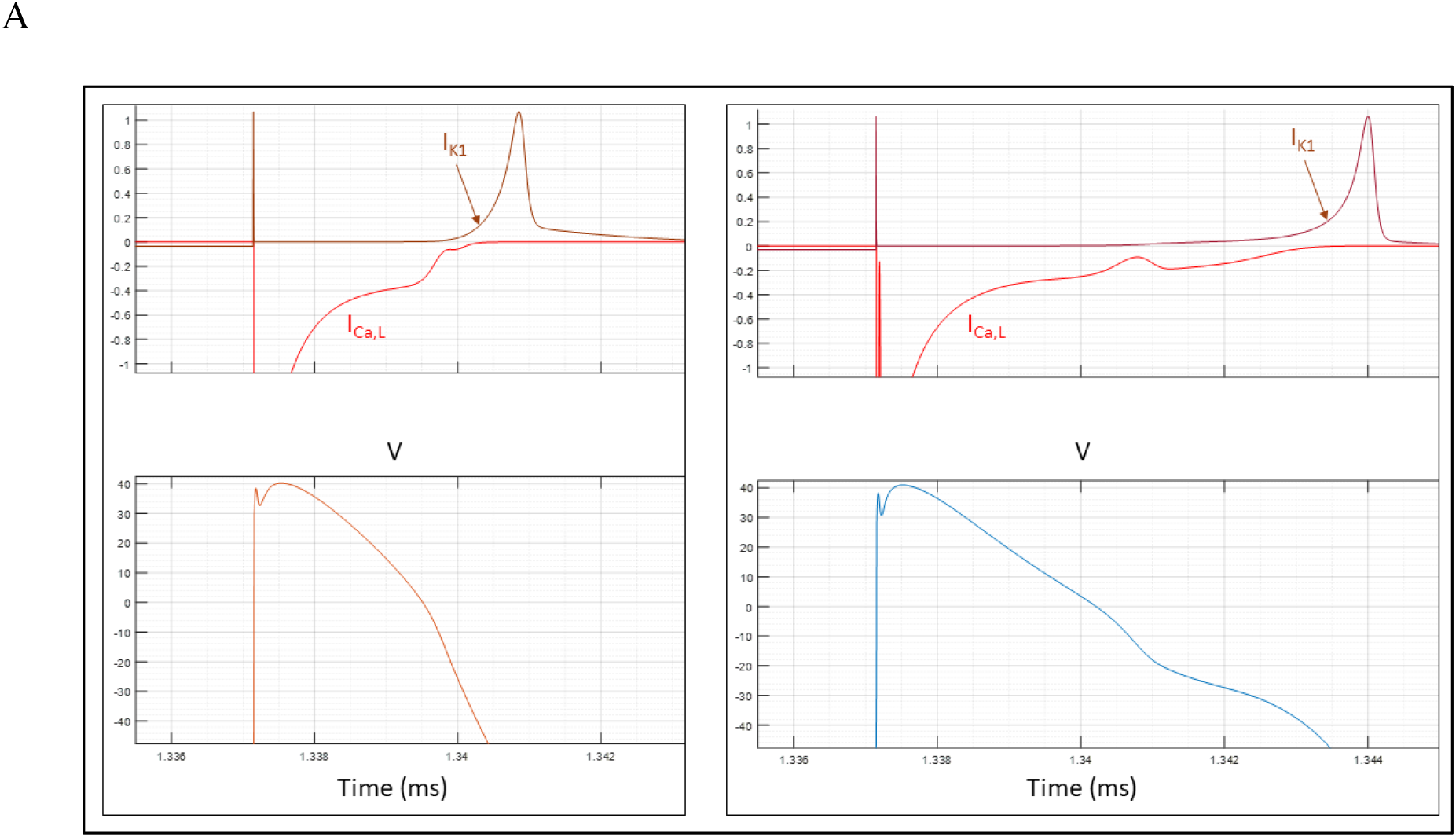
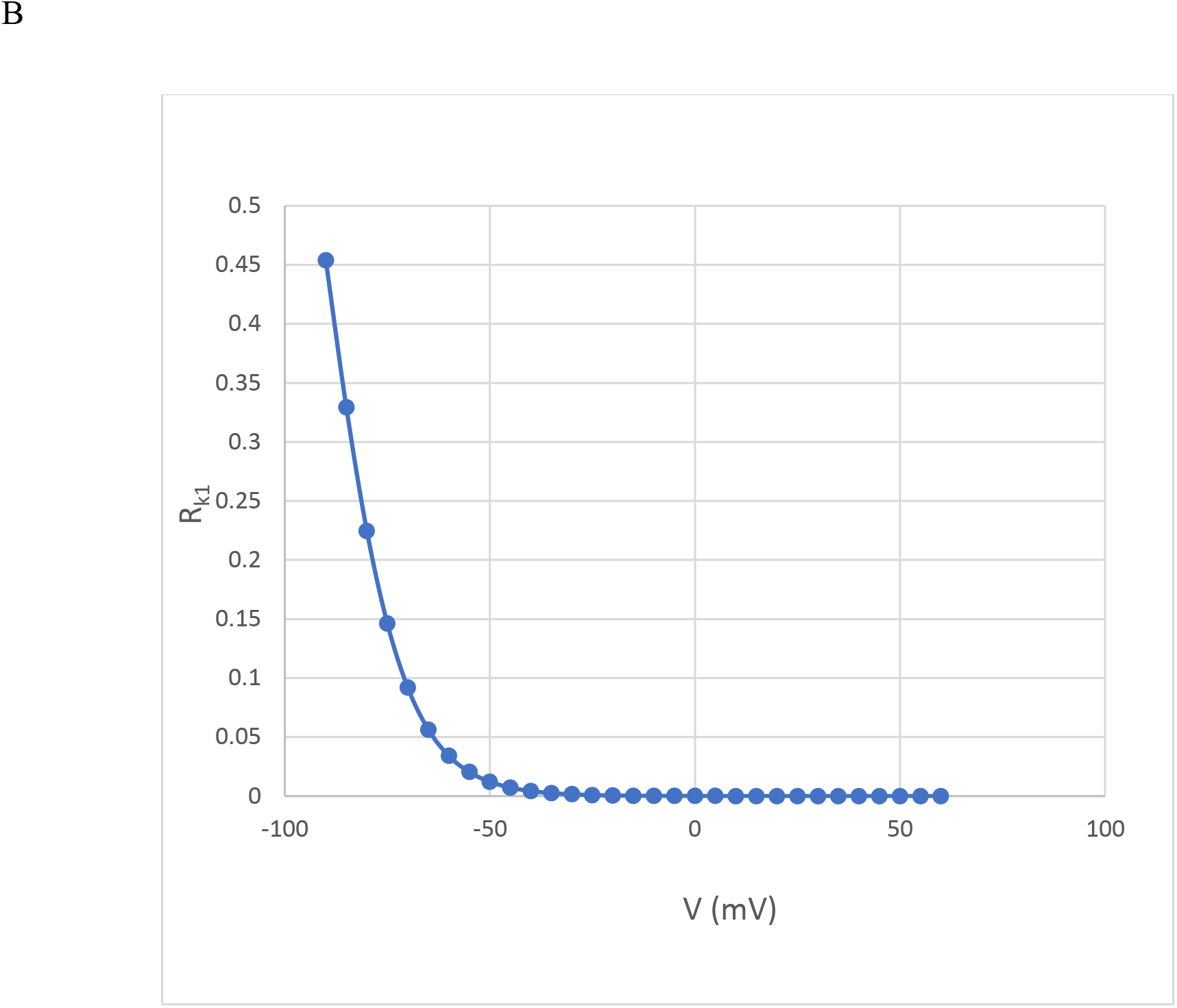
(A) Both control (left) and sub-T_1_ prolonged (right) APs are terminated by I_K1_ activation (signaling the end of open ion channel states). (B) *R*_*k*1_, the voltage-dependence of channel activation in the ORd and ORd/hERG Markov models, 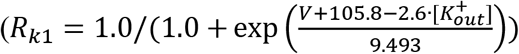, where V is the voltage and 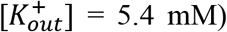. APD is thus directly determined by the rate of I_K1_ activation, which in turn, is governed by *d*(Δ*ψ_m_*(*t*))/*dt* during AP phase 3.

**Figure 10.**
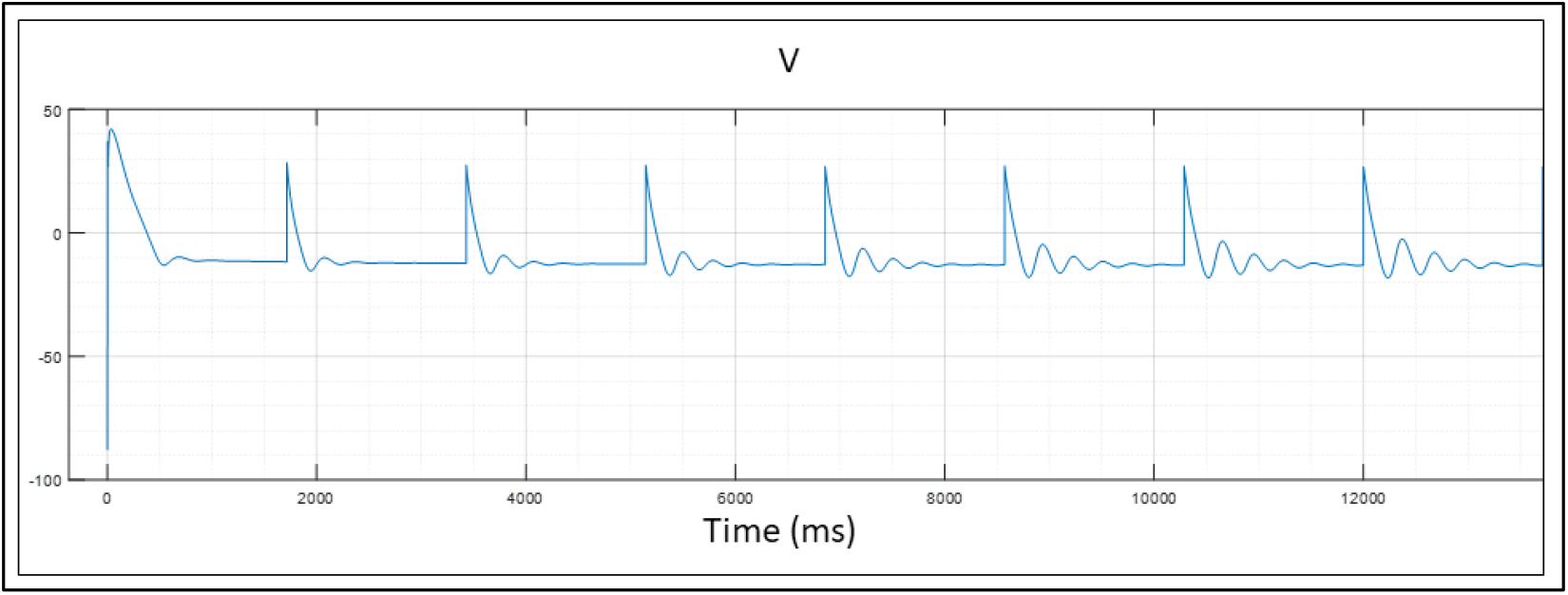
The Δ*ψ_m_*(*t*) waveform under conditions of total I_Kr_ knockout at CL = 1/35 min. Each arrhythmic cycle consists of a paced depolarization, followed by a series of ringing oscillations (noting that the number of oscillations between pacing signals decreases with decreasing CL). The underlying currents driving this process are discussed in a later section.

### The role of Ca^2+^ in arrhythmogenesis

As is widely appreciated, the myocardial contraction cycle is driven by the intracellular Ca^2+^ cycle, which in turn is driven by the AP cycle. The Ca^2+^ cycle consists of the following steps:

1. Release from the sarcoplasmic reticulum (SR) via RyR_2_ channels into the subspace.
2. Diffusion from the subspace to the myoplasm, resulting in transient troponin activation.
3. Re-uptake to the SR from the myoplasm via the SERCA pump (Figure 1) and reversal of troponin activation.

The driving force for cyclic inter-compartmental ion translocation consists of transmembrane Ca^2+^, Na^+^, and K^+^, trans-SR Ca^2+^, and subspace-intracellular Ca^2+^ gradients that are maintained away from their respective equilibrium distributions (as given by the Nernst equation), and undergo partial rundown during translocation (reset by ion exchange pumps). Ca^2+^ directly activates a host of molecular targets, including troponin, RyR_2_, and CaMKII (which phosphorylates ion channels, ion exchangers, RyR_2_, and the SERCA pump), and inactivates Ca_v_1.2 via Ca^2+^-dependent inactivation. Positive or negative attenuation of the aforementioned ion gradients is highly pro-arrhythmic. Elevated intracellular Ca^2+^ slows ion channel activation and speeds inactivation via enhanced CaMKII-mediated phosphorylation (an overall anti-arrhythmic effect). However, elevated subspace Ca^2+^ promotes the CDI states of Ca_v_1.2 (recovery from which drives atypical depolarizations in our simulations). Recovery from both the CDI and VDI states is further enhanced by increased 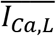 at depolarized Δ*ψ_m_*(*t*) resulting from elevated subspace Ca^2+^. 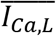 is calculated in the ORd and ORd/hERG Markov models as follows:

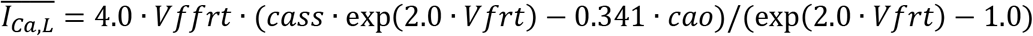

where 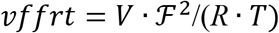 and 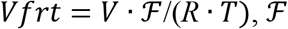 is the Faraday constant, *R* is the gas constant, and *V* is the voltage. Subspace Ca^2+^ is calculated in the ORd and ORd/hERG Markov models, as follows:

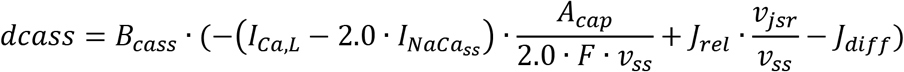

where *dcass* is the derivative of the free subspace Ca^2+^ concentration (solved via numerical integration in the model), *B_cass_* is the free fraction of subspace Ca^2+^, *I_NaCa_ss__* is the subspace Ca^2+^ efflux current, *A_cap_* is the capacitive area, *v_ss_* is the subspace volume, *V_jsr_* is the junctional SR volume, *J_rel_* is the rate of RyR_2_ release, and *J_diff_* is the rate of subspace to myoplasmic Ca^2+^ diffusion. The dynamic subspace Ca^2+^ level is thus coupled recursively to its inputs and outputs. Anomalous subspace Ca^2+^ elevation results in increased 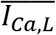 (translating to increased I_Ca,L_), which, in turn, drives increased Ca^2+^ release. Dynamic subspace and intracellular Ca^2+^, the activated CaMKII fraction, and the isolated activated *I_Ca,L, act_* fraction are plotted in Figure 11A, 11B, and 11C, respectively.

**Figure 11.**
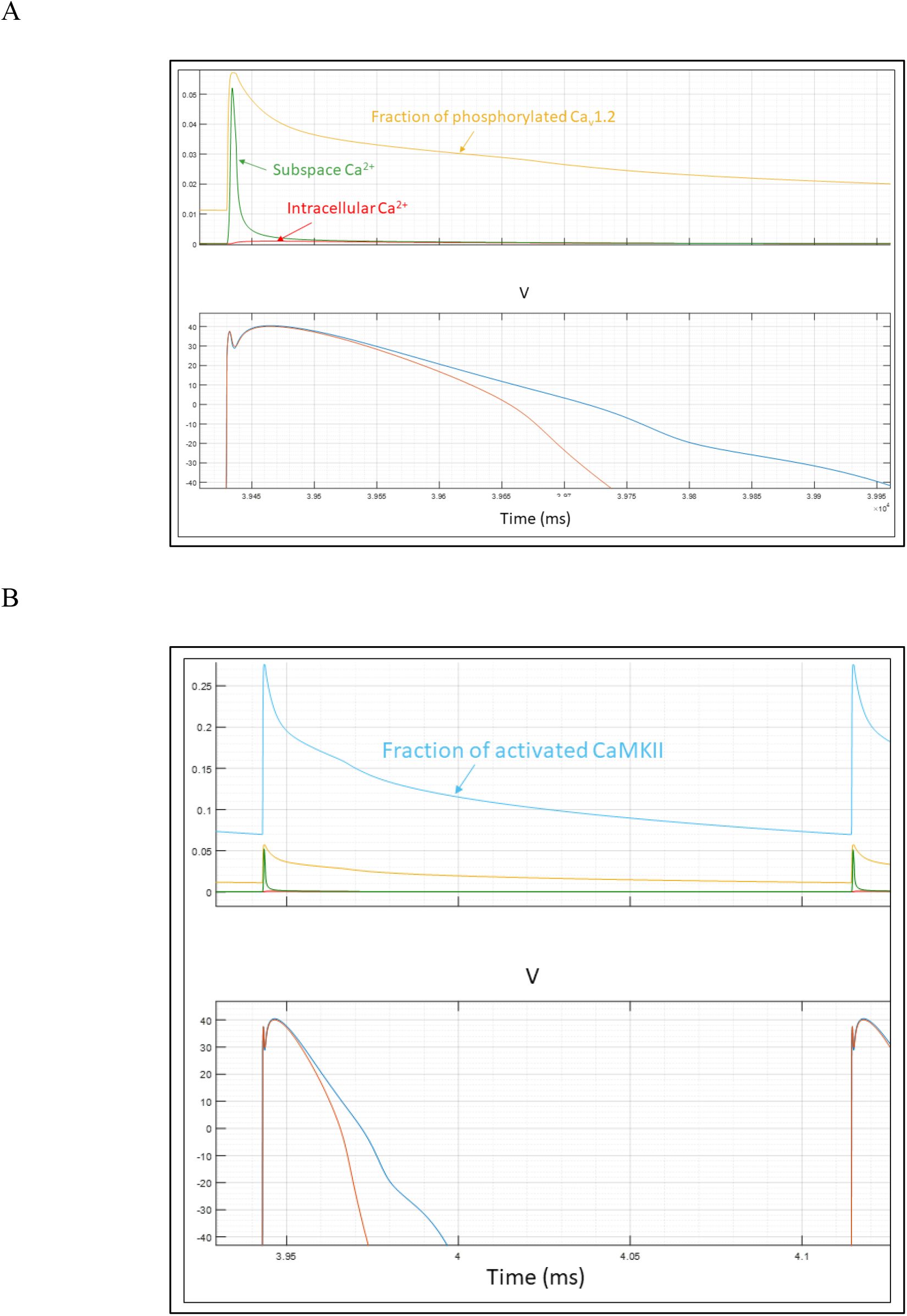
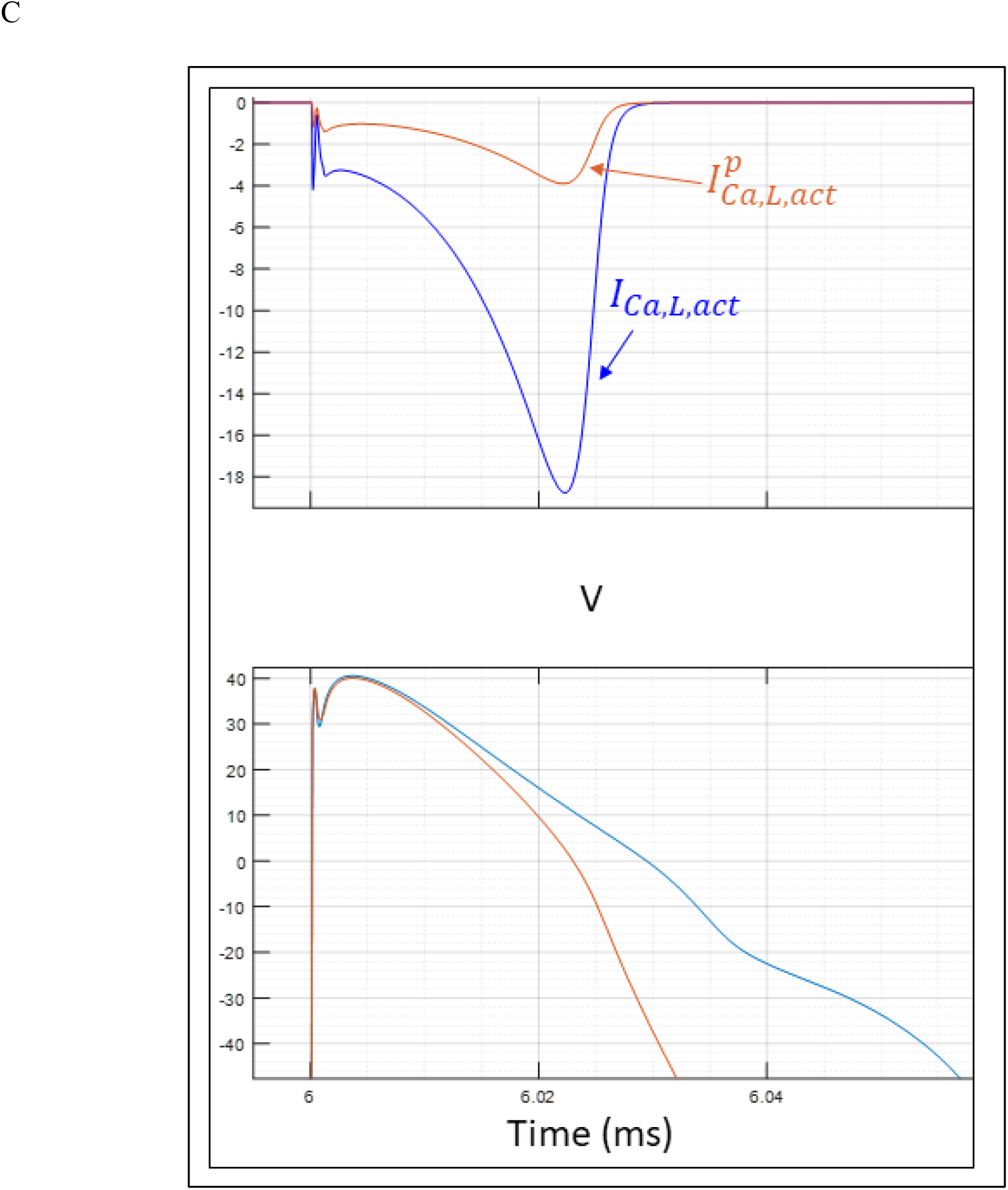
(A) Free subspace and intracellular Ca^2+^ concentrations (mM) (green and red tracings, respectively) and Ca^2+^-activated CaMKII (light blue tracing) under control conditions at CL = 1/35 min. Free subspace Ca^2+^ spikes during AP phase 0, diffuses to the myoplasm, and binds to various proteins, including troponin, calmodulin, and Ca_v_1.2. (B) Ca_v_1.2 is phosphorylated and dephosphorylated during each AP by activated CaMKII (yellow tracing). (C) Phosphorylation slows Ca_v_1.2 activation (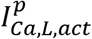 versus *I_Ca,L, act_*).

### The effects of hERG dysfunction on ion currents in the APD/CL < 1 regime (hERG blocker concentration < T_1_ @ CL = 1/35 min)

Next, we studied the ion current profiles during prolonged APs and spontaneous non-paced I_Ca,L_-driven depolarizations in the absence of AP collisions at CL = 1/35 min. We titrated the hERG blocker concentration to the approximate T_1_ threshold (~46.35 nM) at this CL, and simulated the APs under these and control conditions. The first ~100 APs exhibited prolonged, but relatively uniform *d*(Δ*ψ_m_*(*t*))/*dt* in the shoulder region, which became increasingly shallower and variable over the latter ~100 APs where atypical I_Ca,L_-driven depolarizations occurred.

#### I_Ca,L_ effects during the late pro- and early pre-arrhythmic stages

The late pro-arrhythmic stage (approaching T_1_) is characterized by anomalously large *I_VCa,f_* and *I_V,f_* window currents resulting from:

1. Increased dwelling of phosphorylated and non-phosphorylated Ca_v_1.2 in the CDI and VDI states prior to I_K1_-driven channel closing (Figure 12A). Increased dwelling in the CDI states is attributable to abnormal subspace Ca^2+^ levels (Figure 13A-E) and CaMKII activation (Figure 13F).
2. Enhanced recovery of Ca_v_1.2 channels from the CDI and VDI states prior to delayed channel closing. *I_VCa,f_* and *I_V,f_* broadening (corresponding to slowing of the fcaf and ff gates (Figure 12B)) and slowed I_K1_ activation (Figure 14A) are attributable to decreased *d*(Δ*ψ_m_*(*t*))/*dt* within the prolonged shoulder region.

**Figure 12.**
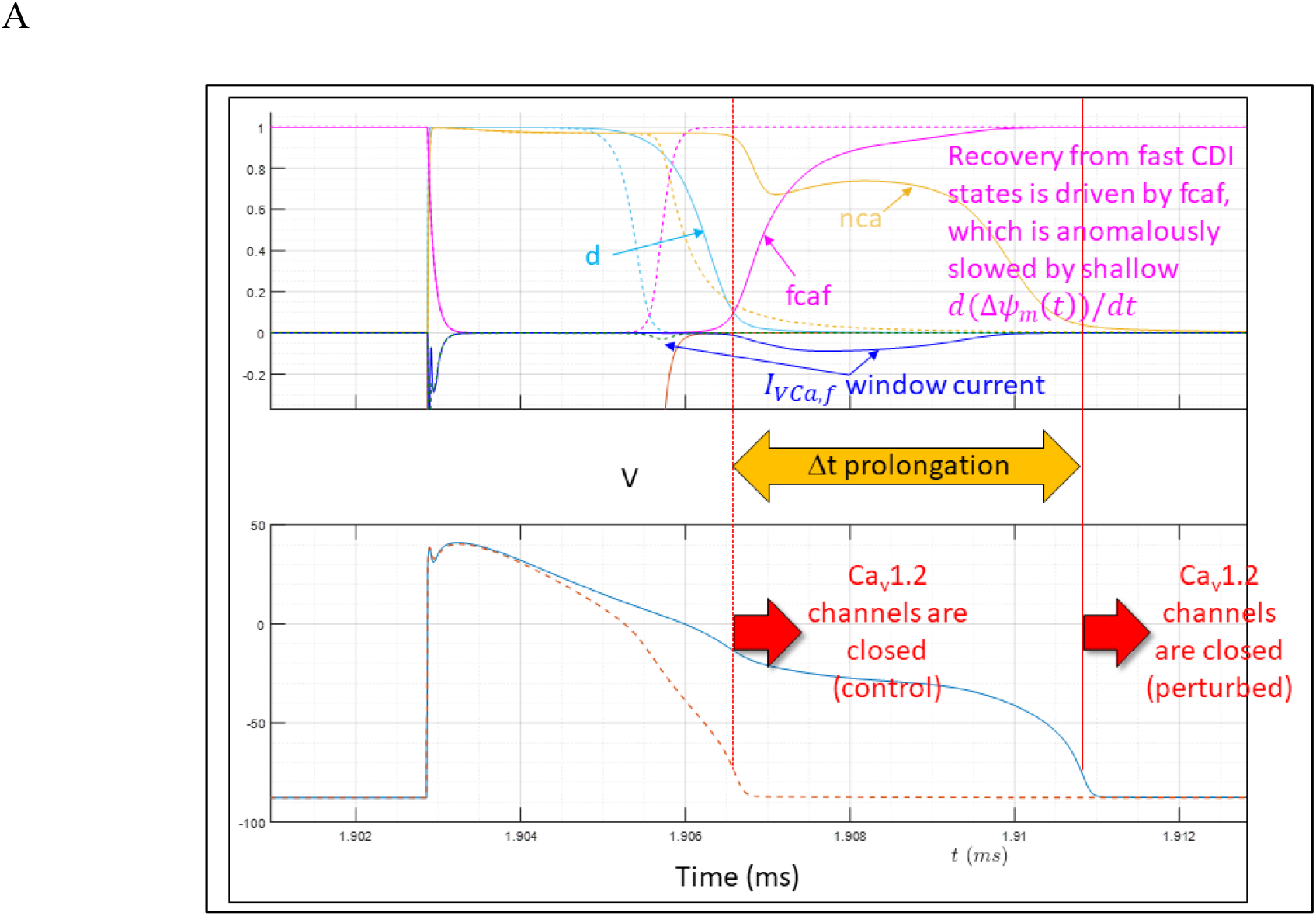
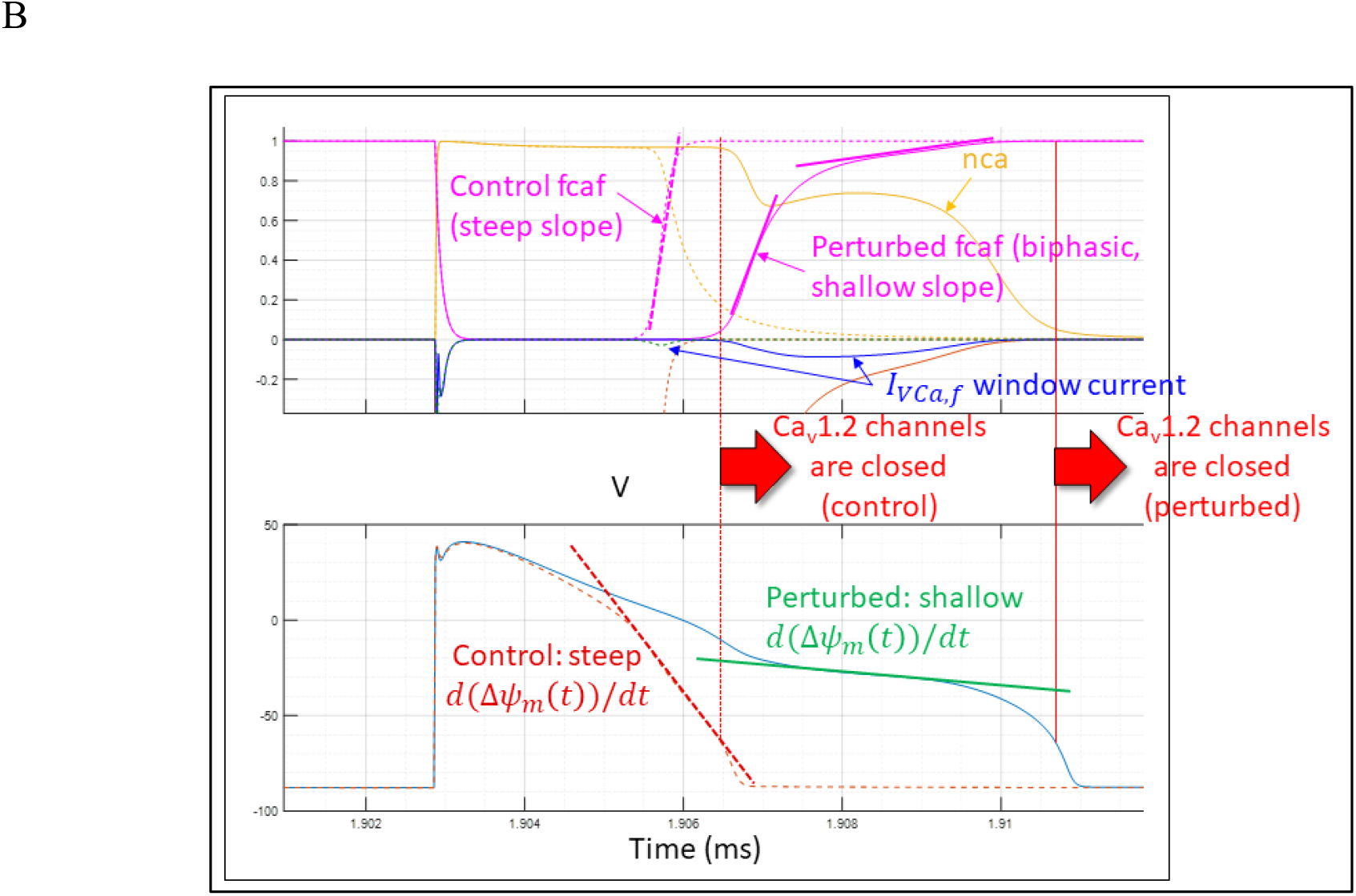
(A) Increased *I_VCa,f_* during prolonged APs that exhibit anomalously shallow *d*(Δ*ψ_m_*(*t*))/*dt*. The fcaf gate slows significantly under perturbed (solid magenta tracing) versus control (dotted magenta tracing) conditions, resulting in prolonged recovery of the fast CDI state (generating *I_VCa,f_*). The fraction of channels in the CDI states (nca) is abnormally large under perturbed (solid gold tracing) versus control (dotted gold tracing) conditions due to Ca^2+^ release from the SR during this time period. (B) Recovery of *I_VCa,f_* and *I_V,s_* results from Δ*ψ_m_*(*t*)-driven fcaf and fs opening. Broadening of these currents results from slowed fcaf and ff opening due to decreased *d*(Δ*ψ_m_*(*t*))/*dt* within the shoulder region.

**Figure 13.**
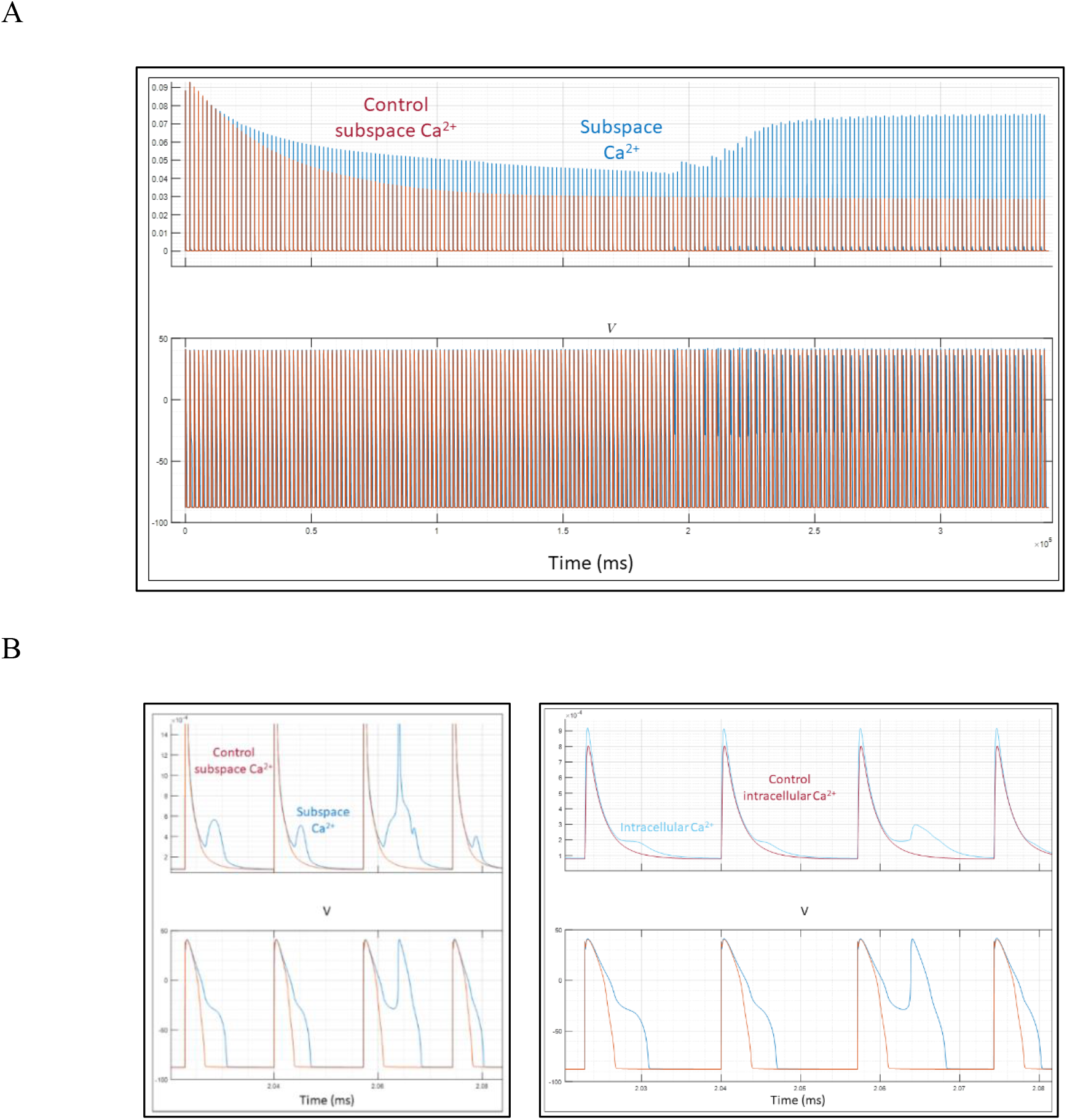
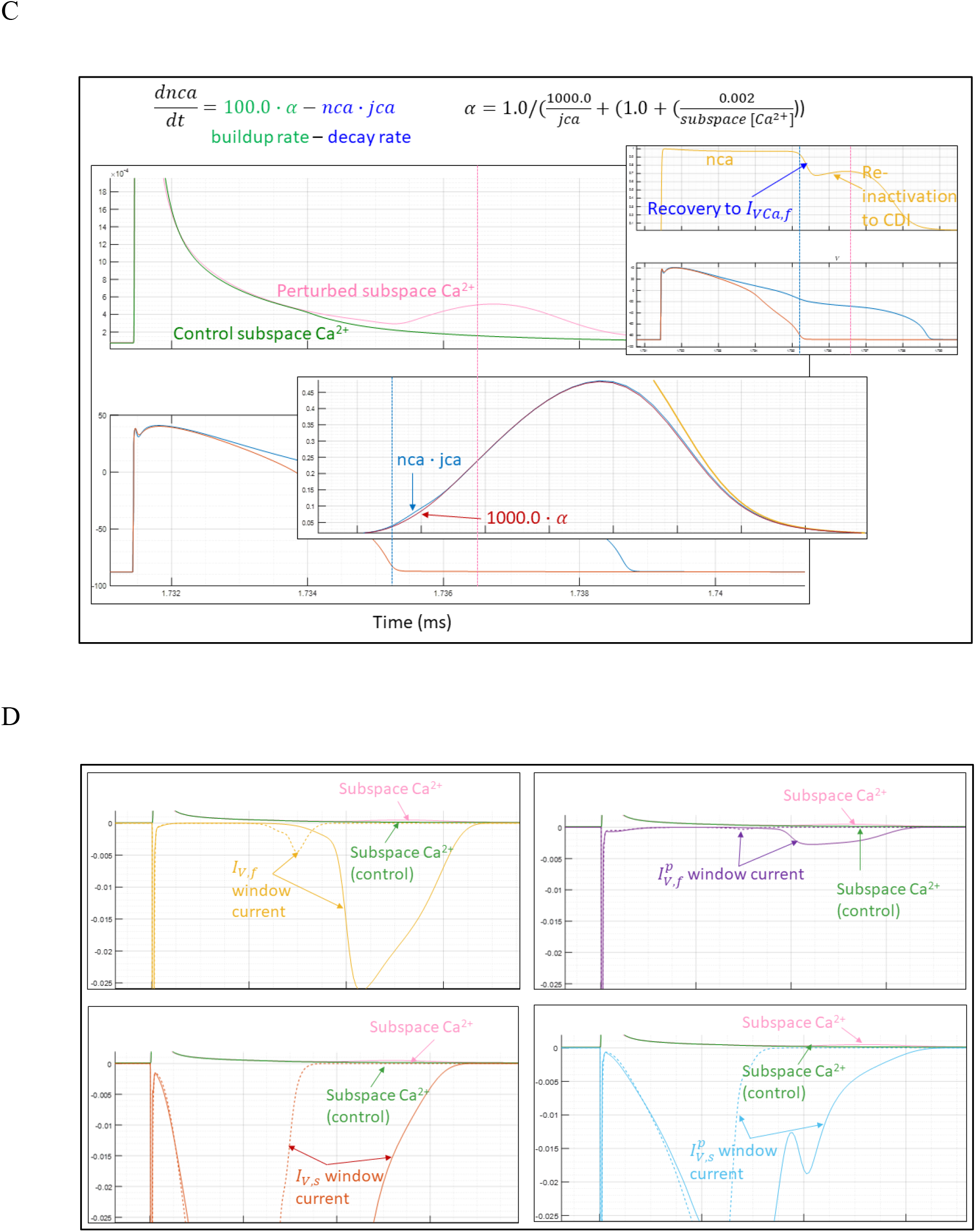
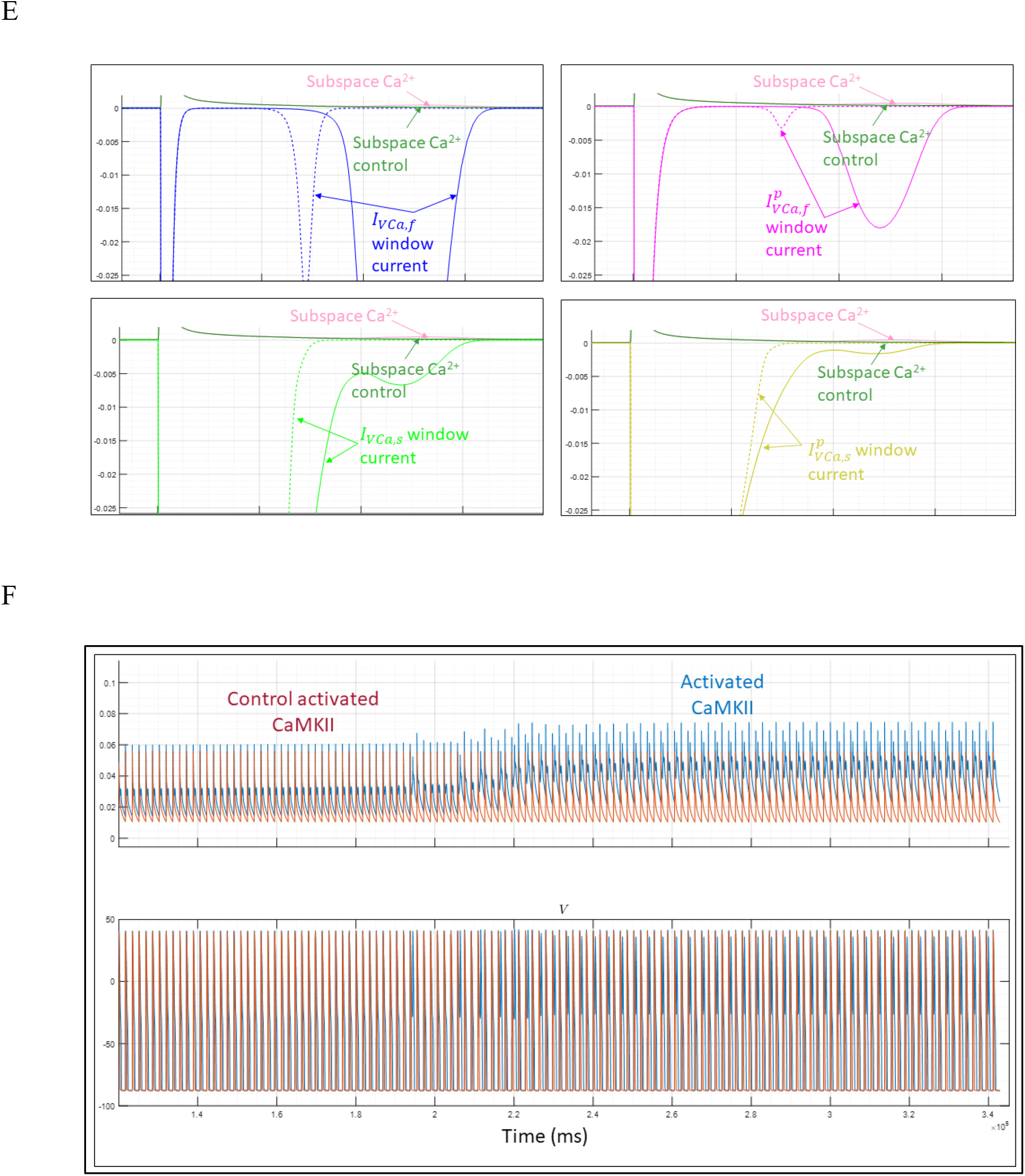
(A) Zoomed out view of subspace Ca^2+^ levels under control (auburn tracing) and perturbed (blue tracing) conditions throughout the 200 APs of the simulation. Elevated Ca^2+^ under perturbed conditions (primarily in the subspace) is due to increased Ca^2+^ cycling (RyR_2_-mediated SR release, translocation within the SR, and SERCA-mediated SR re-uptake). (B) Zoomed in view of time-dependent subspace (left) and intracellular (right) Ca^2+^ levels under control and perturbed conditions (orange and blue tracings, respectively). (C) Ca^2+^ release caused by the Ica,L window currents stimulates anomalous release of Ca^2+^ from the SR (coral tracing) relative to control (green tracing), which further promotes transitioning to the CDI states. nca(t) is determined by the rate of Ca^2+^-dependent buildup *(α)* versus recovery-dependent decay *(nca · jca)* to/from the CDI states (*d*(*nca*(*t*))/*dt*). (D) Co-dependence of *I_V,x_* and 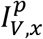 window current and subspace Ca^2+^ time-dependent behavior. (E) Co-dependence of *I_VCa,x_* and 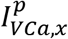 window current and subspace Ca^2+^ time-dependent behavior. (F) Zoomed out view of activated CaMKII levels under control and perturbed conditions throughout the 200 APs of the simulation (orange and blue tracings, respectively). Increased CaMKII activation results in enhanced ion channel, exchanger, RyR_2_, and SERCA phosphorylation.

*I_VCa,f_* and *I_V,f_* broadening grows in a graded fashion with increasing perturbation prior to T_1_. A tipping point in *I_VCa,f_* recovery ensues at T_1_, which promotes voltage-dependent recovery and re-inactivation of all CDI and VDI states via opening and subsequent closing of the fcaf, fcap, fcas, ff, ffp, and fs gates, respectively (manifesting as a spike in I_Ca,L(late)_) (Figures 14A and 14B). The window current magnitudes during recovery from the CDI and VDI states are limited under control conditions by short lag times between inactivation gate opening/resetting and channel closing. *I_VCa,f_* and *I_V,f_* grow with decreasing *d*(Δ*ψ_m_*(*t*))/*dt*, corresponding to slowing of the inactivation gates, together with I_K1_-mediated channel closing. Vulnerability of this mechanism to delayed closing constitutes a major Achilles’ Heel of the Ca_v_1.2 gating system.

**Figure 14.**
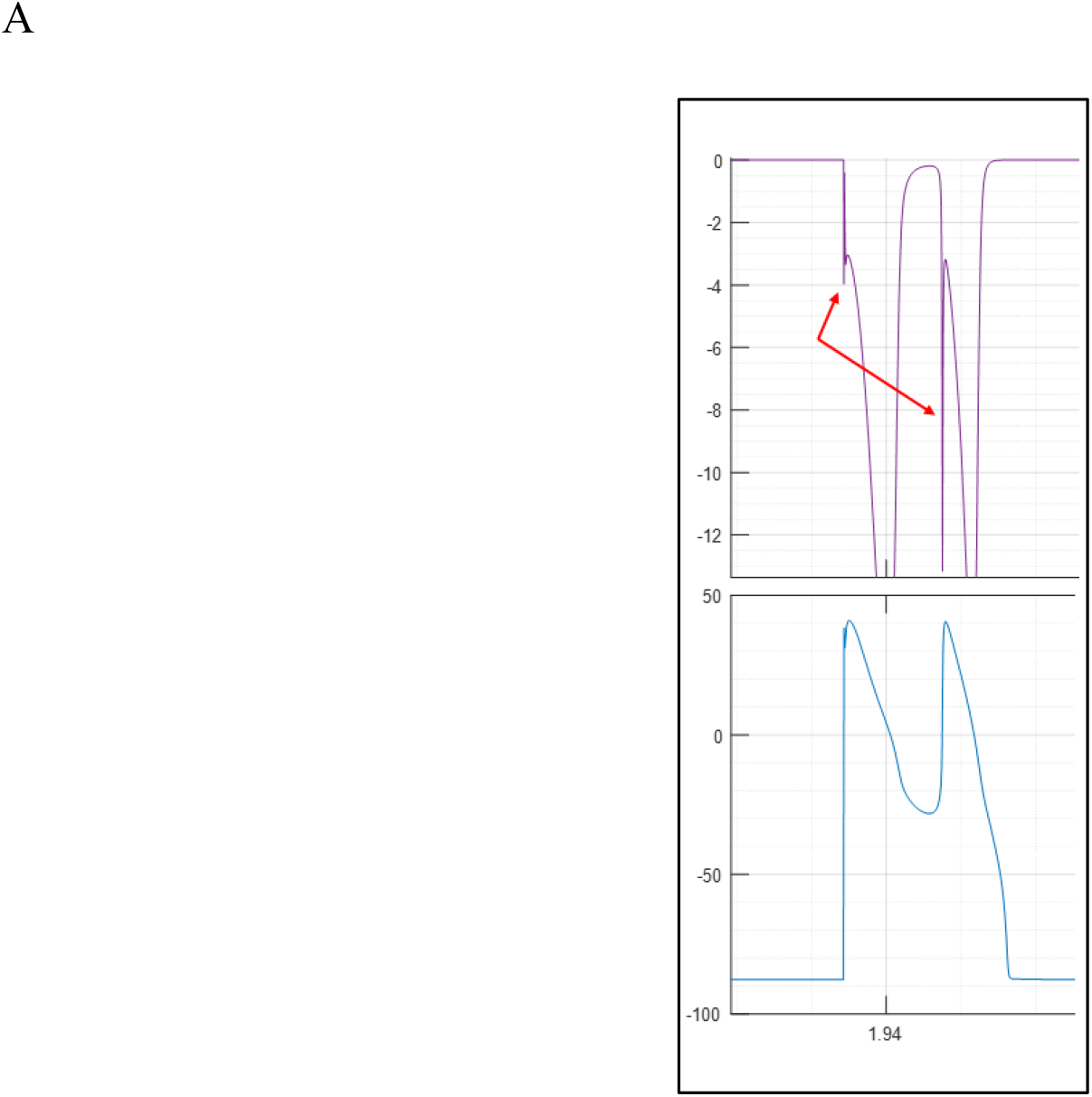
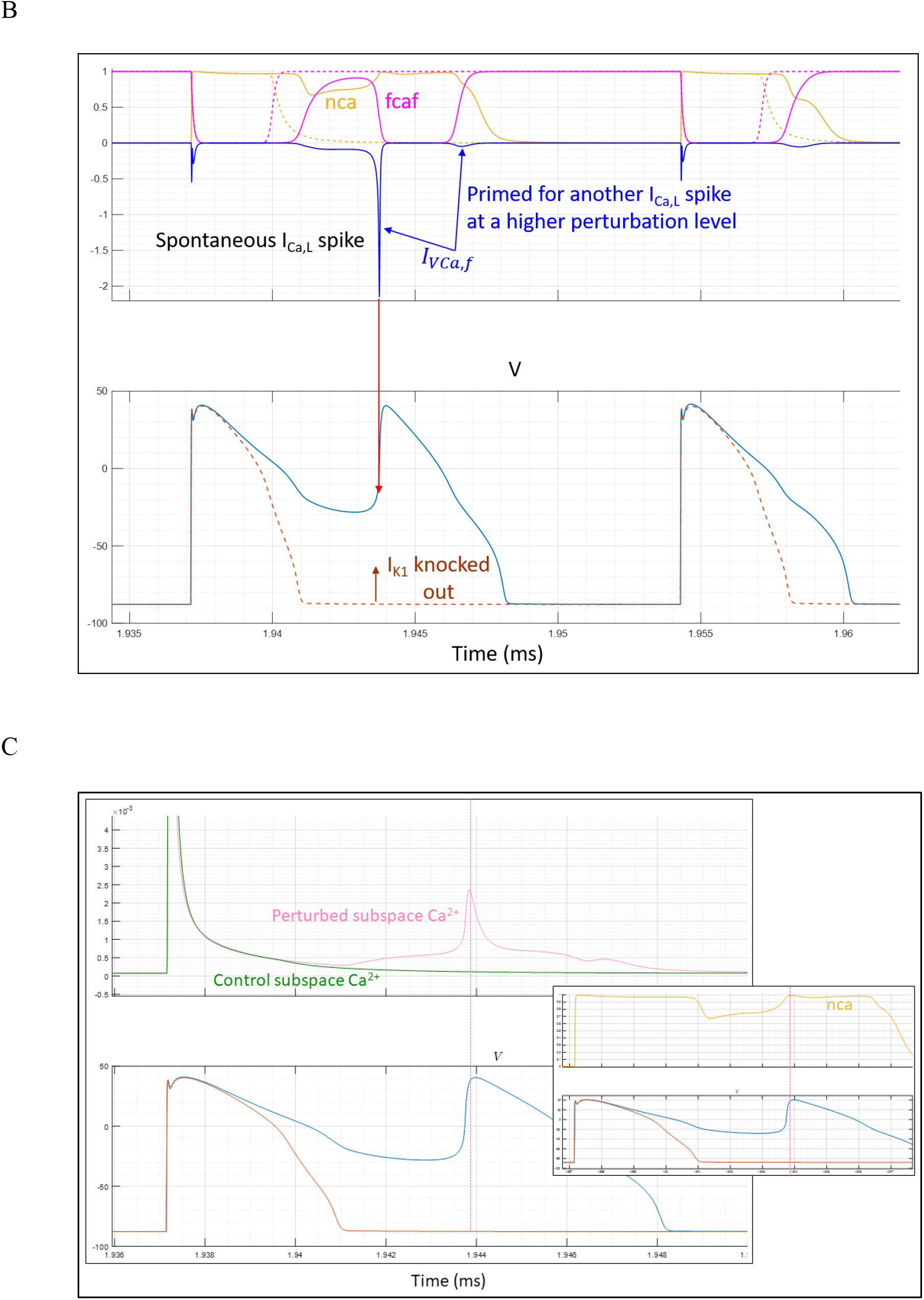
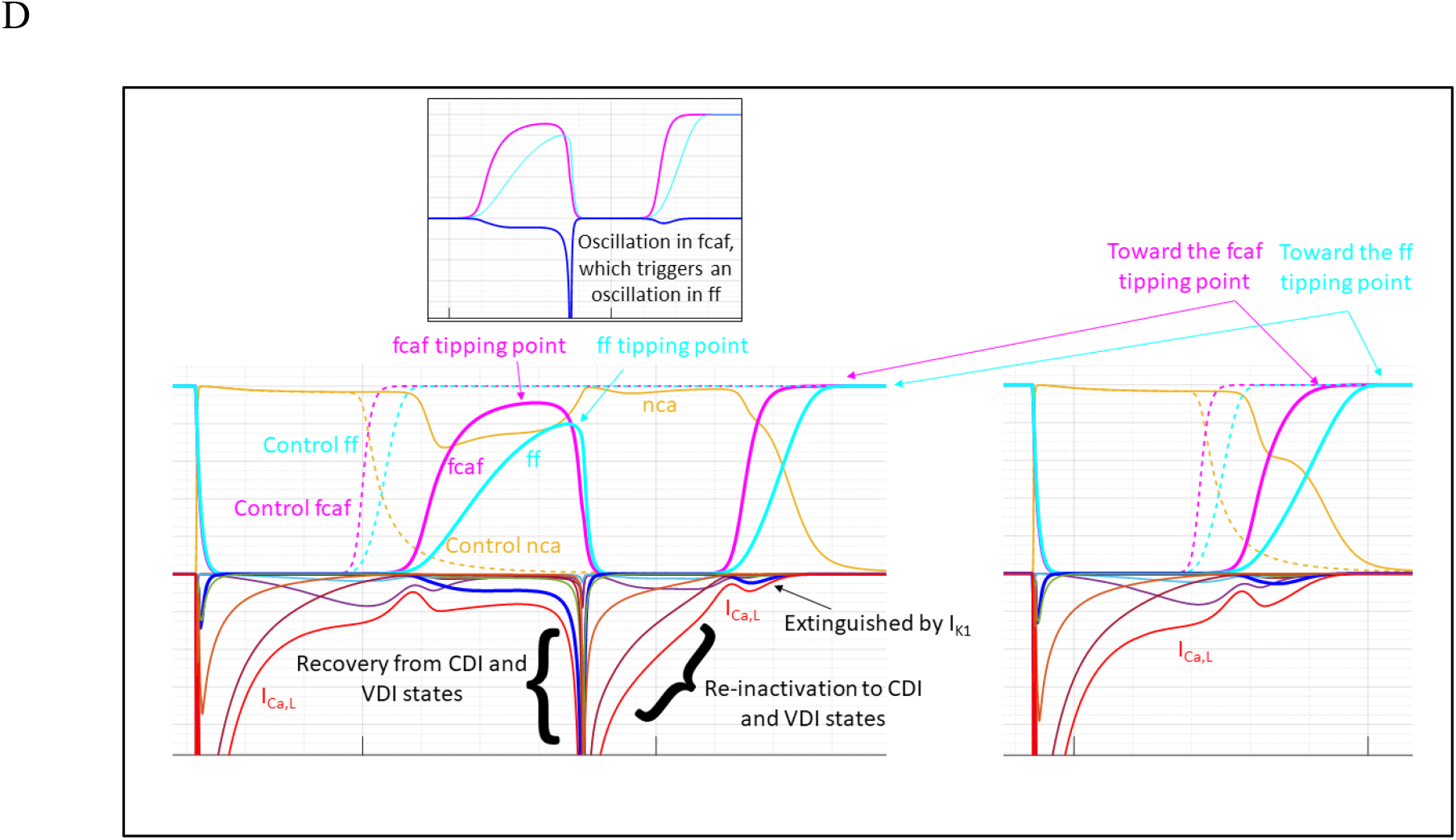
(A) A spontaneous depolarizing I_Ca,L_-driven spike occurs at a tipping point in *d*(Δ*ψ_m_*(*t*))/*dt*, fcaf, and ff slowing, coinciding with one or more oscillations in time-dependent gating behavior. A single oscillation occurs at the T_1_ threshold at this perturbation level, although a second *I_VCa,f_* window current appears at the end of the cycle (i.e. the channels are primed for an additional I_Ca,L_ spike at a higher perturbation level, and additional cycles of CDI/VDI inactivation, recovery, and _ICa,L_ spikes at still higher levels). (B) *I_Ca,L,act_* (purple tracing) at paced and spontaneous depolarizations in the AP complex shown in A and B. Reduced counter-balancing between inward and outward currents during atypical depolarizations results in significantly greater I_Ca,L_ (red arrows), compared with paced depolarizations under both perturbed and control conditions. (C) I_Ca,L_ window current-evoked Ca^2+^ release stimulates anomalous SR-mediated Ca^2+^ release (coral tracing) relative to control (green tracing), which further promotes population of the CDI states. nca(t) is determined by the rate of Ca^2+^-dependent buildup (*α*) versus recoverydependent decay *(nca · jca)* to/from the CDI states (*d*(*nca*(*t*))/*dt*). (D) Left panel: Full recoveryof Ca_v_1.2 channels from all CDI and VDI states occurs at a tipping point of fcaf and ff slowing (at which the two gates undergo a single opening-closing-reopening cycle terminated by I_K1_ activation), which underlies anomalous I_Ca,L_-driven depolarization. Right panel: *I_VCa,f_* partially recovers during the subsequent prolonged AP.

The overall pro-➔ pre-arrhythmic transitioning process can be summarized as follows:

1. The fast and slow Ca_v_1.2 inactivation gates reset to the open state in preparation for the subsequent AP. However, channel closing lags behind recovery from the VDI and CDI states due to decreased *d*(Δ*ψ_m_*(*t*))/*dt*, thereby expanding the I_Ca,L_ window current.
2. Transitions to/from the CDI states are governed by Ca^2+^-Ca_v_1.2 binding, which under control conditions, cycles between high and low levels during the AP (mirroring subspace Ca^2+^ levels). The Ca^2+^ cycle is disrupted under perturbed conditions, which promotes anomalous Ca^2+^-Ca_v_1.2 binding and transition to the CDI states (Figure 14C).
3. The fcaf gate slows in concert with the shoulder region of *d*(Δ*ψ_m_*(*t*))/*dt* under perturbed conditions.
4. fcaf oscillates at a critical perturbation level (corresponding to a threshold slowing of *d*(Δ*ψ_m_*(*t*))/*dt*), which drives recovery of *I_VCa,f_* and *I_V,f_*, which in turn drive recovery of all other CDI and VDI states. Anomalous *I_VCa,f_* is terminated by re-inactivation and channel closing coincident with I_K1_ activation (Figure 14D).
5. The fcaf oscillation may be extinguished by Kir2.1 activation below the T_2_ threshold, or undergo additional recovery/inactivation cycles at or above this threshold (the putative cause of the ringing oscillations observed under I_Kr_ knockout conditions (Figure 10)).

#### I_Na_ effects during the pro- and pre-arrhythmic stages

All time dependent Na_v_1.5 gating behaviors are perturbed as follows at hERG blocker concentrations < PT_1_ (Figure 15):

1. Time-dependent accumulation of phosphorylated Na_v_1.5, mirroring subspace Ca^2+^ levels and Ca^2+^-mediated CaMKII activation.
2. I_Na(early)_ follows the shift from the slower activating *I_Na,act_* to the faster activating phosphorylated 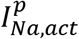 fraction across the 200 APs of the simulation (Figure 15A), which is offset by faster inactivation of 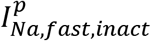 versus 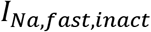 (Figure 15B). *I_Na,slow,inact_* and 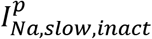 are weakly altered under perturbed conditions (Figure 15C). As a result, total INa(early) remains similar to control across the simulation (Figure 15D).

**Figure 15.**
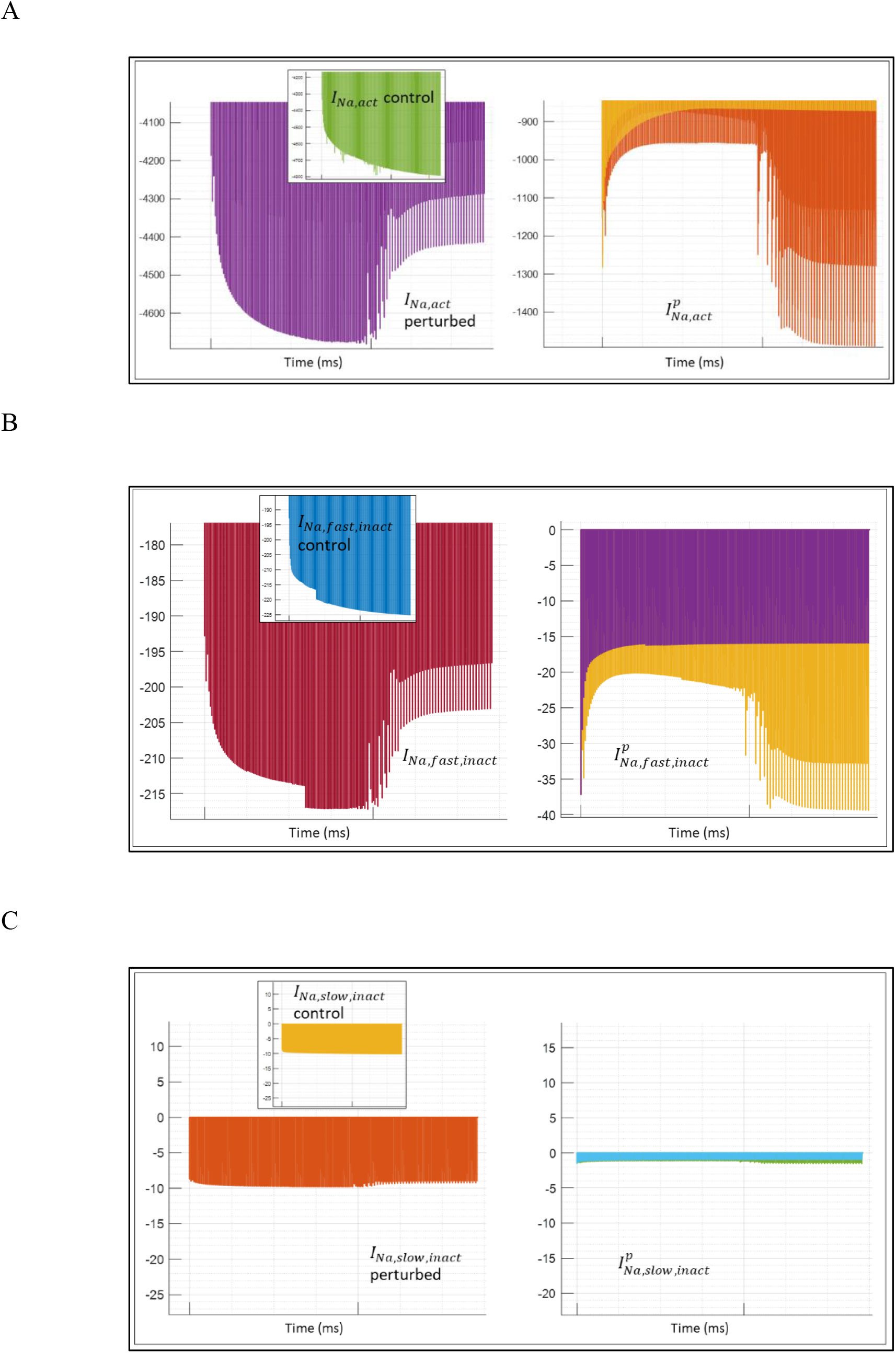
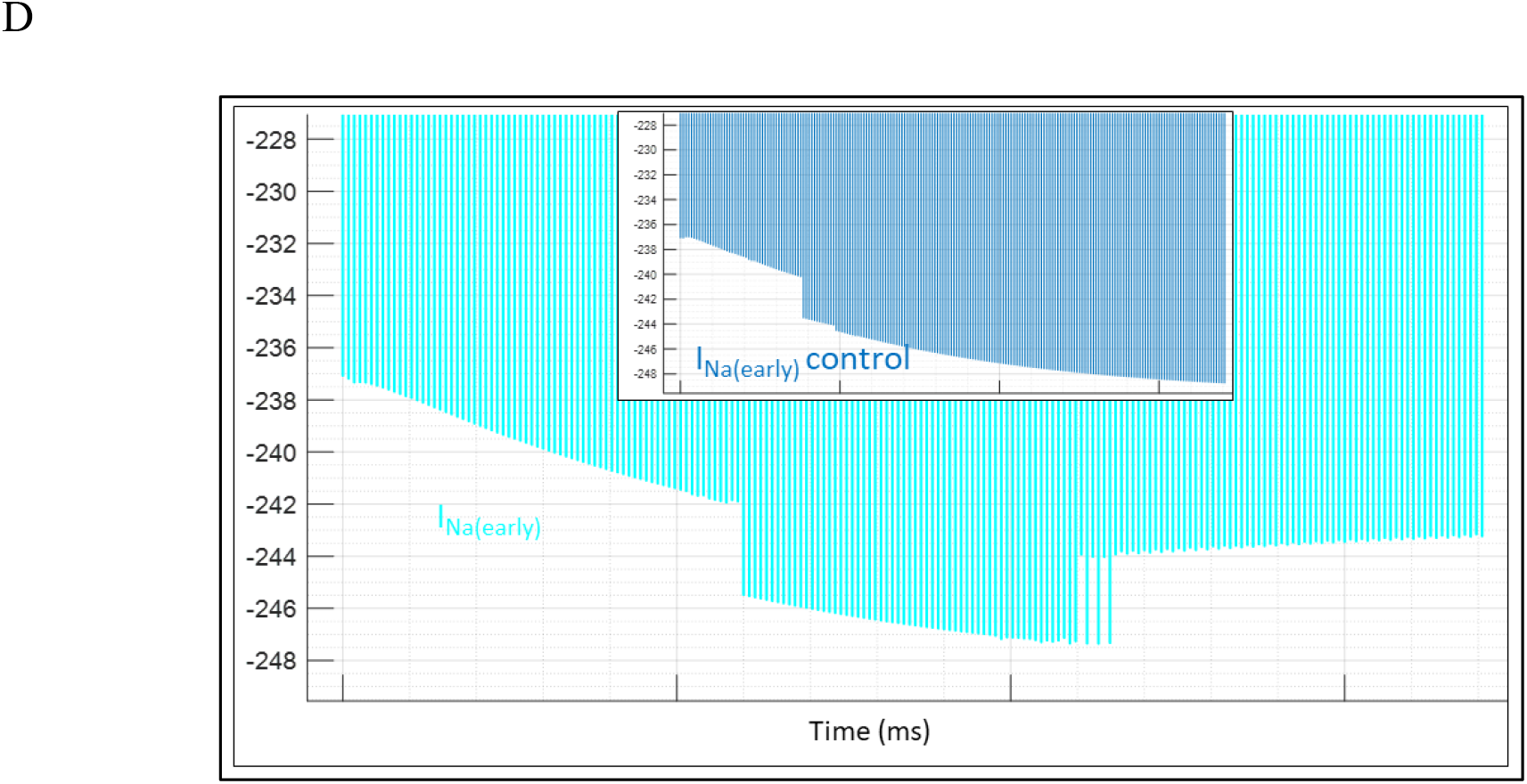
(A) *I_Na,act_* (left) and 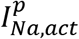 (right) under control (green inset and light orange tracings on the left and right, respectively) and perturbed (purple and dark orange tracings, respectively) conditons. Under perturbed conditions, 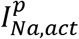 grows and *I_Na,act_* decreases with increasing time-dependent CaMKII-mediated Na_v_1.5 phosphorylation. (B) *I_Na,fast,inact_* (left) and 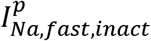 (right) under control (blue inset and purple tracings on the left and right, respectively) and perturbed conditons (auburn and light orange tracings, respectively). These currents likewise mirror increasing time-dependent Na_v_1.5 phosphorylation. (C) *I_Na,slow,inact_* (left) and 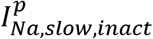 (right) under control (light orange inset and teal blue tracings on the left and right, respectively) and perturbed (dark orange and green tracings, respectively) conditons. As expected, the total I_Na(early)_ (the sum of the phosphorylated and non-phosphorylated contributions) is similar to control (not shown). (D) Total I_Na(early)_ under control (blue tracing) versus perturbed (cyan tracing) conditions are similar due to offsetting between the phosphorylated and non-phosphorylated forms of the channel.

I_Na(early)_ is absent entirely during atypical I_Ca,L_-driven depolarizations at blocker concentrations ≥ T_1_ (due to inactivation of Na_v_1.5 channels at the higher takeoff voltage of such depolarizations), whereas *h_L_* is open and INa(late) is present (Figure 16).

**Figure 16.**
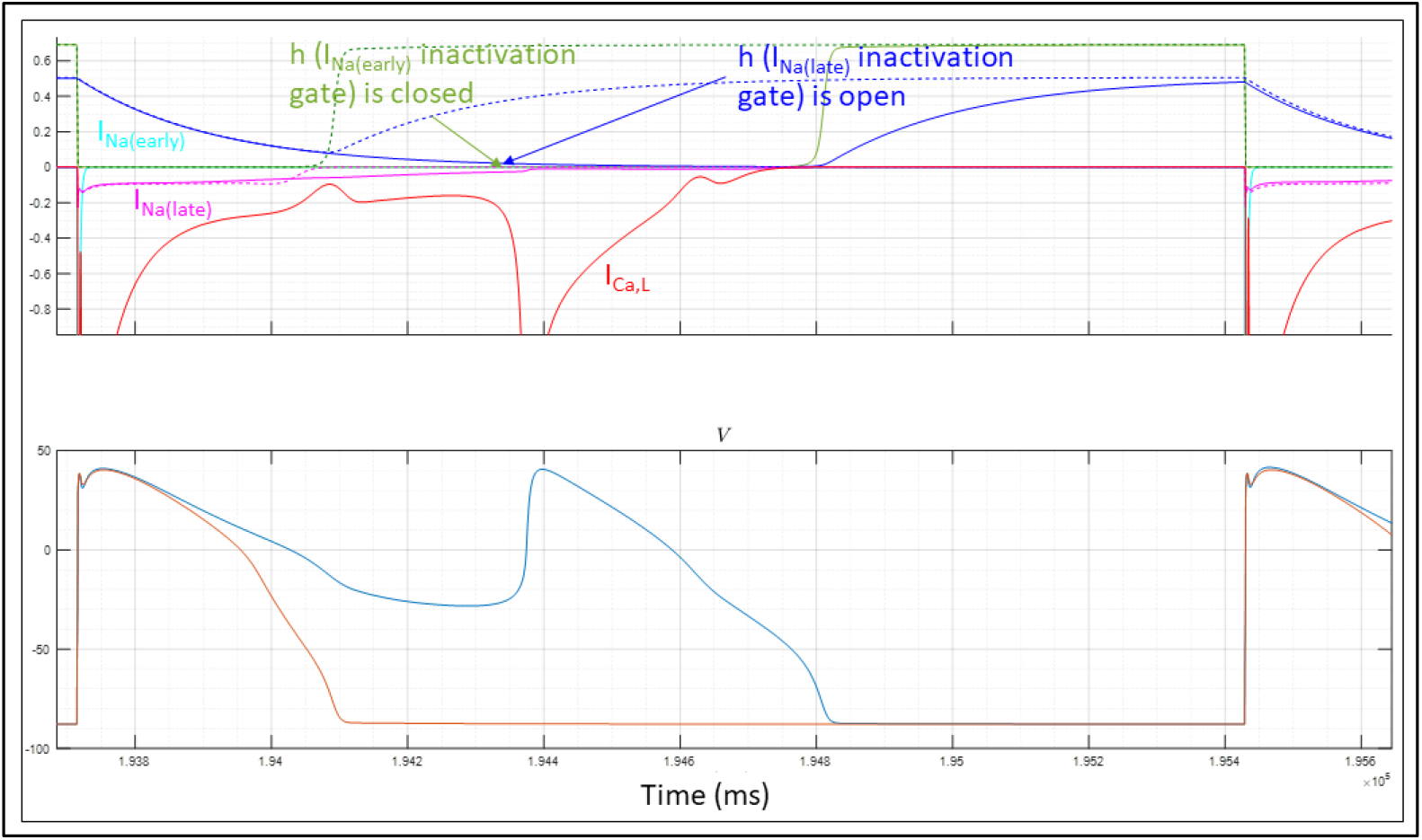
I_Na(early)_ (cyan tracing) is entirely absent during spontaneous I_Ca,L_-driven depolarizations, whereas I_Na(late)_ (solid magenta tracing, perturbed; dotted magenta tracing, control) is present. I_Ca,L_ (red tracing) and control I_Na(late)_ (auburn tracing) are shown for reference. The I_Na(early)_ inactivation gate (denoted as h) (solid green tracing, perturbed; dotted green tracing, control) is closed during spontaneous I_Ca,L_-driven depolarizations, whereas the I_Na(late)_ inactivation gate (denoted as hL) (solid blue tracing, perturbed; dotted blue tracing, control) is open.

#### I_Ca,L_ effects during the extreme arrhythmic stage (>> T_3_)

Next, we examined Ca_v_1.2 inactivation and recovery under total I_Kr_ knockout conditions, in which depolarization and repolarization are entirely I_Ca,L_- (triggered by paced signals) and I_Ks_-driven, respectively (noting that very small ion fluxes underlie the abnormally small voltage range of Δ*φ_m_*(*t*)). The ringing oscillations shown in Figure 10 are attributable largely to the interplay between Δ*ψ_m_*(*t*), the fcaf gate (depolarization), I_Ks_ (repolarization) (Figures 17A-B), and exchanger currents (not shown). The oscillations are mirrored by oscillations in subspace Ca^2+^ (Figure 17C).

**Figure 17.**
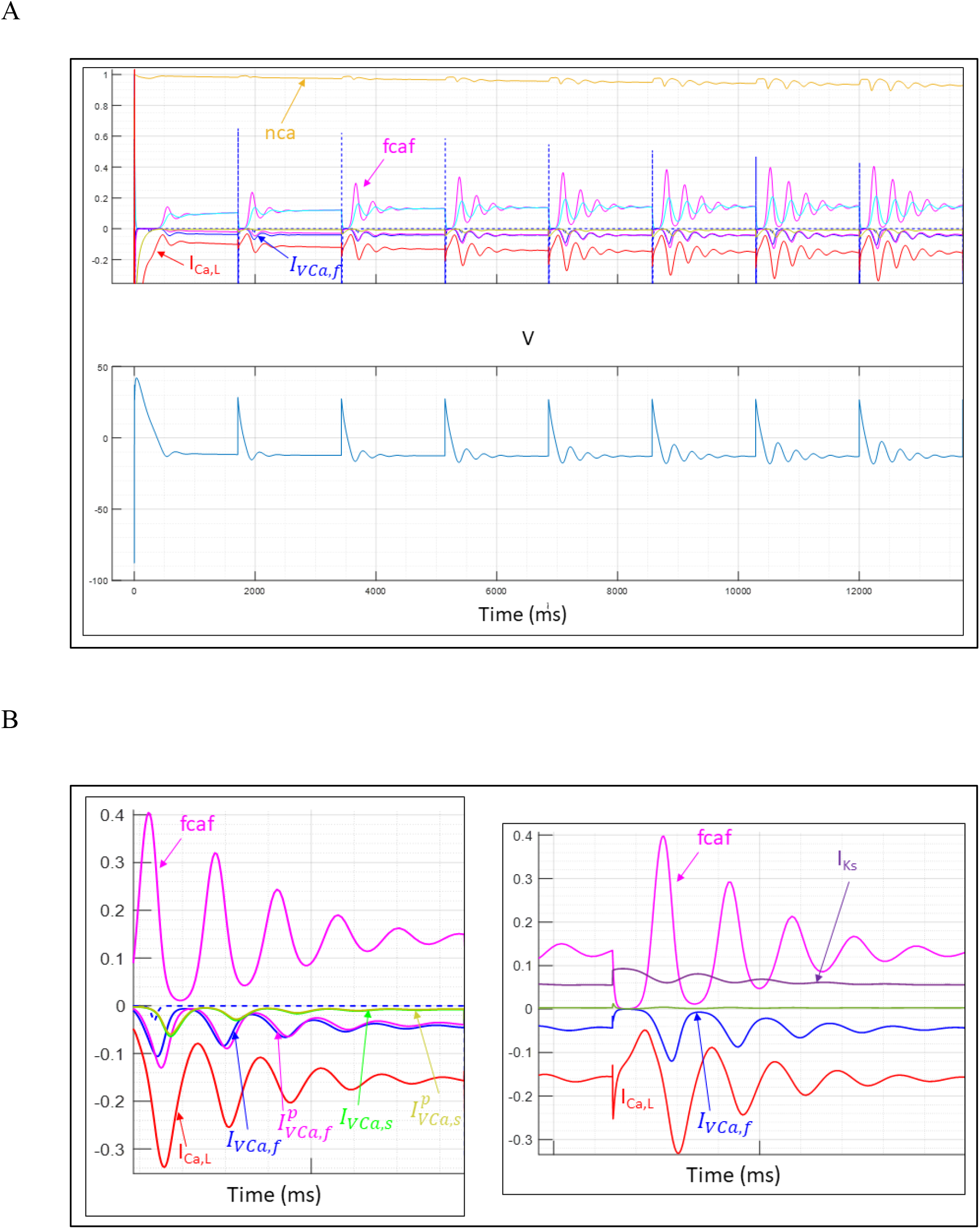
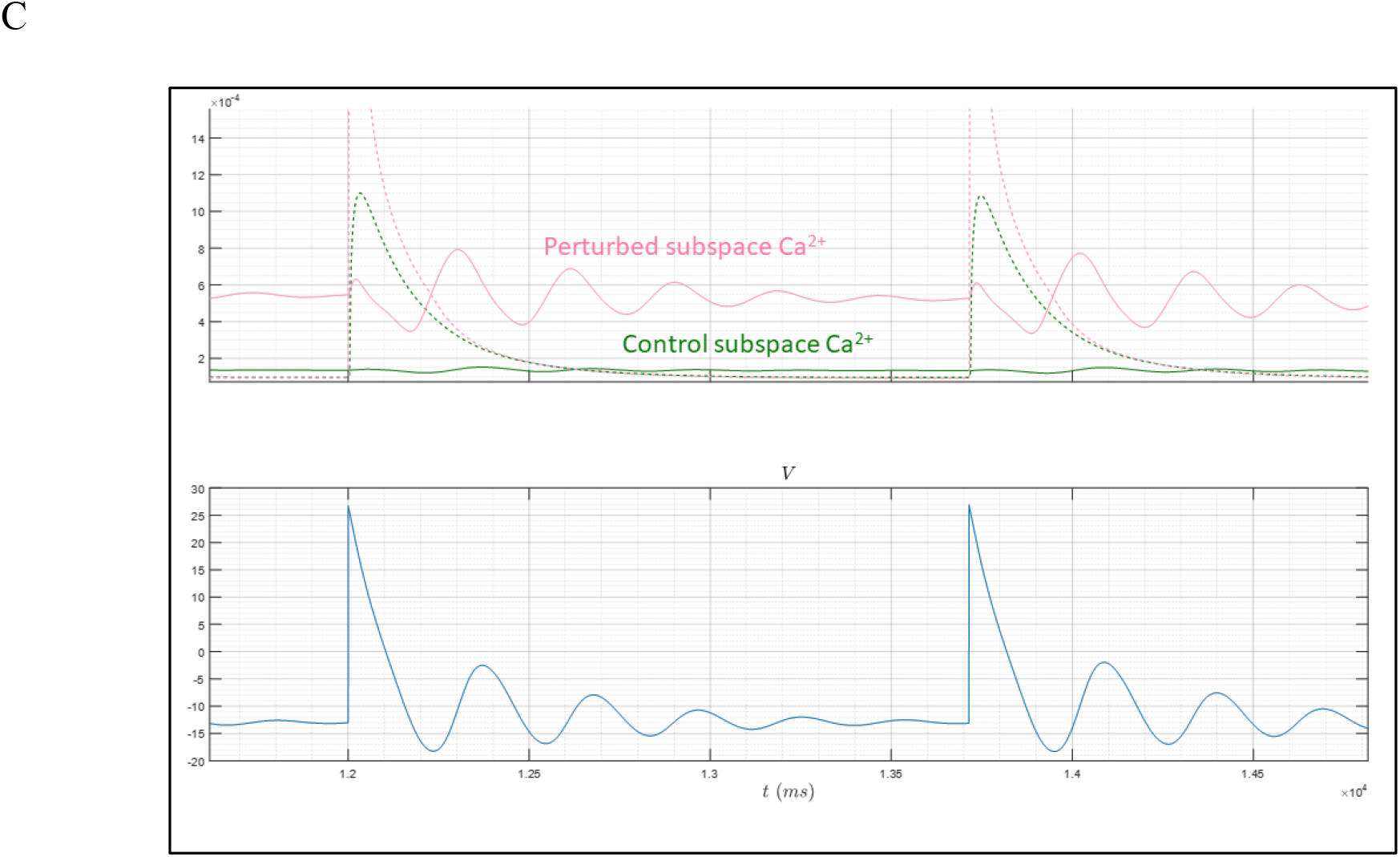
(A) Total I_Kr_ knockout-induced ringing oscillations in Δ*ψ_m_*(*t*) are driven entirely by recovery from the CDI and VDI states of Ca_v_1.2 (primarily CDI). (B) Zoomed out view of ringing oscillations in fcaf (magenta tracing) and the CDI state populations. (C) Dynamic subspace Ca^2+^ behavior under control (green tracing) and total I_Kr_ knockout conditions.

### The effects of hERG dysfunction on ion currents in the APD/CL ➔ 1 regime (hERG blocker concentration ➔ T_1_ @ CL = 1/60 min)

Next, we studied selected snapshots of ion current behaviors of paced AP_i_-AP_j_ collisions at CL = 1/60 min (51.945 ≤ hERG blocker concentration ≤ 51.95 nM) (Figure 8). Such collisions occur prior to the tipping point of spontaneous I_Ca,L(late)_ recovery, which is restricted to longer CL (1/35 min in our study). increasing perturbation severity results in AP_i_-AP_j_ collisions at increasing AP_i_ Δ*ψ_m_*(*t*) (the takeoff voltage of atypical AP_j_ depolarizations). The takeoff voltage ranges between −88 (normal resting) ≾ Δ*ψ_m_*(*t*) ≾ −30 mV. Atypical AP_j_ depolarizations are driven by I_Ca,L_/I_Na_ mixtures (Figure 18A), which are counter-balanced largely by I_K1_ and I_to_. The depolarization takeoff voltage increases with increasing perturbation severity, mirrored by increasing I_Na_ knockdown, I_to_ knockdown (~94% at blocker concentration ≈ 51.9504 nM and takeoff voltage = −75 mV (Figure 18B)), and I_K1_ knockdown (~50% at blocker concentration ≈ 51.9506 nM and takeoff voltage ≈ −60 mV; ~100% at blocker concentration ≈ 51.9510 nM and takeoff voltage ≈ −30 mV (Figure 18C)). Ca_v_1.2 channels are distributed in the VDI and CDI states of AP_i_ prior to the collision, which recover in response to AP_j_ stimuli.

**Figure 18.**
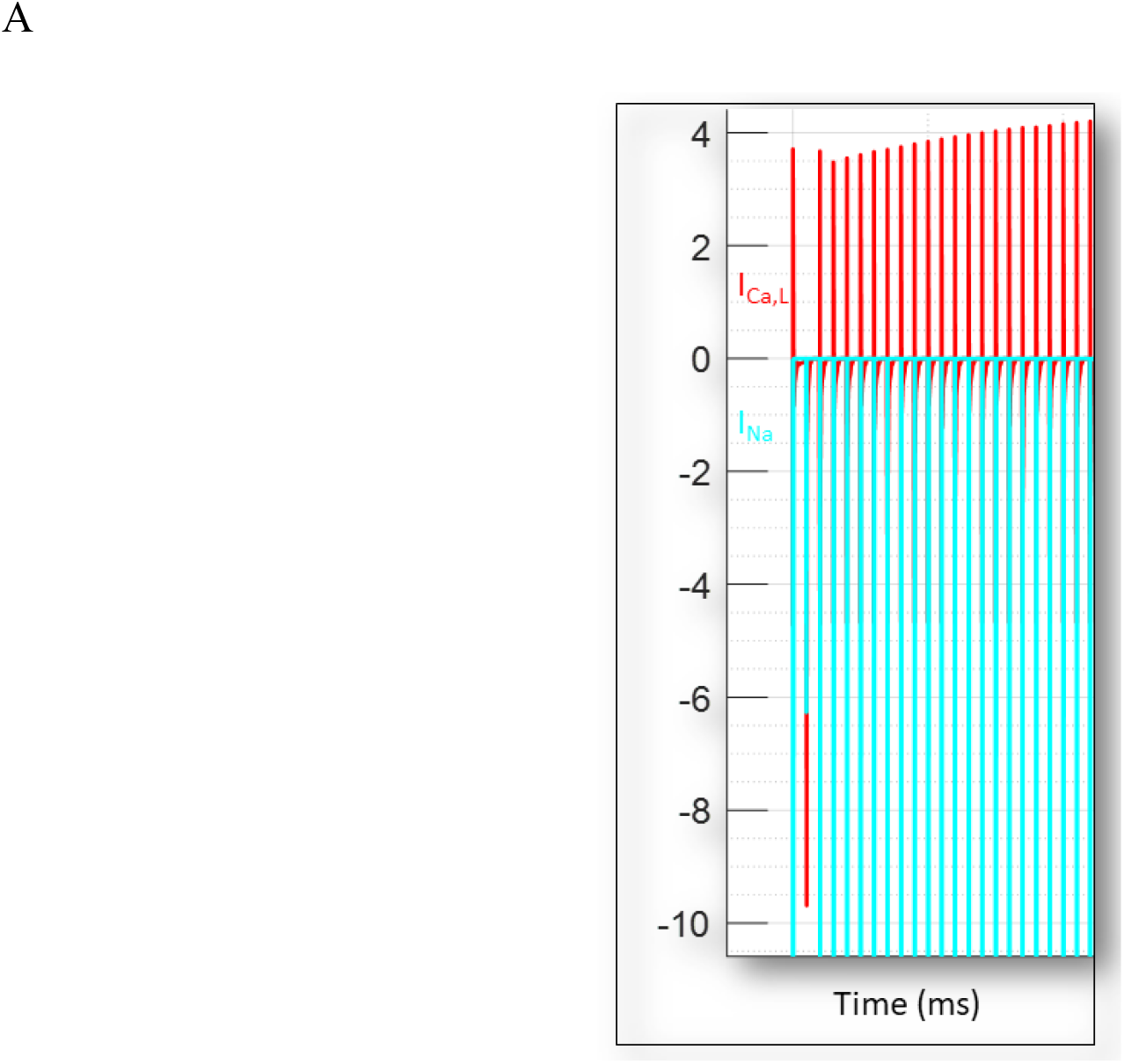
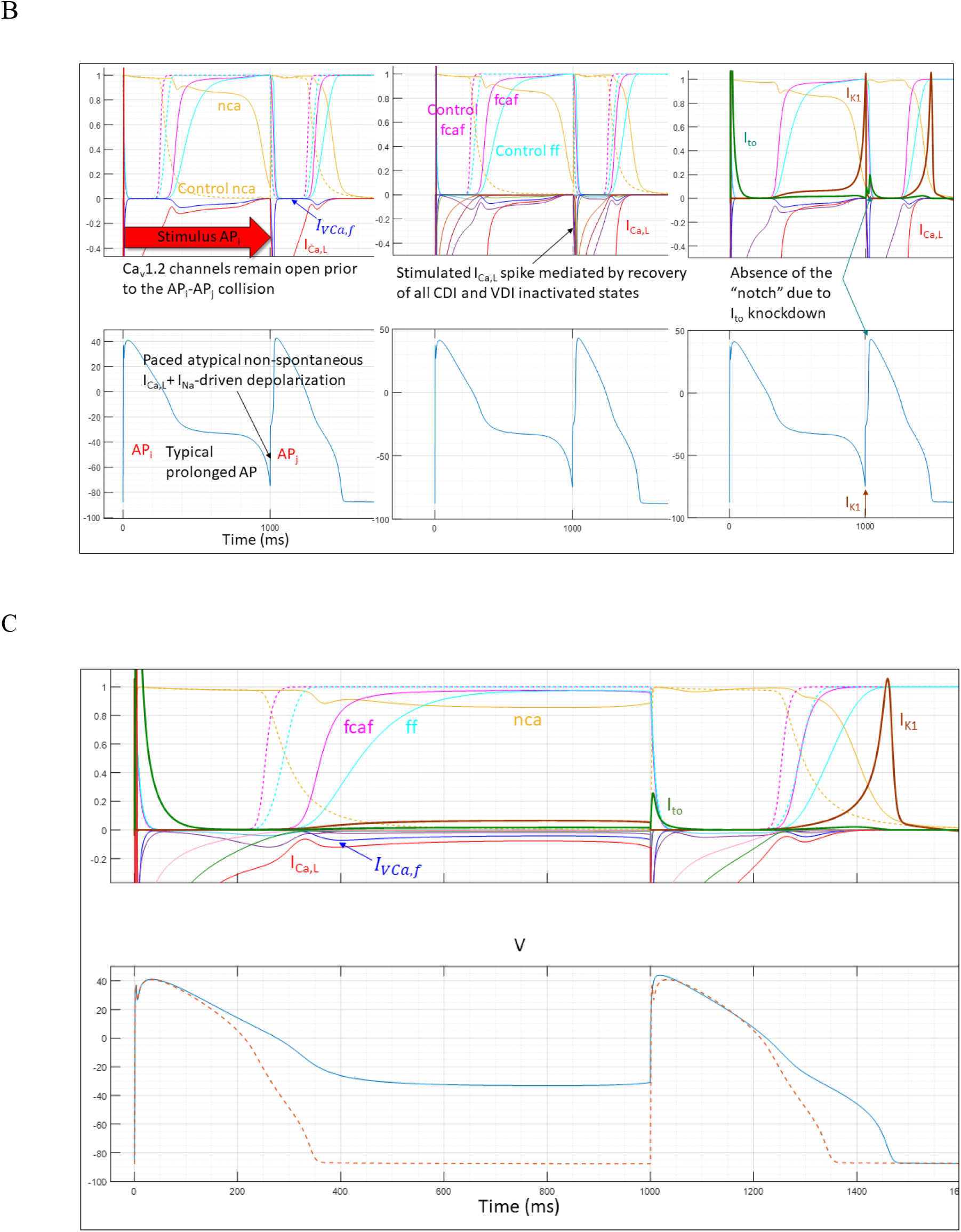
(A) I_Ca,L(peak)_ <u>increases</u> from −4.8 to −9.7 (red tracing) pA/pF, whereas I_Na_ <u>decreases</u> from −237 to −6.3 pA/pF at hERG blocker concentration = 51.9504 nM (cyan tracing) versus control, respectively. (B) I_Ca,L_, I_K1_, and I_to_ profiles at hERG blocker concentration = 51.9504 nM (takeoff voltage = −75 mV). I_K1_ remains similar to control, whereas I_to_ is knocked down from ~3.1 to 0.2 pA/pF under control versus perturbed conditions, respectively. (C) I_Ca,L_, I_K1_, and Ito profiles at hERG blocker concentration = 51.95101 nM (takeoff voltage = −30 mV). I_K1_ is knocked down to ~0, and Ito is knocked down from ~3.1 to 0.2 pA/pF under control versus perturbed conditions, respectively. I_Na(early)_ is likewise knocked out, whereas the magnitude of I_Na(late)_ is similar to control, but prolonged (not shown).

### The effects of hERG dysfunction on ion currents in the APD/CL > 1 regime (hERG blocker concentration ≥ T_2_ @ CL = 1/60 min)

Next, we studied the ion current profiles at selected snapshots (Figure 8) of the paced AP_j_-AP_k_ collision at ≥ T_2_ (51.9511 ≤ hERG blocker concentration ≤ 60.235 nM) (Figure 19). Similar results are expected at CL = 1/80 min. In this scenario, collisions result in Ca_v_1.2 reopening, rather than recovery.

**Figure 19.**
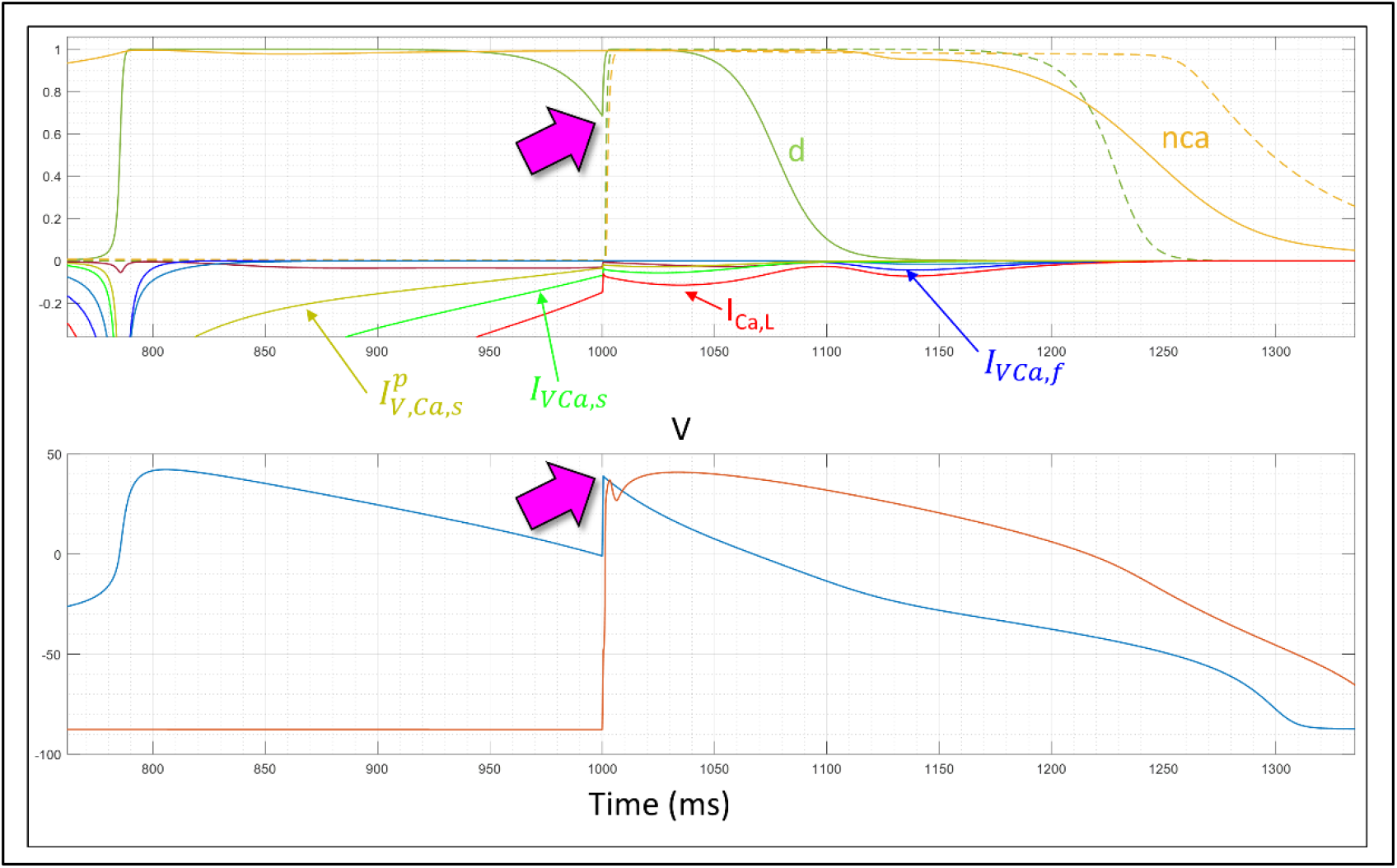
An atypical I_Ca,L(late)_-driven depolarization (AP_j_) collides with the following paced AP (AP_k_) at CL = 1/60 min and blocker concentration = 51.9565 nM (k_on_ = 1e8 M^-1^ s^-1^ and k_off_ = 2.0 s^-1^). This results in d gate reopening (light green tracing in the upper panel) in a small fraction of closed Ca_v_1.2 channels (magenta arrow in the upper panel), which evokes an additional atypical I_Ca,L(late)_-driven depolarization (magenta arrow in the lower panel) conveyed largely by *I_VCa,s_* (bright green tracing in the upper panel). This differs from atypical AP_j_ depolarizations, which are driven by recovery of the CDI and VDI states from inactivation (noting that nca remains constant during AP_j_-AP_k_ collisions).

### hERG blockade dose-response relationships

We monitored the following mutually inclusive pro-arrhythmia contributions as a function of hERG blocker concentration:

1. Maximum peak fractional hERG blocker occupancy (or magnitude of other acquired or inherited pro-arrhythmic modalities).
2. Maximum APD/CL and variability thereof across a large AP series (representative of the degree of instability of the AP generation system), noting that APD alone is an indirect metric of pro-arrhythmic propensity (i.e. a passenger, rather than a driver).
3. Minimum *d*(Δ*ψ_m_*(*t*))/*dt* centered at −30 mV (which mirrors the overall buildup of I_Ca,L_ during AP phase 3) and variability thereof across a large AP series.
4. Maximum fold-increase in |*I_VCa,f_* + *I_V,f_*| relative to control, and variability thereof across a large AP series.

We used ORd/hERG Markov simulations to characterize the aforementioned criteria for trappable and non-trappable hERG blockers in M cells as a function of concentration, BK, and CL (1/80, 1/60, 1/35 min). First, we used the automated algorithm described in Materials and methods to determine the PT_1_ threshold in each scenario, which serves as the endpoint in our dose-response simulations. The results, presented in Table 1 and Figures 20–23, may be summarized as follows:

1. Sensitivity of the AP generation system to perturbation grows with CL in order of 1/80 > 1/35 > 1/60 min, suggesting the existence of an anti-arrhythmic Goldilocks zone of CL centered at 1/60 min.
2. A “shockwave” of instability builds within the first ~100 APs at blocker concentration = PT_1_, settling to a new metastable state after ~450 APs (Figure 20G), suggesting that perturbation levels < T_1_ are buffered by the AP generation system.
3. Variability of APD/CL, *d*(Δ*ψ_m_*(*t*))/*dt*, and |*I_VCa,f_* + *I_V,f_*| grows as a function of time, which is mirrored by short-long-short sequences in the ECG preceding the onset of organ level arrhythmia.

**Table 1.** Magnitudes of the key pro-arrhythmia criteria at hERG blocker concentration = PT_1_ (determined iteratively via the algorithm described in Materials and methods) as a function of k_on_, k_off_, and CL at a fixed IC_50_ = 1.0 μM (calculated parameters at k_off_ < k_off_trappable_floor_ and k_off_non-trappable_floor_ have been omitted). Scalar quantities were calculated from the maxima of each parameter over the 200 APs of the simulations for control (denoted as C) and perturbed (denoted as (P) conditions. The quantities include: 1) maximum peak blocker occupancy (denoted as *γ*_T_1__) calculated using ORd/hERG Markov simulations and equilibrium occupancy (denoted as *γ*_eq_) calculated using the Hill equation (noting that *γ*_T_1__ = *γ*_eq_ for trappable blockers, and *γ*_T_1__ converges to *γ*_eq_ for non-trappable blockers exhibiting k_on_ ≳ 1e8 M^-1^ s^-1^); 2) PT_1_ (rounded to five decimal places); 3) minimum *d*(Δ*ψ_m_*(*t*))/*dt* during AP phase 3; 4) spread in AP slope represented by the number of histogram bins in the dataset; 5) maximum |*I_Ca,L_*|; 6) fold-increase in maximum total late |*I_VCa,f_* + |*I_V,f_*| over control (represented by the area under the current over time tracing); 7) spread in the maximum total late |*I_VCa,f_* + *I_V,f_*|; 8) maximum APD and APD/CL; 9) spread in the maximum APD.

|  |  | **1/35**<br>**(min)** | **1/60**<br>**(min)** | **1/80**<br>**(min)** | **k<sub>on</sub>**<br>**(M<sup>-1</sup> s<sup>-1</sup>)** | **k<sub>off</sub>**<br>**(s<sup>-1</sup>)** |
| --- | --- | --- | --- | --- | --- | --- |
| **T** | **Y<sub>T1</sub>, Y<sub>eq</sub> (%)** | 53.94, 53.94 | 60.11, 60.11 | 55.28, 55.28 | 6.93e5 | 0.693* |
|  | **PT<sub>1</sub> (nM)** | 1,171.24906 | 1,506.7279 | 1,236.3398 |  |  |
|  | **Min $d(\Delta\psi_m(t))/dt$**<br>**(mV/ms)** | -0.035 (P)<br>-0.499 (C) | -0.047 (P)<br>-0.533 (C) | -0.077 (P)<br>-0.534 (C) |  |  |
|  | **Distribution** | 25 (P)<br>11 (C) | 25 (P)<br>6 (C) | 34 (P)<br>9 (C) |  |  |
|  | **Max I<sub>Ca,L</sub>**<br>**(pA/pF)** | 4.79 (P)<br>4.78 (C) | 4.72 (P)<br>4.73 (C) | 5.98 (P)<br>5.0 (C) |  |  |
|  | **Max AUC total**<br>**$I_{Vca,f} + I_{V,f}$ (P/C)** | 6.66 | 2.79 | 1.95 |  |  |
|  | **Distribution** | 39 | 19 | 10 |  |  |
|  | **Max APD (ms)** | 920.83 (P)<br>396.13 (C) | 814.24 (P)<br>363.29 (C) | 708.15 (P)<br>361.39 (C) |  |  |
|  | **Distribution** | 44 (P)<br>7 (C) | 42 (P)<br>1 (C) | 37 (P)<br>4 (C) |  |  |
|  | **Max APD/CL** | 0.537 (P)<br>0.231 (C) | 0.814 (P)<br>0.363 (C) | 0.944 (P)<br>0.482 (C) |  |  |
| NT | **$Y_{T1}, Y_{eq}$ (%)** | 50.31, 68.93 | 53.13, 72.20 | 51.26, 71.0 | 2.0e6 | 2.0** |
|  | **$PT_1$ (nM)** | 2,218.66818 | 2,597.4567 | 2,448.6258 |  |  |
|  | **Min $d(\Delta\psi_m(t))/dt$ (mV/ms)** | -0.089 (P)<br>-0.499 (C) | -0.021 (P) -<br>0.533 (C) | -0.067 (P)<br>-0.534 (C) |  |  |
|  | **Distribution** | 16 (P)<br>11 (C) | 14 (P)<br>6 (C) | 29 (P)<br>9 (C) |  |  |
|  | **Max $I_{Ca,L}$ (pA/pF)** | 4.77 (P)<br>4.79 (C) | 6.0 (P)<br>4.73 (C) | 5.51 (P)<br>5.0 (C) |  |  |
|  | **Max AUC total $I_{VCa,f} + I_{V,f}$ (P/C)** | 5.29 | 1.99 | 2.07 |  |  |
|  | **Distribution** | 35 | 6 | 9 |  |  |
|  | **Max APD (ms)** | 762.13 (P)<br>396.13 (C) | 964.57 (P)<br>363.29 (C) | 689.17 (P)<br>361.39 (C) |  |  |
|  | **Distribution** | 34 (P)<br>7 (C) | 13 (P)<br>1 (C) | 34 (P)<br>4 (C) |  |  |
|  | **Max APD/CL** | 0.445 (P)<br>0.231 (C) | 0.965 (P)<br>0.363 (C) | 0.919 (P)<br>0.482 (C) |  |  |
| **T** | **$\gamma_{T_1}, \gamma_{eq}$ (%)** | 53.96, 53.96 | 59.92, 59.92 | 55.27, 55.27 | 2.0e6 | 2.0** |
|  | **PT<sub>1</sub> (nM)** | 1,171.80766 | 1,494.9204 | 1,235.7236 |  |  |
|  | **Min $d(\Delta\psi_m(t))/dt$ (mV/ms)** | -0.060 (P)<br>-0.499 (C) | -0.042 (P)<br>-0.533 (C) | -0.079 (P)<br>-0.533 (C) |  |  |
|  | **Distribution** | 21 (P)<br>11 (C) | 22 (P)<br>6 (C) | 30 (P)<br>9 (C) |  |  |
|  | **Max I<sub>Ca,L</sub> (pA/pF)** | 4.77 (P)<br>4.79 (C) | 4.94 (P)<br>4.73 (C) | 5.98 (P)<br>5.00 (C) |  |  |
|  | **Max AUC total $I_{Vca,f} + I_{V,f}$ (P/C)** | 4.88 | 2.61 | 1.94 |  |  |
|  | **Distribution** | 33 | 15 | 8 |  |  |
|  | **Max APD (ms)** | 821.28 (P)<br>396.13 (C) | 901.7 (P)<br>363.29 (C) | 706.48 (P)<br>361.39 (C) |  |  |
|  | **Distribution** | 38 (P)<br>7 (C) | 37 (P)<br>1 (C) | 33 (P)<br>4 (C) |  |  |
|  | **Max APD/CL** | 0.479 (P)<br>0.231 (C) | 0.902 (P)<br>0.363 (C) | 0.942 (P)<br>0.482 (C) |  |  |
| NT | **$\gamma_1, \gamma_{eq}$ (%)** | 57.1, 59.39 | 59.67, 62.13 | 56.47, 59.15 | 1.0e7 | 1.0e1 |
|  | **PT<sub>1</sub> (nM)** | 1,462.17 | 1,640.3048 | 1,447.9159 |  |  |
|  | **Min $d(\Delta\psi_m(t))/dt$ (mV/ms)** | -0.034 (P)<br>-0.499 (C) | -0.03 (P)<br>-0.534 (C) | -0.068 (P)<br>-0.534 (C) |  |  |
|  | **Distribution** | 20 (P)<br>11 (C) | 15 (P)<br>6 (C) | 30 (P)<br>9 (C) |  |  |
|  | **Max I<sub>Ca,L</sub> (pA/pF)** | 4.77 (P)<br>4.79 (C) | 6.0 (P)<br>4.73 (C) | 5.98 (P)<br>5.00 (C) |  |  |
|  | **Max AUC total $I_{Vca,f} + I_{V,f}$ (P/C)** | 5.43 | 1.98 | 2.05 |  |  |
|  | **Distribution** | 35 | 7 | 9 |  |  |
|  | **Max APD (ms)** | 873.87 (P)<br>396.13 (C) | 964.86 (P)<br>363.29 (C) | 705.92 (P)<br>361.39 (C) |  |  |
|  | **Distribution** | 37 (P)<br>7 (C) | 15 (P)<br>1 (C) | 34 (P)<br>4 (C) |  |  |
|  | **Max APD/CL** | 0.051 (P)<br>0.231 (C) | 0.965 (P)<br>0.363 (C) | 0.941 (P)<br>0.482 (C) |  |  |
| **T** | **$\gamma_{T1}, \gamma_{eq}$ (%)** | 53.92, 53.92 | 57.7, 57.7 | 55.27, 55.27 | 1.0e7 | 1.0e1 |
|  | **$PT_1$** | 1,170.22907 | 1,364.095 | 1,235.5668 |  |  |
|  | **Min $d(\Delta\psi_m(t))/dt$**<br>**(mV/ms)** | -0.074<br>-0.499 | -0.04 (P)<br>-0.533 (C) | -0.079 (P)<br>-0.534 (C) |  |  |
|  | **Distribution** | 18 (P)<br>11 (C) | 14 (P)<br>6 (C) | 27 (P)<br>9 (C) |  |  |
|  | **Max $I_{Ca,L}$**<br>**(pA/pF)** | 4.77 (P)<br>4.79 (C) | 6.0 (P)<br>4.73 (P) | 5.92 (P)<br>5.0 (C) |  |  |
|  | **Max AUC total**<br>**$I_{Vca,f} + I_{V,f}$ (P/C)** | 4.65 | 1.86 | 1.94 |  |  |
|  | **Distribution** | 31 | 6 | 8 |  |  |
|  | **Max APD (ms)** | 778.19 (P)<br>396.13 (C) | 964.14 (P)<br>363.29 (C) | 704.8 (P)<br>361.39 (C) |  |  |
|  | **Distribution** | 37 (P)<br>7 (C) | 13 (P)<br>1 (C) | 34 (P)<br>4 (C) |  |  |
|  | **Max APD/CL** | 0.454 (P)<br>0.231 (C) | 0.964 (P)<br>0.363 (C) | 0.94 (P)<br>0.482 (C) |  |  |
| NT | **$\gamma_{T_1}, \gamma_{eq}$ (%)** | 54.49, 54.62 | 58.18, 58.31 | 55.61, 55.74 | 1.0e8 | 1.0e2 |
|  | **PT<sub>1</sub> (nM)** | 1,203.3485 | 1,398.5543 | 1,259.2552 |  |  |
|  | **Min $d(\Delta\psi_m(t))/dt$ (mV/ms)** | -0.077 (P)<br>-0.499 (C) | -0.039 (P)<br>-0.533 (C) | -0.076 (P)<br>-0.534 (C) |  |  |
|  | **Distribution** | 18 (P)<br>11 (C) | 14 (P)<br>6 (C) | 29 (P)<br>9 (C) |  |  |
|  | **Max I<sub>Ca,L</sub> (pA/pF)** | 4.79 (P)<br>4.77 (C) | 6.0 (P)<br>4.73 (C) | 5.81 (P)<br>5.0 (C) |  |  |
|  | **Max AUC total $I_{Vca,f} + I_{V,f}$ (P/C)** | 4.66 | 1.89 | 1.95 |  |  |
|  | **Distribution** | 31 | 6 | 8 |  |  |
|  | **Max APD (ms)** | 784.1 (P)<br>396.13 (C) | 964.40 (P)<br>363.29 (C) | 701.71 (P)<br>361.39 (C) |  |  |
|  | **Distribution** | 35 (P)<br>7 (C) | 14 (P)<br>1 (C) | 35 (P)<br>4 (C) |  |  |
|  | **Max APD/CL** | 0.457 (P)<br>0.231 (C) | 0.964 (P)<br>0.363 (C) | 0.936 (P)<br>0.482 (C) |  |  |
| T | **$\gamma_{T1}, \gamma_{eq}$ (%)** | 53.96, 53.96 | 57.77, 57.77 | 55.26, 55.26 | 1.0e8 | 1.0e2 |
|  | **PT<sub>1</sub> (nM)** | 1,171.8390 | 1,367.7385 | 1,235.301 |  |  |
|  | **Min $d(\Delta\psi_m(t))/dt$ (mV/ms)** | -0.033 (P) | -0.04 (P) | -0.078 (P) |  |  |
|  |  | -0.499 (C) | -0.533 (C) | -0.534 (C) |  |  |
|  | **Distribution** | 21 (P) | 13 (P) | 29 (P) |  |  |
|  |  | 11 (C) | 6 (C) | 9 (C) |  |  |
|  | **Max I<sub>Ca,L</sub> (pA/pF)** | 4.77 (P) | 6.0 (P) | 5.95 (P) |  |  |
|  |  | 4.79 (C) | 4.73 (C) | 5.0 (C) |  |  |
|  | **Max AUC total $I_{Vca,f} + I_{V,f}$ (P/C)** | 5.40 | 1.86 | 1.94 |  |  |
|  | **Distribution** | 35 | 6 | 8 |  |  |
|  | **Max APD (ms)** | 923.15 (P) | 964.74 (P) | 706.27 (P) |  |  |
|  |  | 396.13 (C) | 363.29 (C) | 361.39 (C) |  |  |
|  | **Distribution** | 40 (P) | 13 (P) | 33 (P) |  |  |
|  |  | 7 (C) | 1 (C) | 4 (C) |  |  |
|  | **Max APD/CL** | 0.539 (P) | 0.965 (P) | 0.942 (P) |  |  |
|  |  | 0.231 (C) | 0.363 (C) | 0.482 (C) |  |  |
| NT | **$\gamma_{T_1}, \gamma_{eq}$ (%)** | 53.92, 54.0 | 57.73, 57.81 | 55.23, 55.31 | 1.0e9 | 1.0e3 |
|  | **PT<sub>1</sub> (nM)** | 1,173.73975 | 1,370.41727 | 1,237.5011 |  |  |
|  | **Min $d(\Delta\psi_m(t))/dt$ (mV/ms)** | -0.066 (P)<br>-0.499 (C) | -0.037 (P)<br>-0.535 (C) | -0.078 (P)<br>-0.534 (C) |  |  |
|  | **Distribution** | 19 (P)<br>11 (C) | 13 (P)<br>6 (C) | 28 (P)<br>9 (C) |  |  |
|  | **Max I<sub>Ca,L</sub> (pA/pF)** | 4.77 (P)<br>4.79 (C) | 6.0 (P)<br>4.73 (C) | 5.95 (P)<br>5.0 (C) |  |  |
|  | **Max AUC total $I_{Vca,f} + I_{V,f}$ (P/C)** | 4.78 | 1.87 | 1.95 |  |  |
|  | **Distribution** | 32 | 6 | 8 |  |  |
|  | **Max APD (ms)** | 802.65 (P)<br>396.13 (C) | 964.72 (P)<br>363.29 (C) | 705.55 (P)<br>361.39 (C) |  |  |
|  | **Distribution** | 37 (P)<br>7 (C) | 14 (P)<br>1 (C) | 35 (P)<br>4 (C) |  |  |
|  | **Max APD/CL** | 0.468 (P)<br>0.231 (C) | 0.965 (P)<br>0.363 (C) | 0.941 (P)<br>0.482 (C) |  |  |
| **T** | **$\gamma_{T_1}, \gamma_{eq}$ (%)** | 53.93, 53.93 | 57.77, 57.77 | 55.26, 55.26 | 1.0e9 | 1.0e3 |
|  | **$PT_1$ (nM)** | 1,170.52515 | 1,368.07097 | 1,235.244 |  |  |
|  | **Min $d(\Delta\psi_m(t))/dt$ (mV/ms)** | -0.073 (P)<br>-0.507 (C) | -0.04 (P)<br>-0.535 (C) | -0.079 (P)<br>-0.534 (C) |  |  |
|  | **Distribution** | 18 (P)<br>11 (C) | 14 (P)<br>6 (C) | 29 (P)<br>9 (C) |  |  |
|  | **Max $I_{Ca,L}$ (pA/pF)** | 4.69 (P)<br>4.79 (C) | 6.0 (P)<br>4.73 (C) | 5.97 (P)<br>5.00 (C) |  |  |
|  | **Max AUC total $I_{Vca,f} + I_{V,f}$ (P/C)** | 4.59 | 1.86 | 1.95 |  |  |
|  | **Distribution** | 31 | 6 | 8 |  |  |
|  | **Max APD (ms)** | 795.06 (P)<br>396.13 (C) | 964.40 (P)<br>363.29 (C) | 707.24 (P)<br>361.39 (C) |  |  |
|  | **Distribution** | 37 (P)<br>7 (C) | 13 (P)<br>1 (C) | 35 (P)<br>4 (C) |  |  |
|  | **APD/CL** | 0.464 (P)<br>0.231 (C) | 0.964 (P)<br>0.363 (C) | 0.943 (P)<br>0.482 (C) |  |  |

\* $k_{off\_trappable\_floor}$ ; \*\* $k_{off\_non-trappable\_floor}$ .

**Figure 20.**
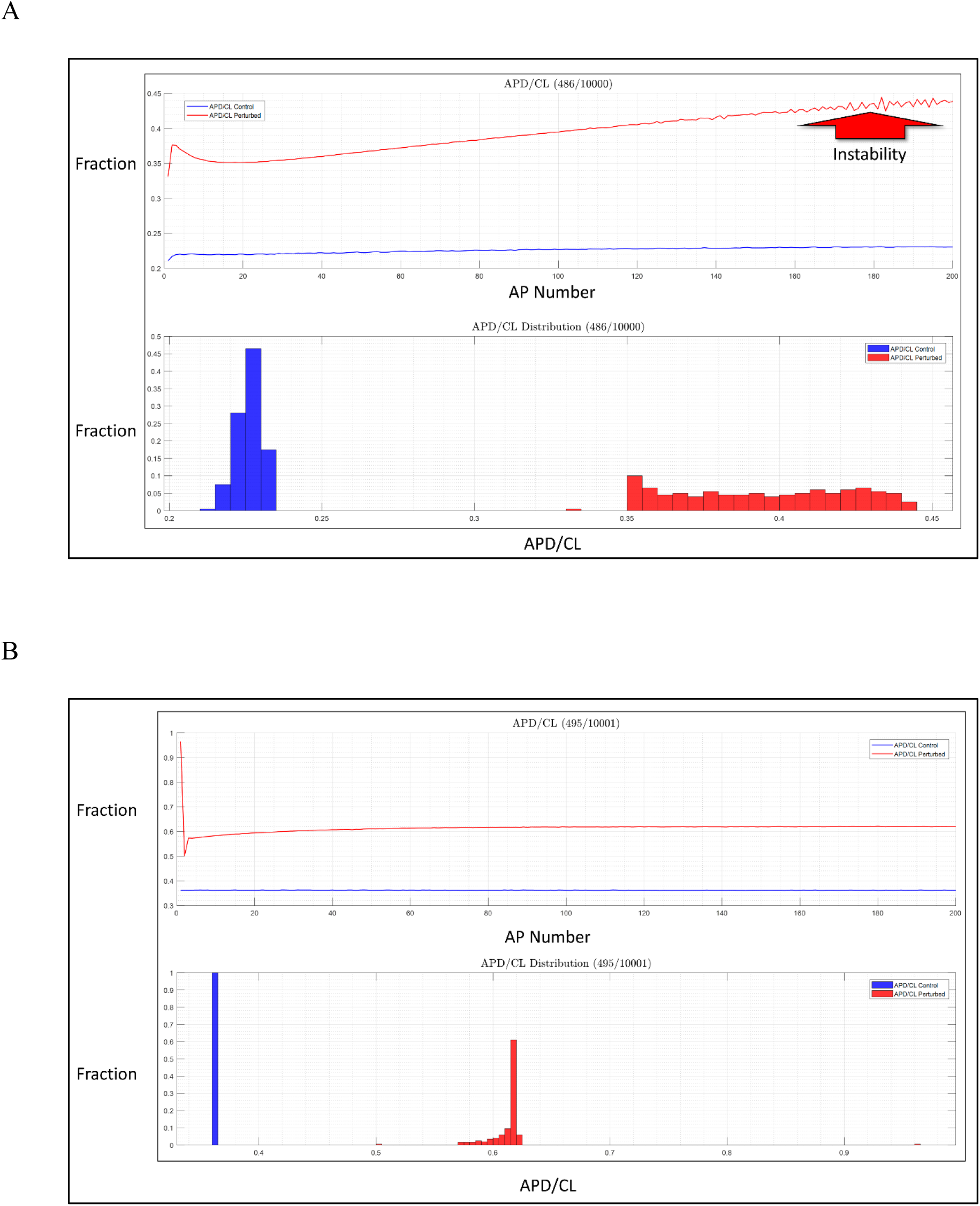
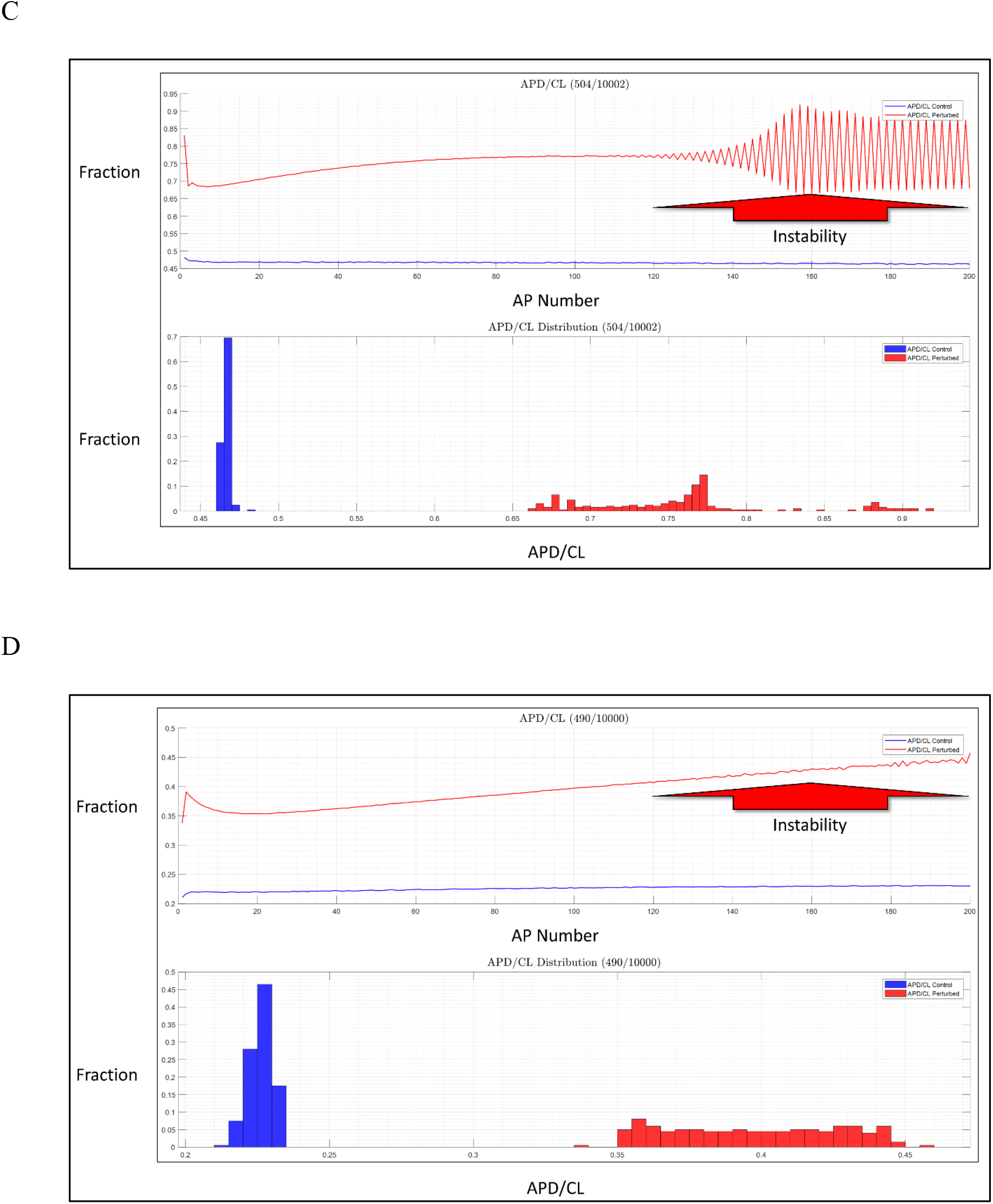
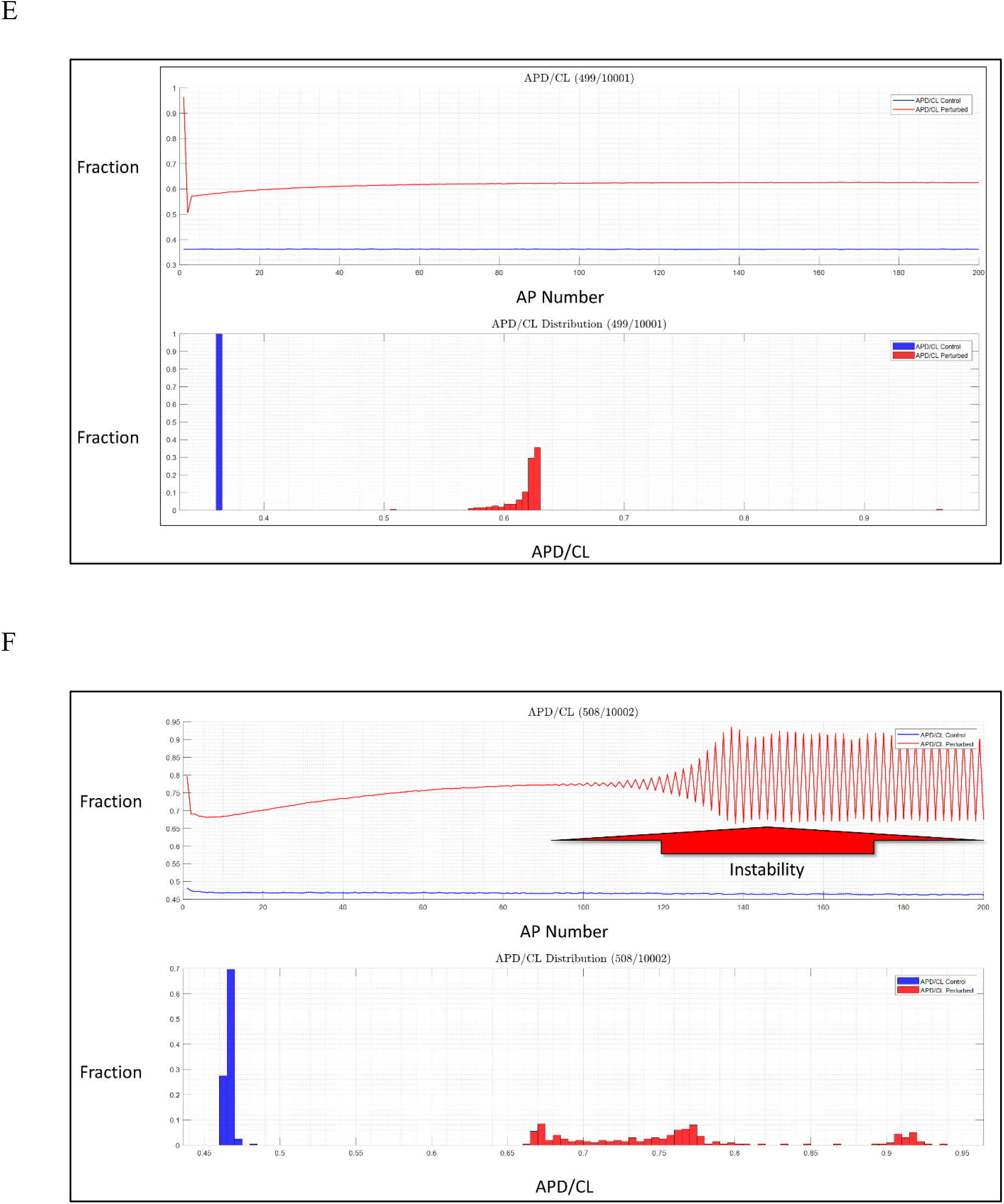
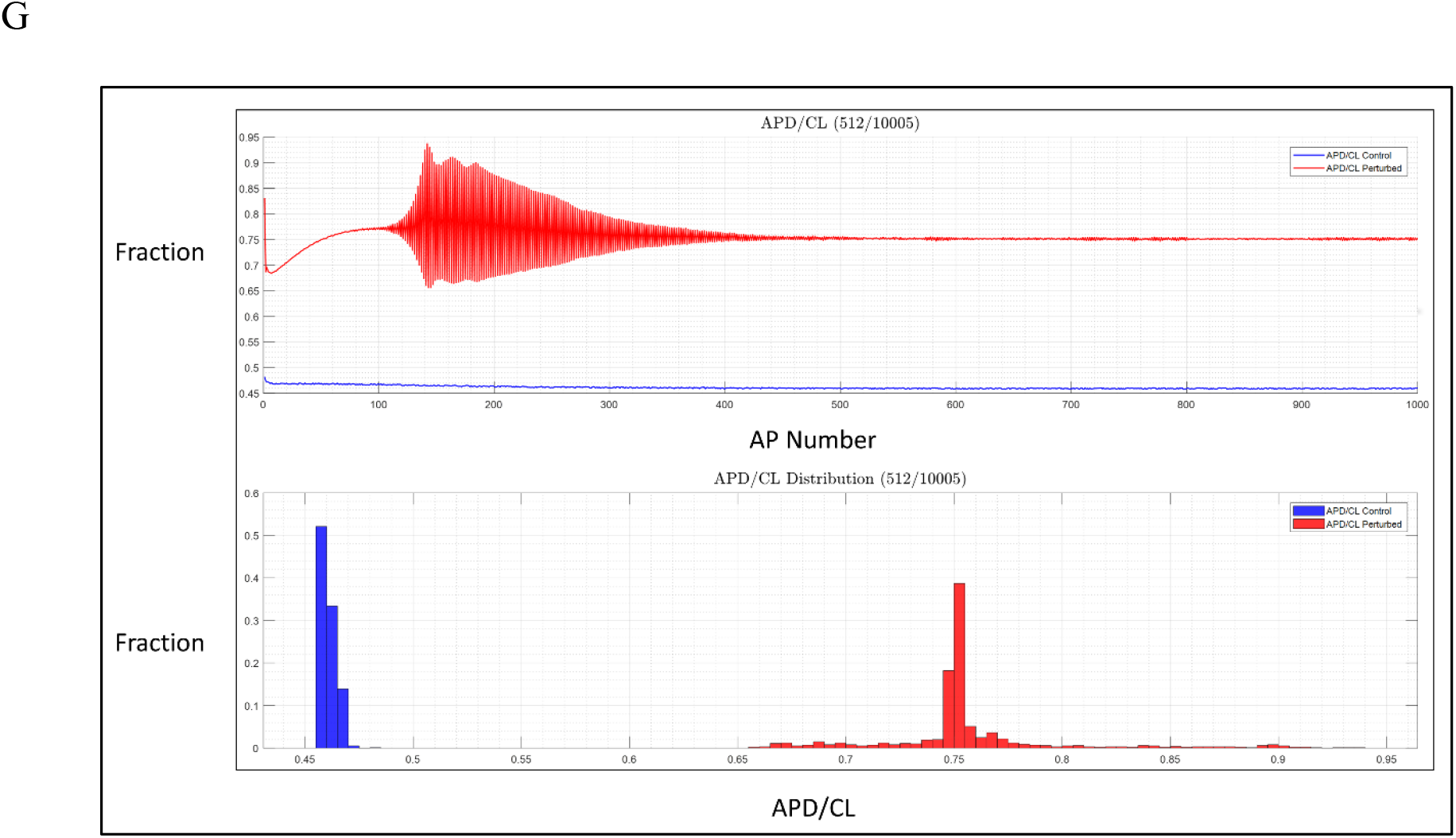
(A) Upper panel: APD/CL at CL = 1/35 min for control (blue tracing) versus non-trappable hERG blocker k_on_ = 2.0e6 M^-1^ s^-1^, k_off_ = 2.0 s^-1^, blocker concentration = PT_1_ = 2,218.66818 nM rounded to five decimal places (blue tracing). Oscillations in APD/CL build throughout the latter half of the simulation (red arrow), reflecting increasing instability of the system, which nevertheless remains non-arrhythmic at this perturbation level. Lower panel: Distribution of APD/CL across the 200 APs of the simulation under control versus perturbed conditions (blue and red bars, respectively). The broad distribution of APD/CL under perturbed conditions reflects high instability of the system in the PT_1_ regime. (B) Same as A, except CL = 1/60 min and PT_1_ = 2,597.4567 nM. (C) Same as A, except CL = 1/80 min and PT_1_ = 2,448.6258 nM. (D) Same as A, except k_on_ = 1.0e8 M^-1^ s^-1^, k_off_ = 2.0 s^-1^, and PT_1_ = 1,203.3485 nM. (E) Same as D, except CL = 1/60 min and PT_1_ = 1,398.5543 nM. (F) Same as D, except CL = 1/80 min and PT_1_ = 1,259.2552 nM. (G) Same as C, except PT_1_ = 2,448.1306 nM and the number of simulated APs = 1,000. Overt instability present in the previous tracings (resembling a shockwave) settles to a new metastable non-arrhythmic steady state after ~450 APs. We further note that PT_1_ is significantly higher at 60 versus 35 and 80 BPM (Table 1).

**Figure 21.**
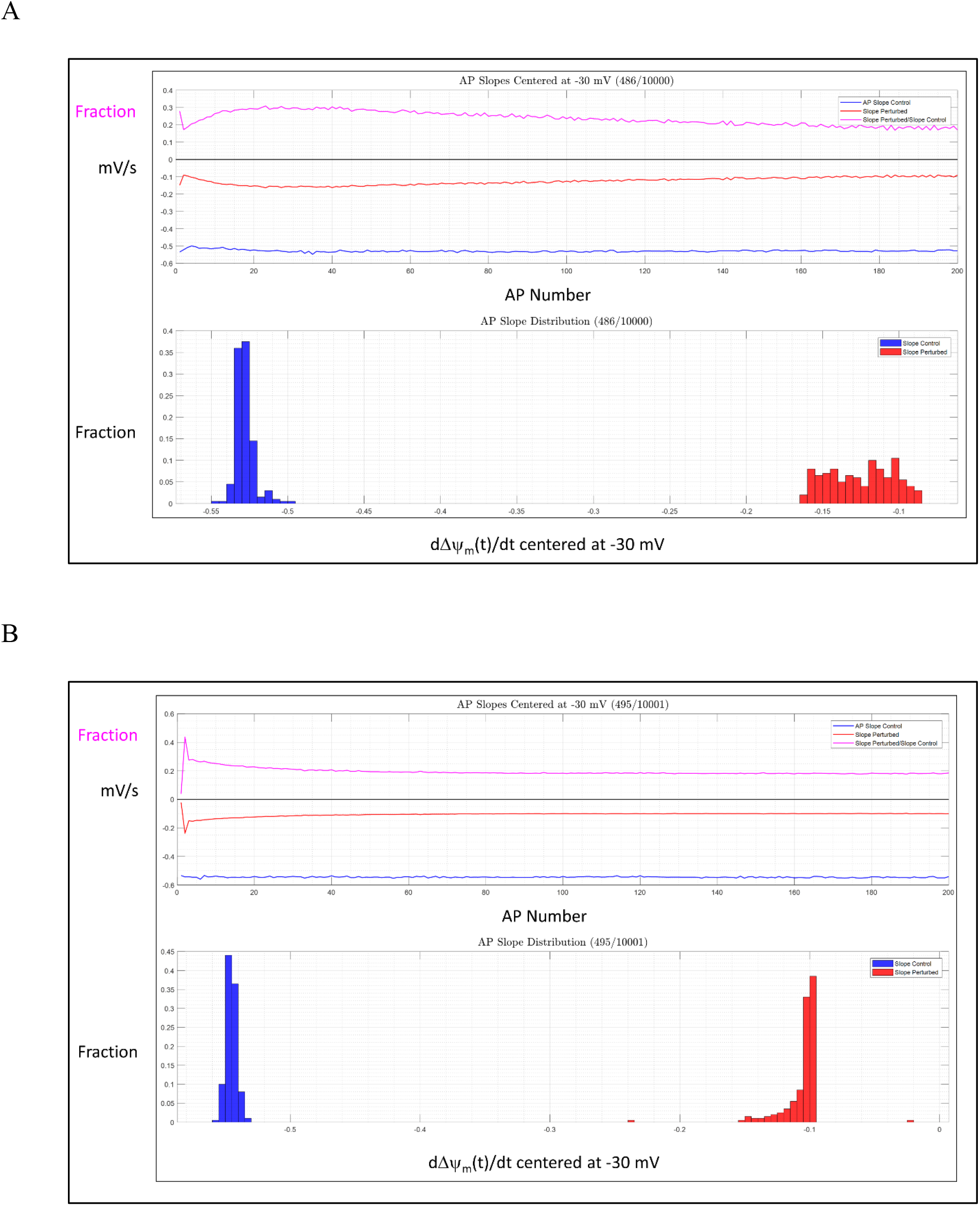
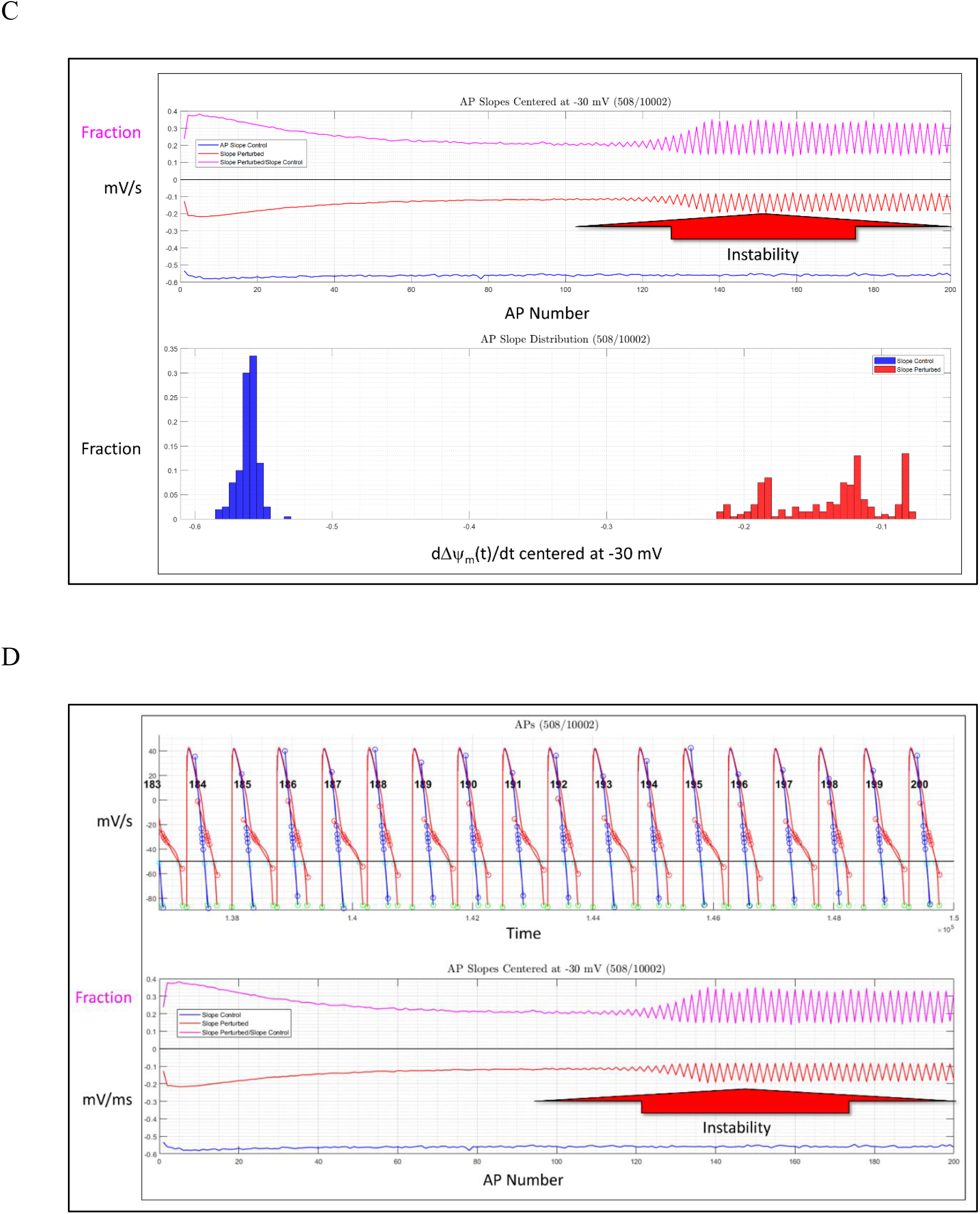
(A) Upper panel: Same as Figure 20A, except showing control versus perturbed *d*(Δ*ψ_m_*(*t*))/*dt* (blue and red tracings, respectively). Fold-difference in perturbed/control is shown in magenta. Lower panel: Same as Figure 20A, except showing control versus perturbed *d*(Δ*ψ_m_*(*t*)) / *dt*. (B) Same as A, except CL = 1/60 min and PT_1_ = 2,597.4567 nM. (C) Same as A, except CL = 1/80 min and PT_1_ = 2,448.6258 nM. (D) Same as C, except showing the last 16/200 APs of the simulation under control (blue) versus perturbed (red) conditions, with *d*(Δ*ψ_m_*(*t*))/*dt* tangent lines (red vectors) superimposed on the perturbed APs (slope calculation method described in Materials and methods). The AP numbers in the upper panel (corresponding to those on the x-axis of the lower panel) are shown in bold.

**Figure 22.**
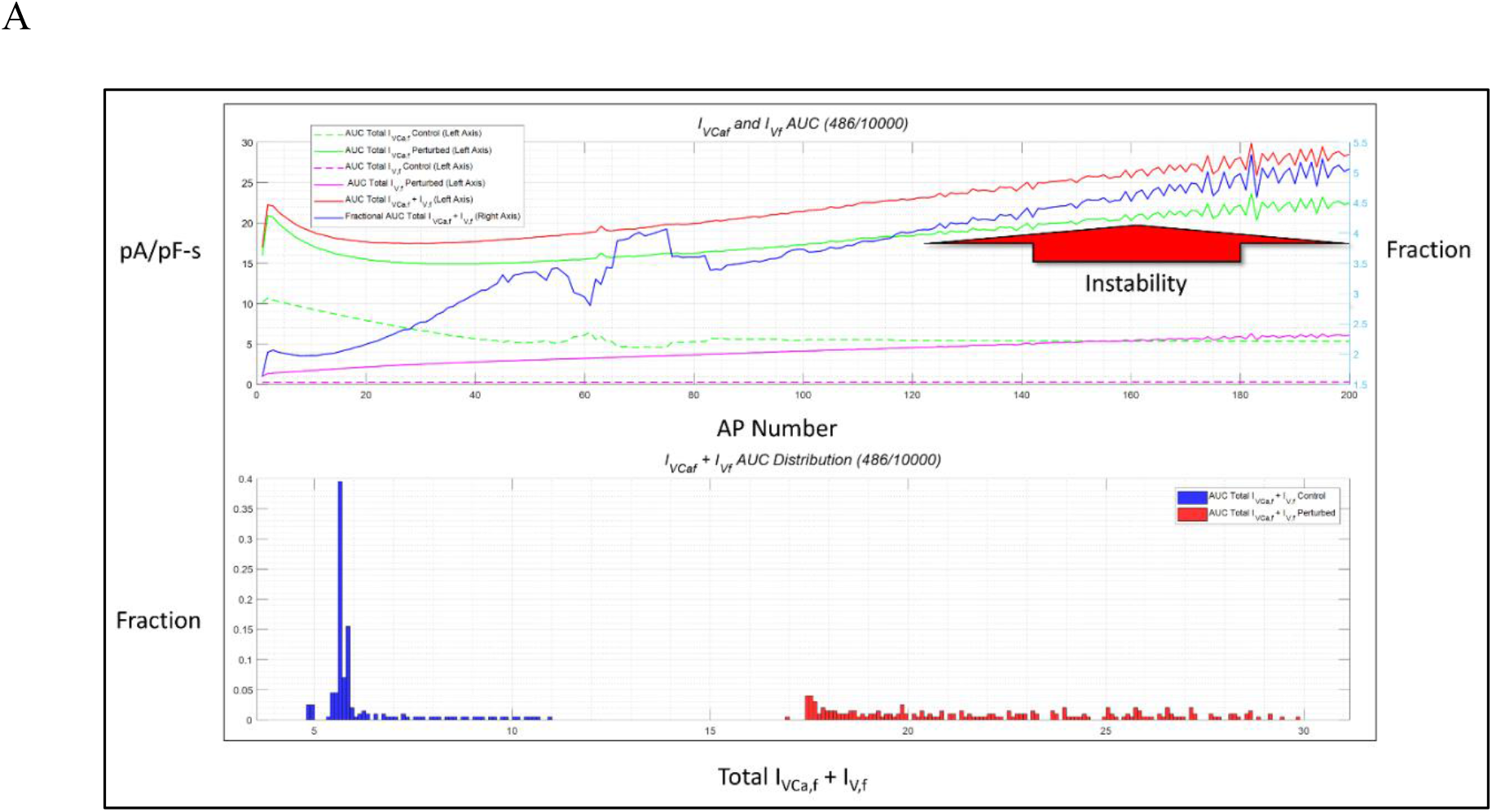
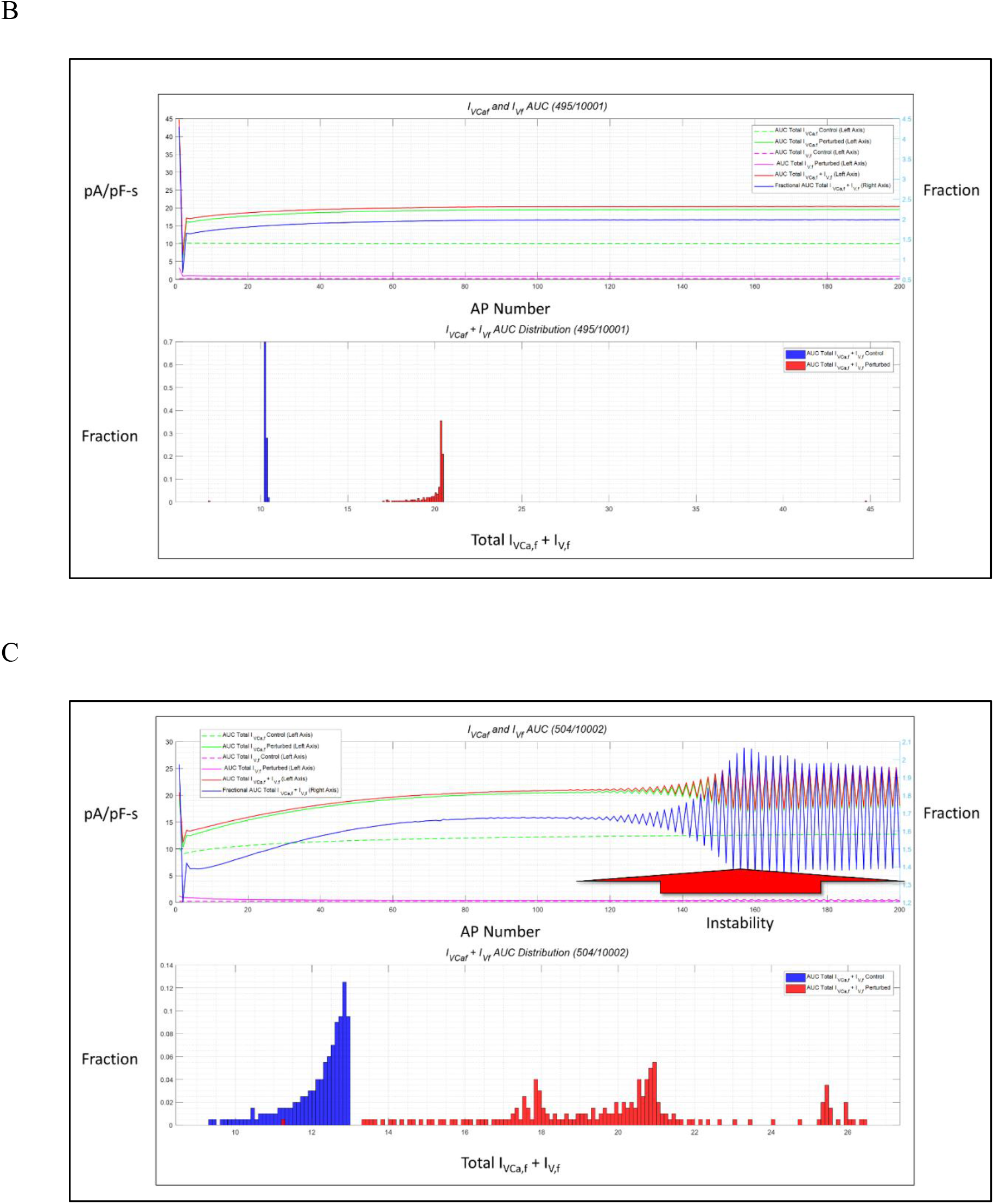
(A) Upper panel: Same as Figure 20A, except showing control versus perturbed |*I_VCa,f_*|*I_V,f_*|, |*I_VCa,f_* + *I_V,f_*|, and (|*I_VCa,f_*| + |*I_V,f_*|) perturbed/control (color coding noted in the legend). Lower panel: Same as Figure 20A, except showing |*I_VCa,f_*| + |*I_V,f_*|. (B) Same as A, except CL = 1/60 min and PT_1_ = 2,597.4567 nM. (C) Same as A, except CL = 1/80 min and PT_1_ = 2,448.6258 nM.

**Figure 23.**
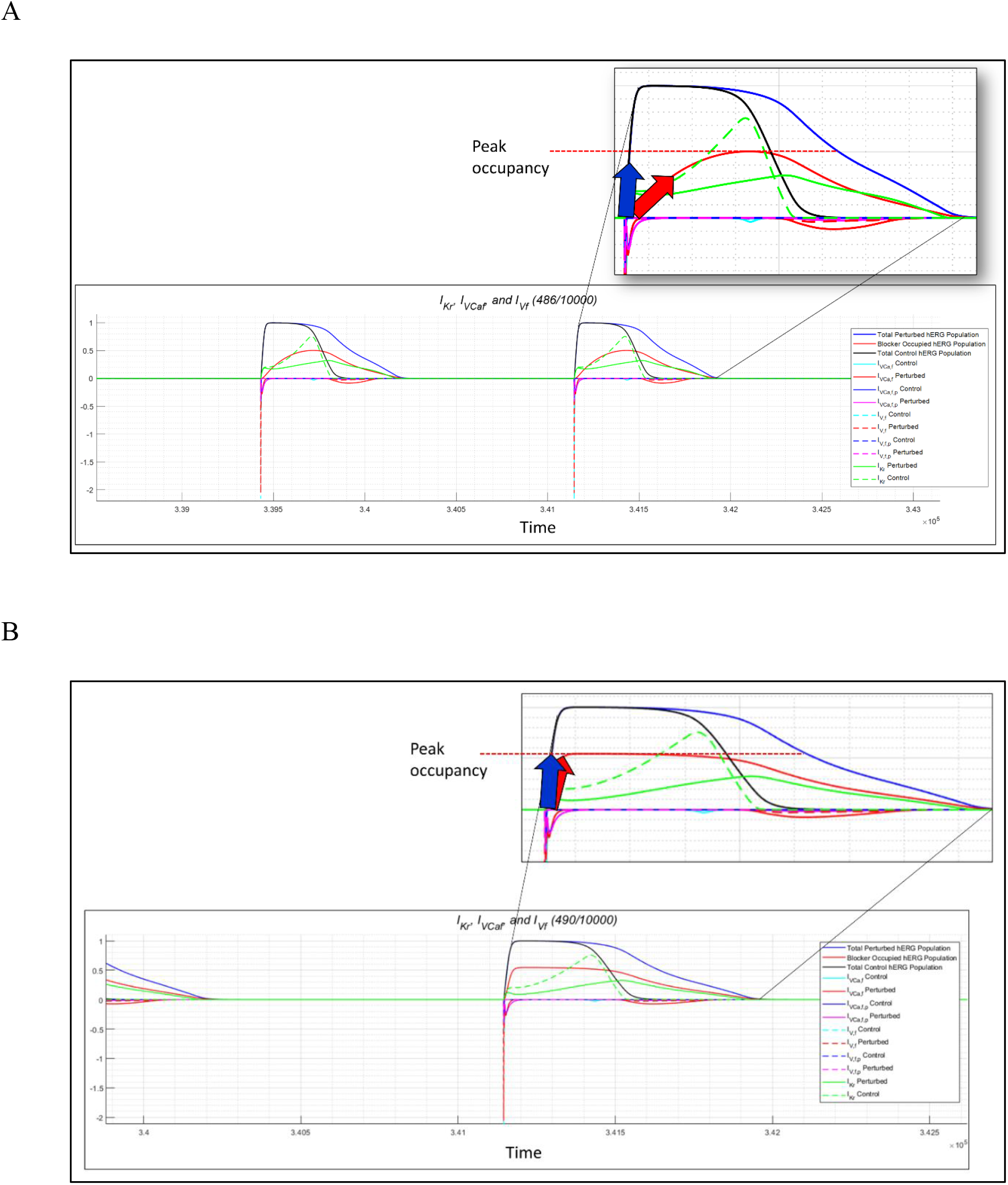
(A) I_Kr_, *I_VCa,f_, I_V,f_*, and fractional blocked and free hERG populations at the 200^th^ AP (mapped to the far right position in the upper panel of Figure 21D). Inset: zoomed in view of the current tracings, showing the rates of buildup of the open + inactivated channel population (blue arrow) and blocker occupancy (red arrow). Peak occupancy is also shown (red dotted line). CL = 1/35 min and blocker k_on_ = 2.0e6 M^-1^ s^-1^, k_off_ = 2.0 s^-1^, blocker concentration = PT_1_ = 2,218.66818 nM (color coding noted in the legend). Late *I_VCa,f_* (solid red tracing in the negative direction) is far above control, I_Kr_ is approximately 50% of control (equal to the unblocked hERG fraction). Buildup of the blocker-occupied channel population (solid red tracing in the positive direction) lags behind channel activation/inactivation at this comparatively slow k_on_, and decay of the occupied population lags behind channel deactivation (solid blue curve), peaking at ~50% of the total population. We hypothesize that arrhythmia is driven by peak, rather than fractional AUC occupancy (AUC of the bound population/AUC of the open + inactivated population ≈ 43% in this scenario). (B) Same as A, except k_on_ = 2.0e6 M^-1^ s^-1^, k_off_ = 2.0 s^-1^, and blocker concentration = PT_1_ = 1,203.3490 nM. The lag in buildup of the bound state is considerably reduced by faster k_on_ compared with that in A, and consequently, the peak occupancy lifetime is longer (recalling that IC_50_ = 1.0 μM in both scenarios). These observations support our claim that occupancy is kinetically, rather than thermodynamically, driven.

Next, we simulated APs for each BK and CL scenario over a series of blocker concentrations ranging from 0 to PT_1_ in 100 nM increments (finer increments were sampled proximal to PT_1_ due to the highly non-linear behavior of the pro-arrhythmic criteria within this regime). The results, which are plotted in Figures 24–25, may be summarized as follows:

1. Maximum hERG occupancy as a function of blocker concentration across each simulation builds to the 50-60% level at PT_1_ according to the Hill equation for all trappable and non-trappable blockers exhibiting k_on_ ≽ 1.0e8 M^-1^ s^-1^. However, occupancy builds more slowly than the rate of channel opening and decays faster than the rate of channel closing for non-trappable blockers exhibiting k_on_ ≤ 1.0e7 M^-1^ s^-1^ (Figure 23A versus Figure 23B), as explained in our previous work [12,19] (reflected in distinct banding in Figure 24A). hERG occupancy drives all pro-arrhythmic effects by tipping the dynamic net inward-outward current balance toward the inward direction (in a graded fashion up to slightly below PT_1_), which in turn, slows Kir2.1 activation, and thereby generates an anomalous I_Ca,L(late)_ window current. At longer CL, I_Ca,L(late)_ builds to the spontaneous depolarizing level, reaching this level at shorter CL via AP_i_-AP_j_ collisions (paced stimulation). Further increases in blocker concentration > T_3_ result in sustained cellular arrhythmia.
2. Maximum APD/CL (Figure 20) grows quasi-linearly across each simulation as a function of blocker occupancy, becoming highly non-linear within a small increment below PT_1_ (Figure 24B). At CL = 1/60 and 1/80 min, APD/CL approaches 1.0 at PT_1_, whereas at CL = 1/35 min, APD/CL approaches ~0.5 at PT_1_, reflecting the differing pro-arrhythmic mechanisms at longer versus shorter CL (Figure 5A). Furthermore, APD/CL grows more slowly with increasing blocker concentration for non-trappable blockers exhibiting k_on_ ≤ 1.0e7 M^-1^ s^-1^ compared with all trappable blockers and non-trappable blockers exhibiting faster k_on_ (e.g. a non-trappable blocker exhibiting k_on_ = 2.0e6 M^-1^ s^-1^ reaches PT_1_ at approximately twice the concentration of a trappable blocker exhibiting the same k_on_ at CL = 1/35 min).
3. A steeper quasi-linear relationship exists between blocker concentration and minimum *d*(Δ*ψ_m_*(*t*))/*dt* across each simulation (Figure 24C), while retaining the biphasic morphology of the other dose-response tracings. At a given blocker concentration, *d*(Δ*ψ_m_*(*t*))/*dt* decreases throughout each simulation (Figure 21D), the overall minimum of which approaches zero at PT_1_. APD and Kir2.1 activation are governed directly, and I_Ca,L(late)_ directly by *d*(Δ*ψ_m_*(*t*))/*dt*.
4. (total |*I_VCa,f_* + *I_V,f_*| perturbed AUC)/(total |*I_VCa,f_* + |*I_V,f_*| control AUC), equating to the fold increase in I_Ca,L(late)_ above control (calculated within a time window between AP start + 150 ms and AP end) serves as a direct window into the act of cellular arrhythmogenesis (Figure 24D). I_Ca,L(late)_ at PT_1_ grows by ~2-fold at CL = 1/60 and 1/80 min and ~5-fold at CL = 1/35 min.
5. Biphasic behavior of hERG blockade dose-response curves is analogous to pH buffering in acid-base titration, in which large perturbations are absorbed until near saturation limit of the system (exemplified in Figure 24E). However, unlike acid-base titration, pro-arrhythmic behaviors undergo high frequency oscillations on approach to PT_1_ (exemplified in Figure 20G), reflecting high instability of the AP generation system.
6. The relationships between the key arrhythmia-conveying parameters *d*(Δ*ψ_m_*(*t*))/*dt* versus (total |*I_VCa,f_* + *I_V,f_* perturbed AUC)/(total |*I_VCa,f_* + *I_V,f_*| control AUC) and hERG occupancy versus (total |*I_VCa,f_* + |*I_V,f_*| perturbed AUC)/(total |*I_VCa,f_* + *I_V,f_*, control AUC) are shown in Figures 25A and B, respectively.
7. Snapshots of key ion current and Ca_v_1.2 gating effects at arbitrary BK profile and sub-PT_1_ hERG blocker concentrations (500 and 1,000 nM) are shown in Figures 26A and B, respectively.

**Figure 24.**
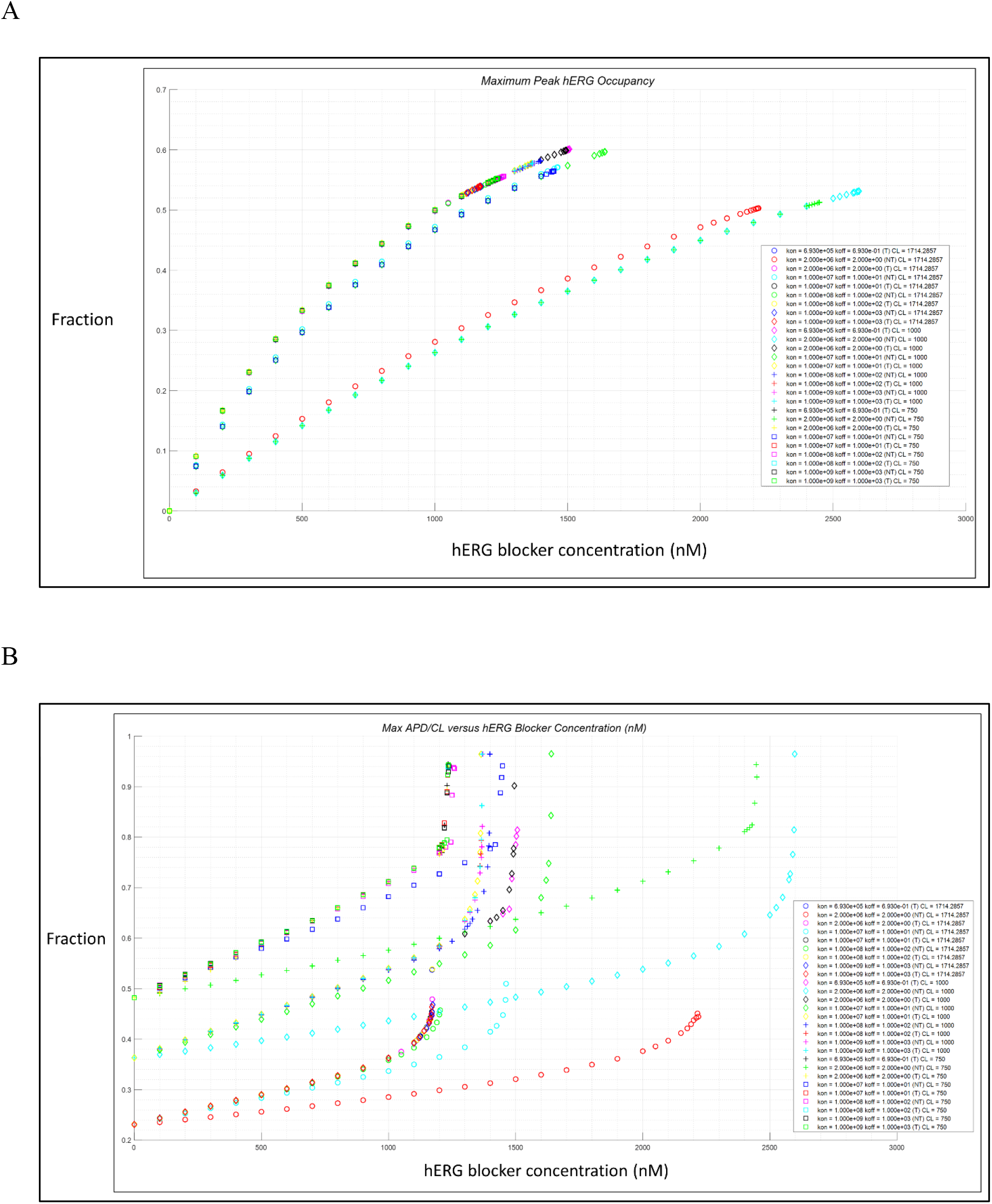
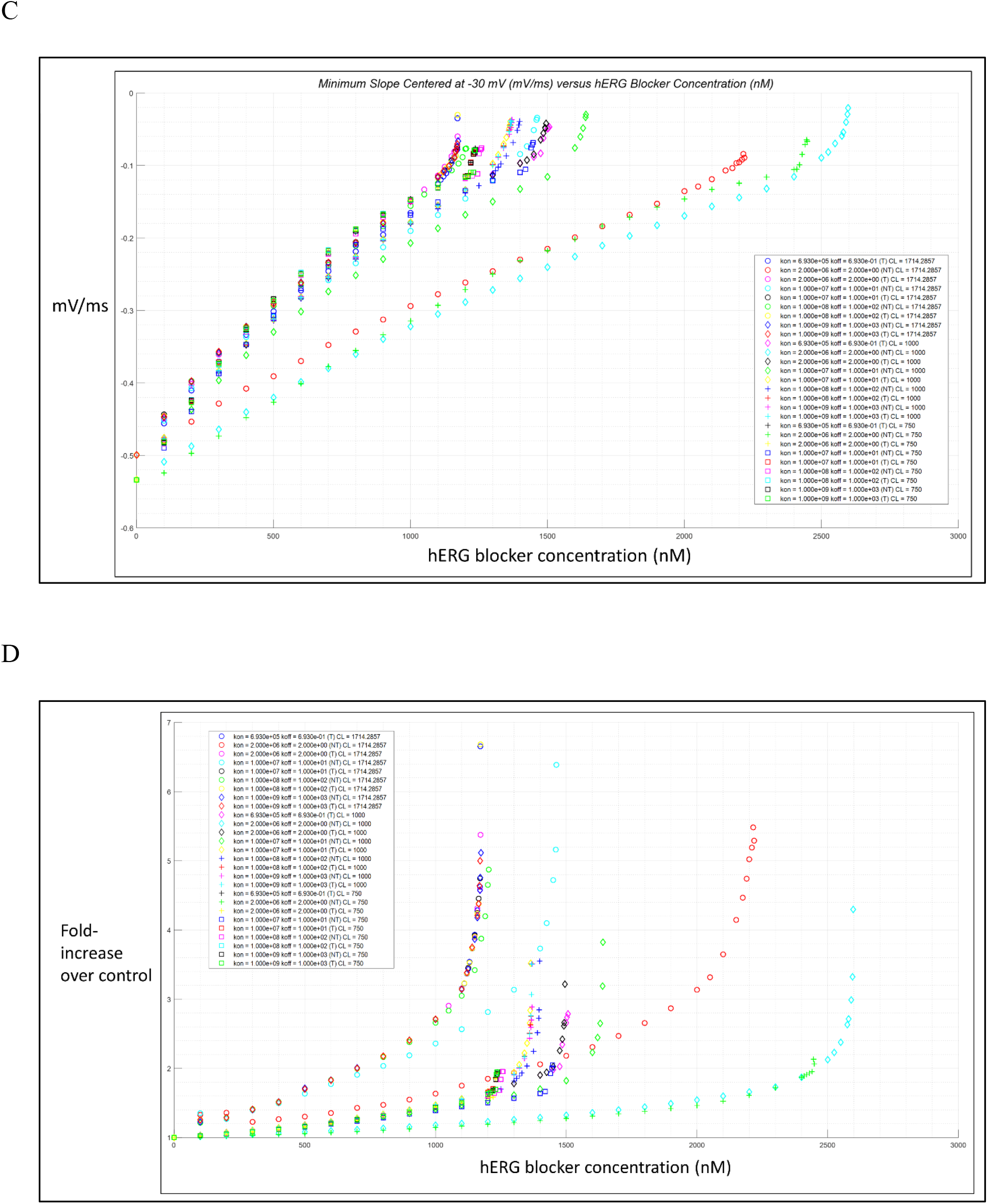
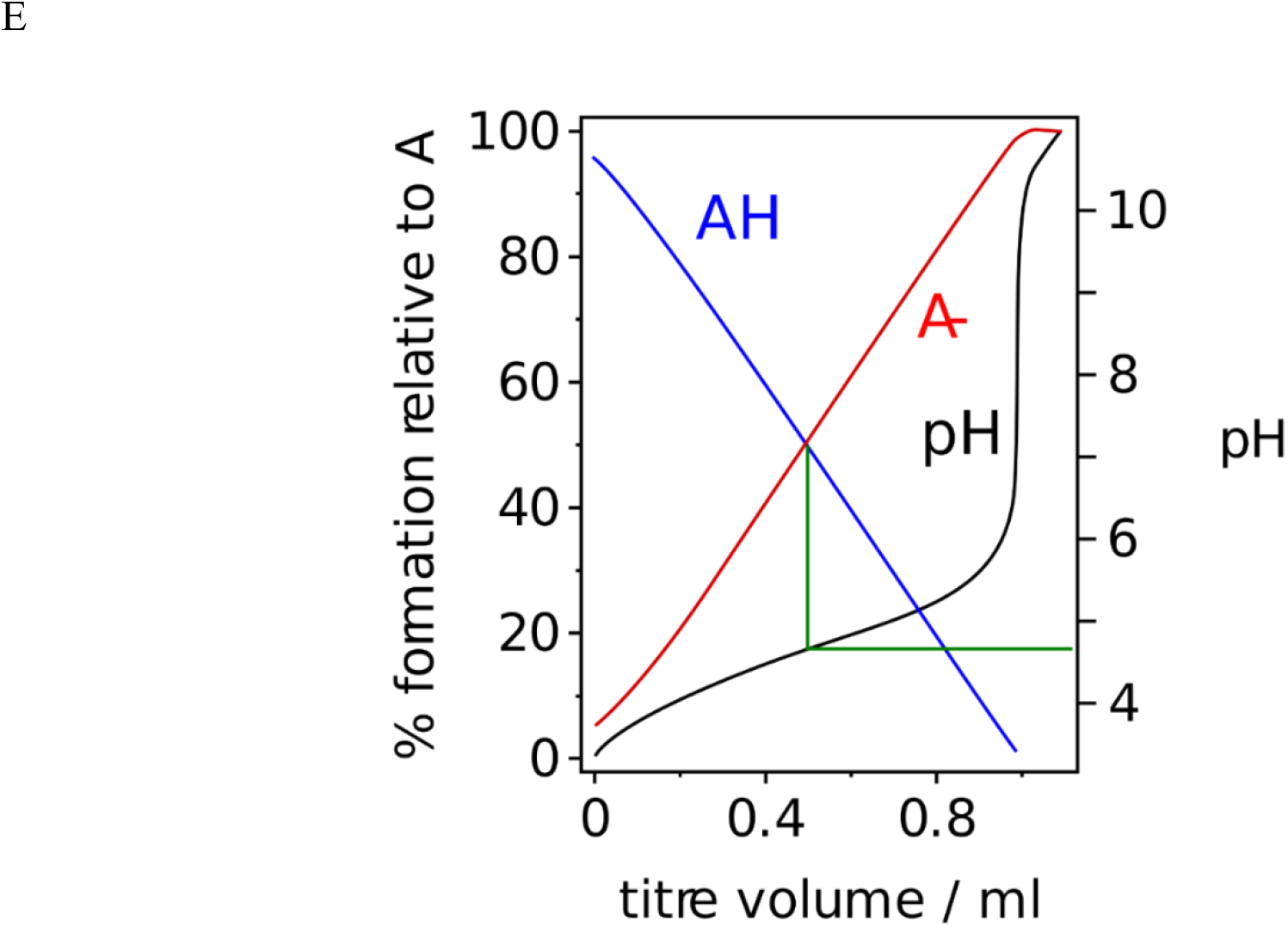
Dose-response tracings for the pro-arrhythmia criteria described in the text. The final concentration in each plot corresponds to PT_1_. (A) Dose-peak hERG occupancy relationships for each of the BK and CL scenarios listed above (color-coding as denoted in the legend). (B) Same as A, except showing dose-APD/CL relationships. (C) Same as A, except showing dose-*d*(Δ*ψ_m_*(*t*))/*dt* relationships. (D) Same as A, except showing dose-(total |*I_VCa,f_* + *I_V,f_*| perturbed)/(total |*I_VCa,f_* + *I_V,f_*| control). (E) The dose-response tracings in B, C, and D resemble generic acid-base titration in the presence of a buffer (Lasse Havelund, https://creativecommons.org/licenses/by-sa/4.0/deed.en).

**Figure 25.**
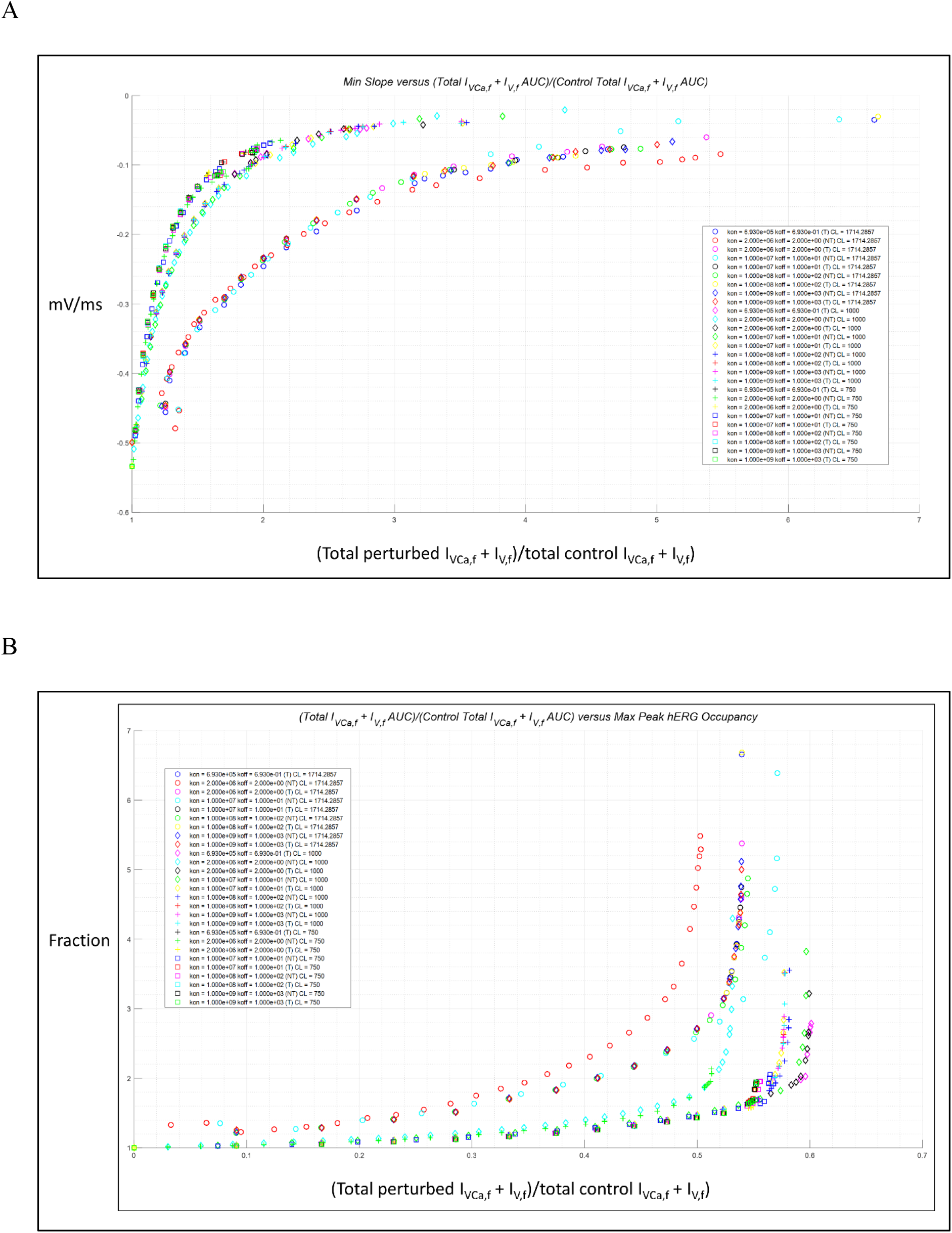
(A) Perturbed minimum *d*(Δ*ψ_m_*(*t*))/*dt* plotted against (total |*I_VCa,f_* + *I_V,f_*| perturbed)/(total |*I_YCa,f_* + *I_V,f_*| control) (color-coding noted in the legend). A quasi-hyperbolic relationship between these quantities is apparent (consistent with the asymptotic approach of *d*(Δ*ψ_m_*(*t*))/*dt* to 0). The different CL-dependent mechanisms of atypical depolarization result in the appearance of two distinct bands in the plot. (B) Peak fractional hERG occupancy plotted against (total |*I_VCa,f_* + *I_V,f_*| perturbed)/(total |*I_VCA,f_* + *I_V,f_*| control). A quasi-hyperbolic relationship between these quantities suggests that hERG occupancy approaches a maximum at the T_1_ threshold of Ca_v_1.2 recovery.

**Figure 26.**
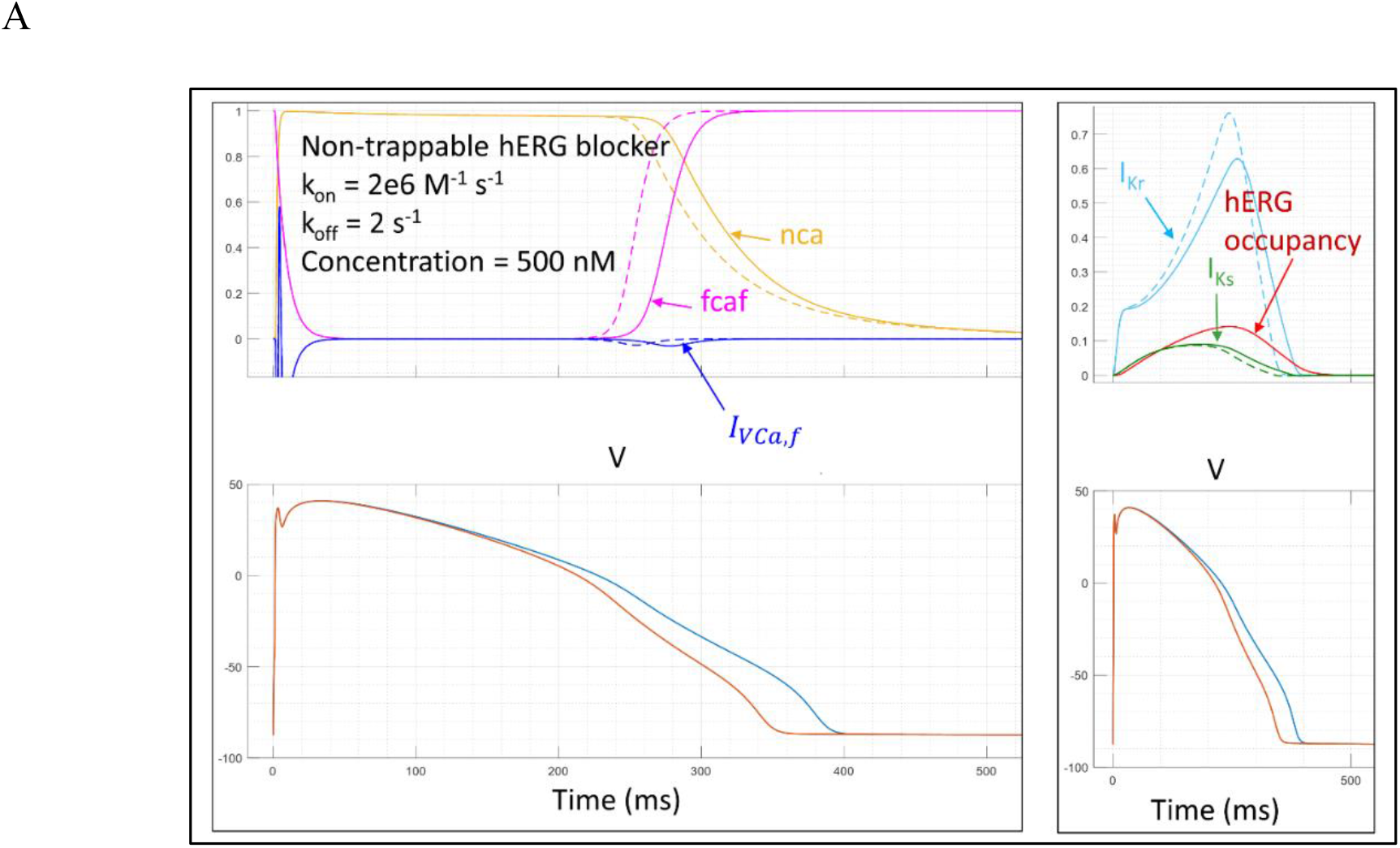
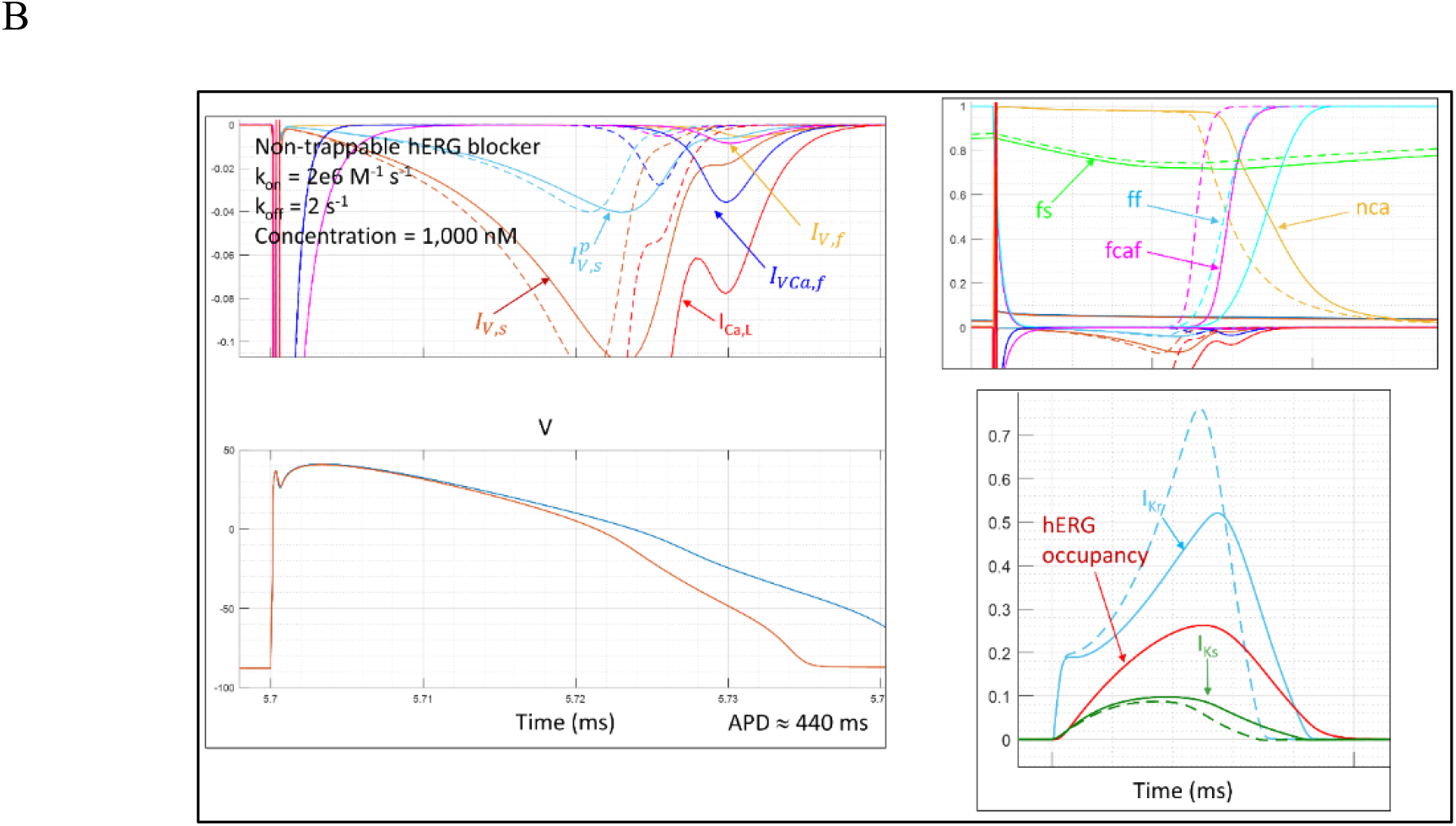
(A) Upper left panel: snapshot of the *I_VCa,f_* window current and fcaf gate under control conditions (dotted tracings) and in the presence of a non-trappable hERG blocker exhibiting k_on_ = 2.0e6 M^-1^ s^-1^ and k_off_ = 2.0 s^-1^ at a concentration = 500 nM (solid tracings). Lower left panel: prolonged and control AP (blue and orange tracings, respectively) corresponding to the tracings in the upper panel. Upper right panel: hERG occupancy (red tracing) I_Kr_ (light blue tracing) and I_Ks_ (green tracing) under control (dotted tracings) and perturbed conditions (solid tracings). Lower right panel: same as lower left panel. (B) Upper left panel: snapshot of all contributions to the I_Ca,L(late)_ window current under control conditions (dotted tracings) and in the presence of the hERG blocker described in A (solid tracings), except at 1,000 nM. *I_VCa,f_* differs the most from control among these currents, accounting for the bump in the I_Ca,L(late)_ tracing that grows to the depolarizing level at the T_1_ threshold of blockade. Lower left panel: same as lower left pane in A. Upper right panel: I_Ca,L(late)_ gating under control (dotted tracings) and perturbed conditions (solid tracings) underlying the anomalous window current. Lower right panel: same as upper right panel in A.

As expected, the T_1_ threshold differs between APD/CL < 1 (corresponding to the spontaneous depolarizing threshold of the I_Ca,L_ window current), and APD/CL → 1 (corresponding to recovery of Ca_v_1.2 channels from the CDI and VDI states triggered by paced AP_i_-AP_j_ collisions). The results of our simulations suggest that the effects of blockade on I_Ca,L_, including the CDI state population level (nca), fcaf gating, *I_VCa,f_* window current, APD/CL, and shoulder *d*(Δ*ψ_m_*(*t*))/*dt* (a putative “pro-arrhythmia fingerprint”) may serve as a more direct metric of pro-arrhythmic propensity than both APD and QT prolongation.

### Na_v_1.5 loss of inactivation function (LOF)-induced arrhythmogenesis in LQT3

Next, we set about to understand the pro-arrhythmic and arrhythmic effects of I_Na(late)_ gain of function (Figure 27), which we simulated by slowing the rate of channel inactivation, as described in Materials and methods (equations 4 and 5). We manually titrated *h_L, ss_* to the PT_1_ threshold via adjustment of *V_shift_* at CL = 1/35, 1/60, and 1/80 min (Table 2). The lowest *V_shift_* at this threshold is achieved at CL = 1/35 min (i.e. bradycardic conditions), consistent with the known trigger of arrhythmia in LQT3 (which is exacerbated by the sinoatrial node suppressing effects of LQT3 mutations [20]). This may be explained by the presence of intact repolarizing currents in this scenario (compared with hERG blockade and LQT2), which affords greater capacity for I_K1_ activation during AP-AP collisions at shorter CL. A snapshot of *I_VCa,f_* at a single prolonged AP at PT_1_ is shown in Figure 28A, and APD/CL and (total |*I_VCa,f_* + |*I_V,f_*| perturbed AUC)/(total |*I_VCa,f_* + *I_V, f_*| control AUC) are plotted over the 200 APs of the simulation in Figure 28B.

**Figure 27.**
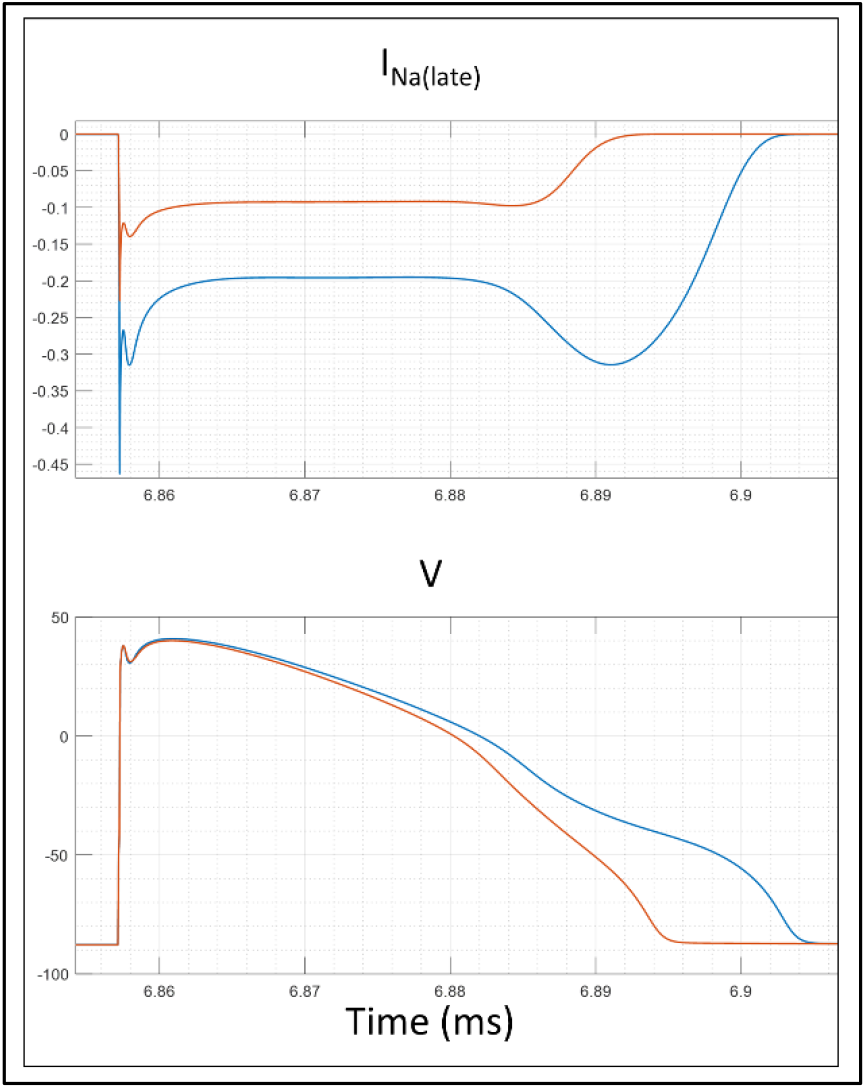
*V_shift_* = −74.1893 at CL = 1/35 min. (A) I_Na(late)_ under perturbed and control conditions (blue and orange tracings, respectively. The maximum perturbed I_Na(late)_ during AP phase 3 is approximately 3-fold greater than control (~3.2 versus −0.1 pA/pF).

**Table 2.**
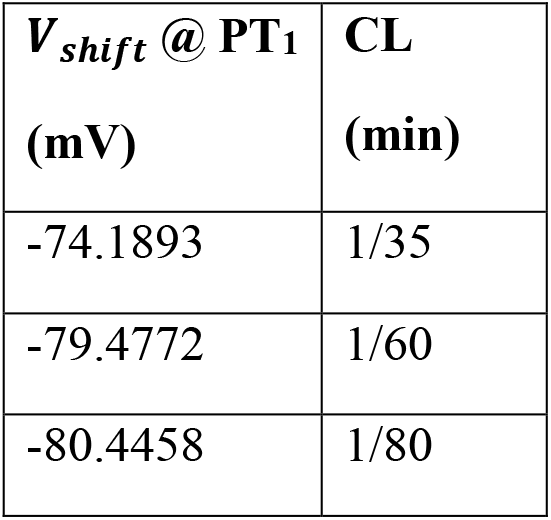
*V_shift_* titration to the PT_1_ level (performed manually, unlike in our hERG blocker doseresponse studies).

| $V_{shift}$ @ $PT_1$<br>(mV) | CL<br>(min) |
| --- | --- |
| -74.1893 | 1/35 |
| -79.4772 | 1/60 |
| -80.4458 | 1/80 |

**Figure 28.**
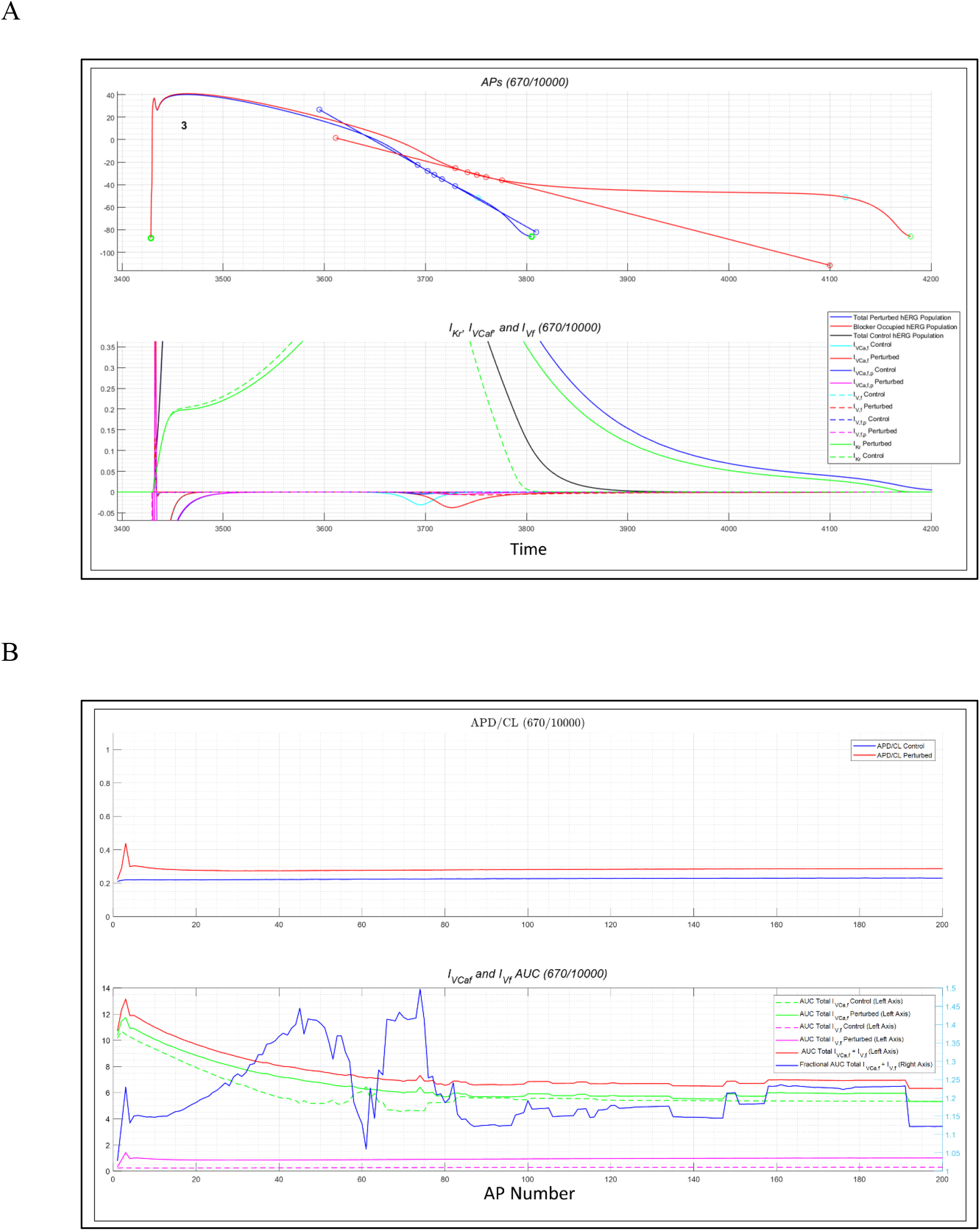
*V_shift_* = −74.1893 at CL = 1/35 min. (A) Upper panel: A representative highly prolonged AP (red) versus control (red and blue, respectively), together with their slopes centered at −30 mV. Lower panel: Repolarizing currents and *I_VCa,f_* (color-coding denoted in the legend). *I_VCa,f_* is significantly larger and broader than control at this *V_shift_*.

### hERG gain of inactivation function (GOF)-induced arrhythmogenesis in LQT2

Next, we set about to understand the mechanism of I_Kr_ knockdown-driven arrhythmia, which we simulated by separately speeding the rate of hERG inactivation (*α_i_*) and decreasing the rate of recovery (*β_i_*) in the Markov model, as described in Materials and methods (equations 1 and 2, respectively). We manually titrated *α_i_* and *β_i_* to the PT_1_ threshold via adjustment of *α_i__V_shift_* and *β_i_-V_shift_* (Tables 3 and 4, respectively). The resulting effects on IKr are shown in Figure 29.

**Table 3.** *α_i__V_shift_* titration to the PT_1_ level.

| $\alpha_i V_{shift}$ @ $PT_1$<br>(mV) | CL<br>(min) |
| --- | --- |
| 65.0538 | 1/35 |
| 66.8892 | 1/60 |
| 63.0618 | 1/80 |

**Table 4.** *β_i__V_shift_* titration to the PT_1_ level.

| $\beta_i V_{shift}$ @ $PT_1$<br>(mV) | CL<br>(min) |
| --- | --- |
| 106.17925 | 1/35 |
| 109.26436 | 1/60 |
| 102.93924 | 1/80 |

**Figure 29.**
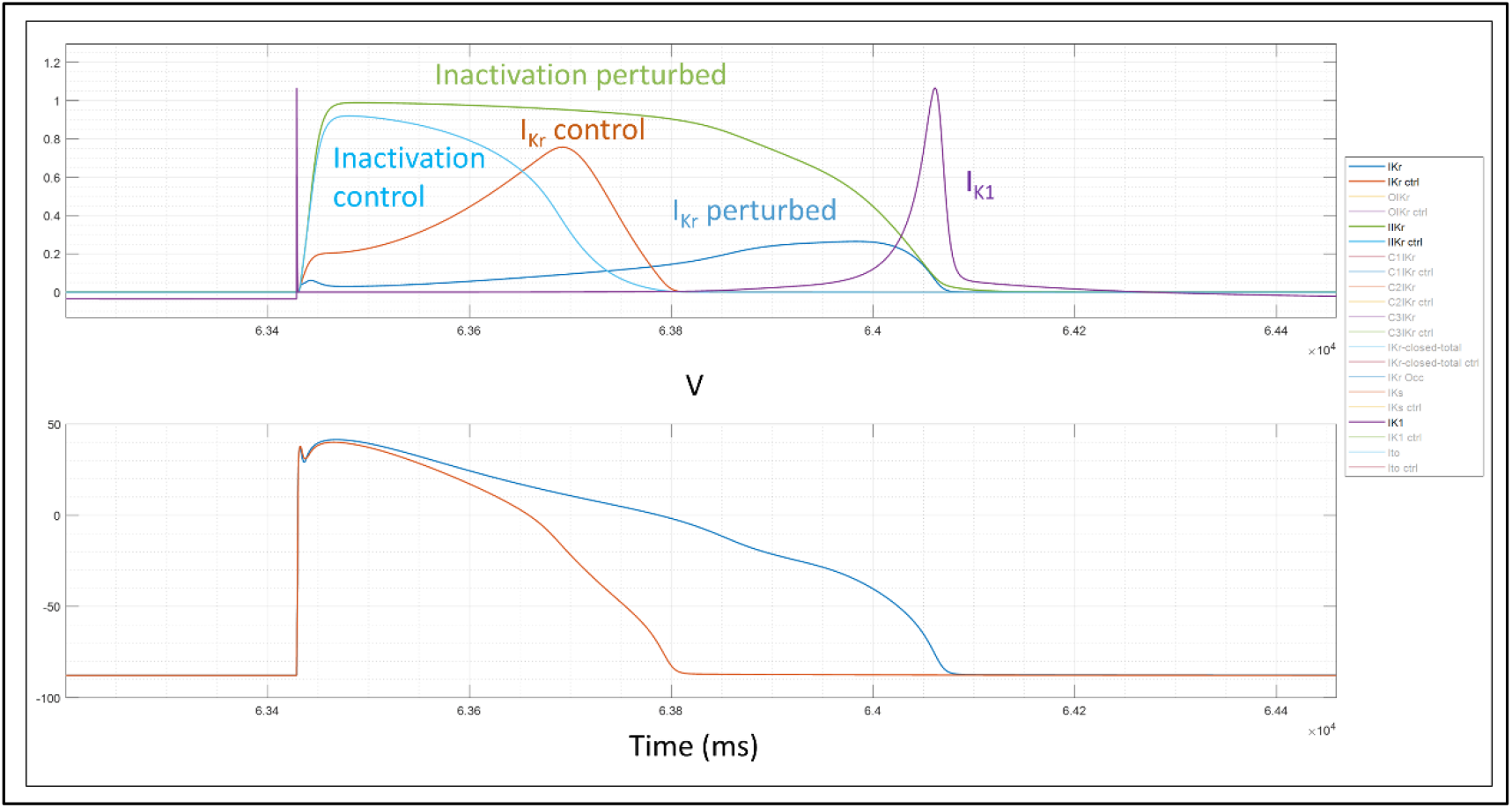
Upper panel: Time-dependent hERG inactivation, I_Kr_, and I_K1_ under control and *α_i_-V_shift_* = PT_1_ = 65.0538 mV (light blue and green, orange and blue, and purple tracings, respectively) conditions at CL = 1/35 min. Lower panel: (Δ*ψ_m_*(*t*) under control and perturbed conditions (orange and blue tracings, respectively). Nearly identical behavior occurs at *β_i__V_shift_* = 106.178925 mV.

As for hERG blockade, the PT_1_ level for both sped inactivation and slowed recovery is greatest at CL = 1/60 min, which further supports the possibility of enhanced arrhythmia resistance at this CL. Furthermore, far greater sensitivity to sped inactivation versus slowed recovery is apparent (noting that these effects are non-mutually exclusive and may occur in combination in LQT2-afflicted individuals). The current effects at PT_1_ are shown in Figure 30.

**Figure 30.**
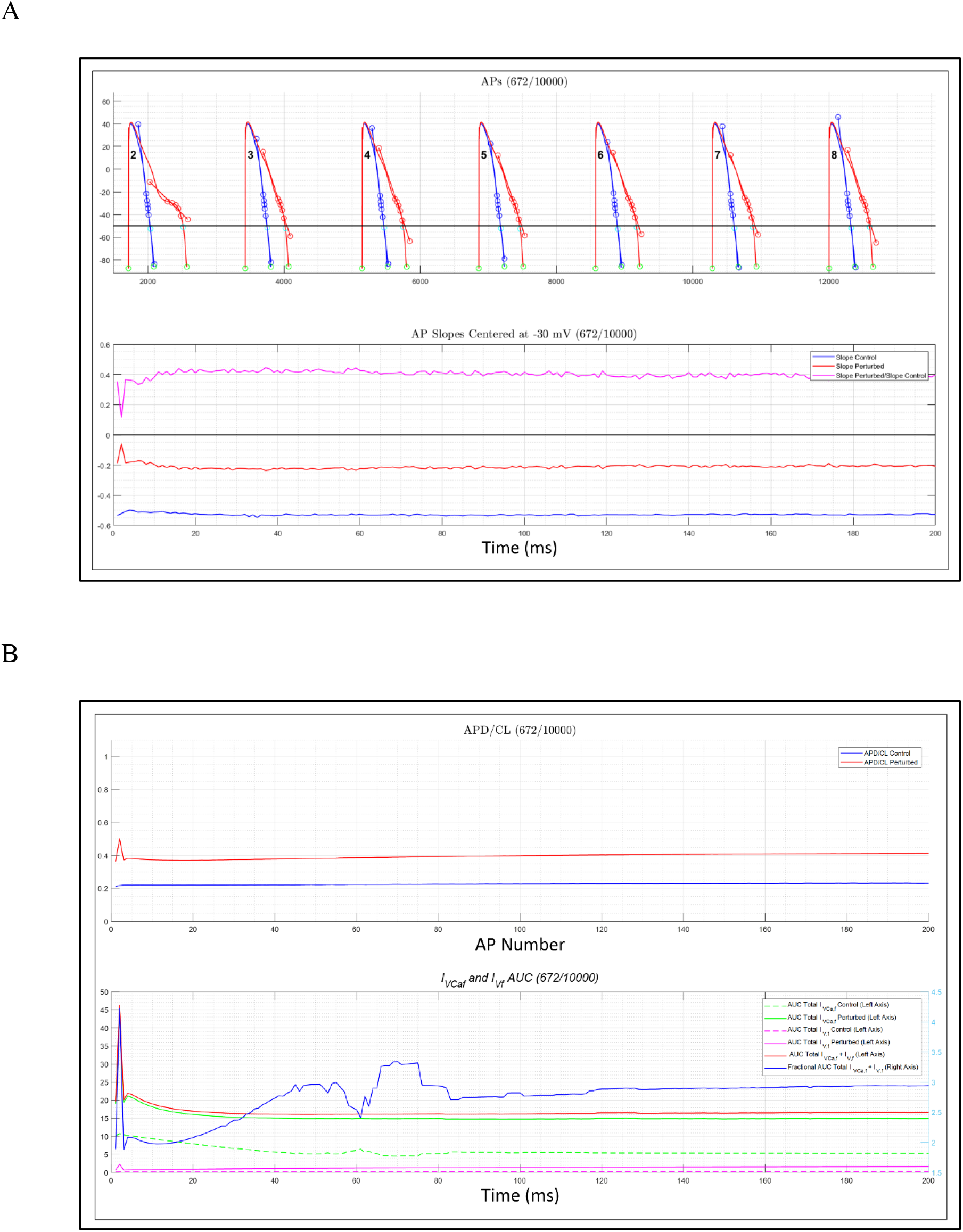
(A) Upper panel: Simulated APs 2-8 under control (blue) versus *α_i__V_shift_* = PT_1_ = 65.0538 mV (red) conditions at CL = 1/35 min, with *d*(Δ*ψ_m_*(*t*))/*dt* tangent lines (red vectors) superimposed on the perturbed APs (slope calculation method described in Materials and methods). The AP numbers in the upper panel (corresponding to those on the x-axis of the lower panel) are shown in bold. Lower panel: Plot of *d*(Δ*ψ_m_*(*t*))/*dt* centered at −30 mV as a function of time. (B) Upper panel: APD/CL as a function of time under control versus perturbed conditions (blue and red tracings, respectively). Lower panel: |*I_VCa,f_*|, |*I_V,f_*|, *I_VCa,f_* + |*I_V,f_*|, and (|*I_VCa,f_*| + |*I_V,f_*|) perturbed/control (color coding noted in the legend).

### Arrhythmogenesis is graded according to hERG blockade, LQT2, and LQT3 severity

Lastly, we examined the progression of arrhythmia at perturbation levels ≥ T_1_ (versus PT_1_) as a function of time and CL = 1/35 and 1/60 min. As described at length in the previous sections, arrhythmogenesis is fueled by the anomalous buildup of I_Ca,L_ (principally *I_VCa,f_ + I_V,f_*) to depolarizing levels as a function of time and perturbation severity, and conveyed by paced AP-AP collisions at CL = 1/35 min and spontaneous depolarizations at CL = 1/60 and 1/80 min. Both forms of pro-arrhythmia similarly progress to full arrhythmia in a graded fashion as a function of time and perturbation severity. Chronic, sub-acute pro-arrhythmic perturbations necessarily reside in the quasi-linear sub-PT_1_ regime, escalating to the T_1_ level and beyond in a condition-dependent manner, either due to downward shifting of the thresholds or increased perturbation severity in the case of hERG blockade (consistent with the stochastic nature of arrhythmia). Examples include changes in CL, hERG blocker exposure due to drug-drug interactions (DDIs)/overdose, physical activity, adrenergic stimulation, CaMKII activity, etc.

Our results suggest that arrhythmogenesis begins with graded instability of the AP generation system. Whereas a background of sub-arrhythmic instability exists below the PT_1_ threshold, further escalation of the perturbation level to PT_1_ results in a shockwave (Figure 20G), in which APD/CL, *d*(Δ*ψ_m_*(*t*))/*dt*, and *I_VCa,f_* + *I_V,f_* undergo high frequency non-arrhythmic oscillations that: 1) settle to a new metastable state at PT_1_; 2) progress to sporadic arrhythmic waveforms at T_1_ and T_2_ (at shorter CL); and 3) progress to lengthy or sustained arrhythmic wave trains (consisting of tetrads of atypical I_Ca,L_-driven depolarization cycles at CL = 1/35 min) at T_3_ (Figure 31). This process is exemplified for simulated LQT3 (noting that hERG blockade and loss of hERG inactivation function behave similarly at this CL), as follows:

1. Paced + non-paced AP dyads occur sporadically at perturbation levels ≥ T_1_ and CL = 1/35 min (Figure 32A).
2. AP triads and tetrads resulting from additional atypical I_Ca,L_-driven depolarizations (reflecting oscillations in Ca_v_1.2 inactivation gating shown in Figure 14) appear over time (Figures 32B and 32C).
3. The T_2_ threshold is crossed when the wavelength of arrhythmic wave packets expands to the CL (corresponding to total loss of restitution time) (Figure 32C), at which point collisions with succeeding paced APs occur (Figure 32D).
4. The wave packet wavelength further expands at the T_3_ threshold to >> CL (Figure 33), becoming continuous at perturbation levels >> T_3_ (Figure 34).

**Figure 31.**
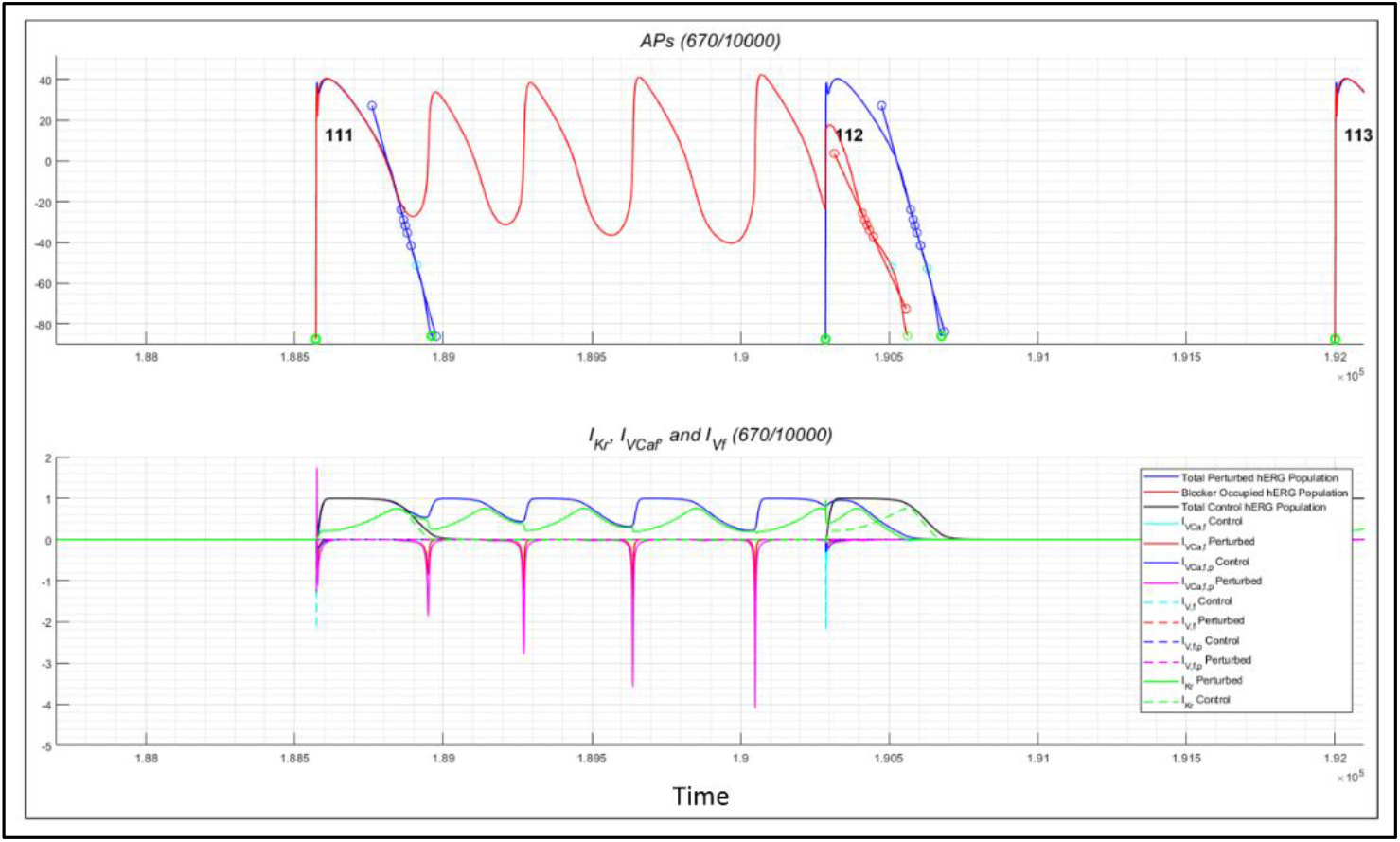
*V_shift_* = −77.0 (simulated LQT3) at CL = 1/35 min. (A) Upper panel: Arrhythmic wave packets that are terminated by intact I_Kr_, I_Ks_, and I_K1_ activation at this perturbation level. Lower panel: “Daisy-chained” repolarizing currents above the T_1_ threshold, which consist principally of *I_VCa,f_*, and *I_V,f_* (color-coding denoted in the legend).

**Figure 32.**
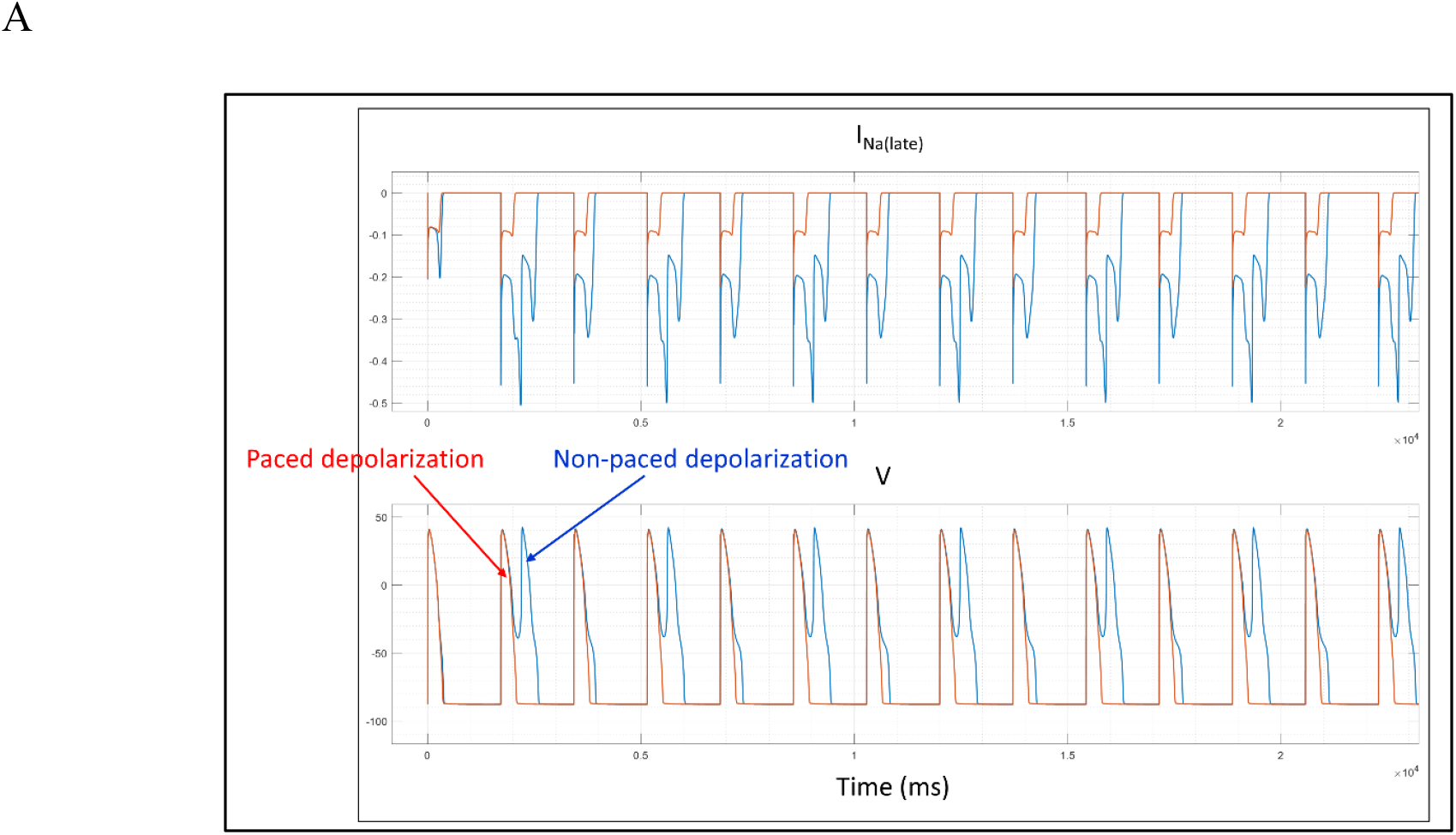
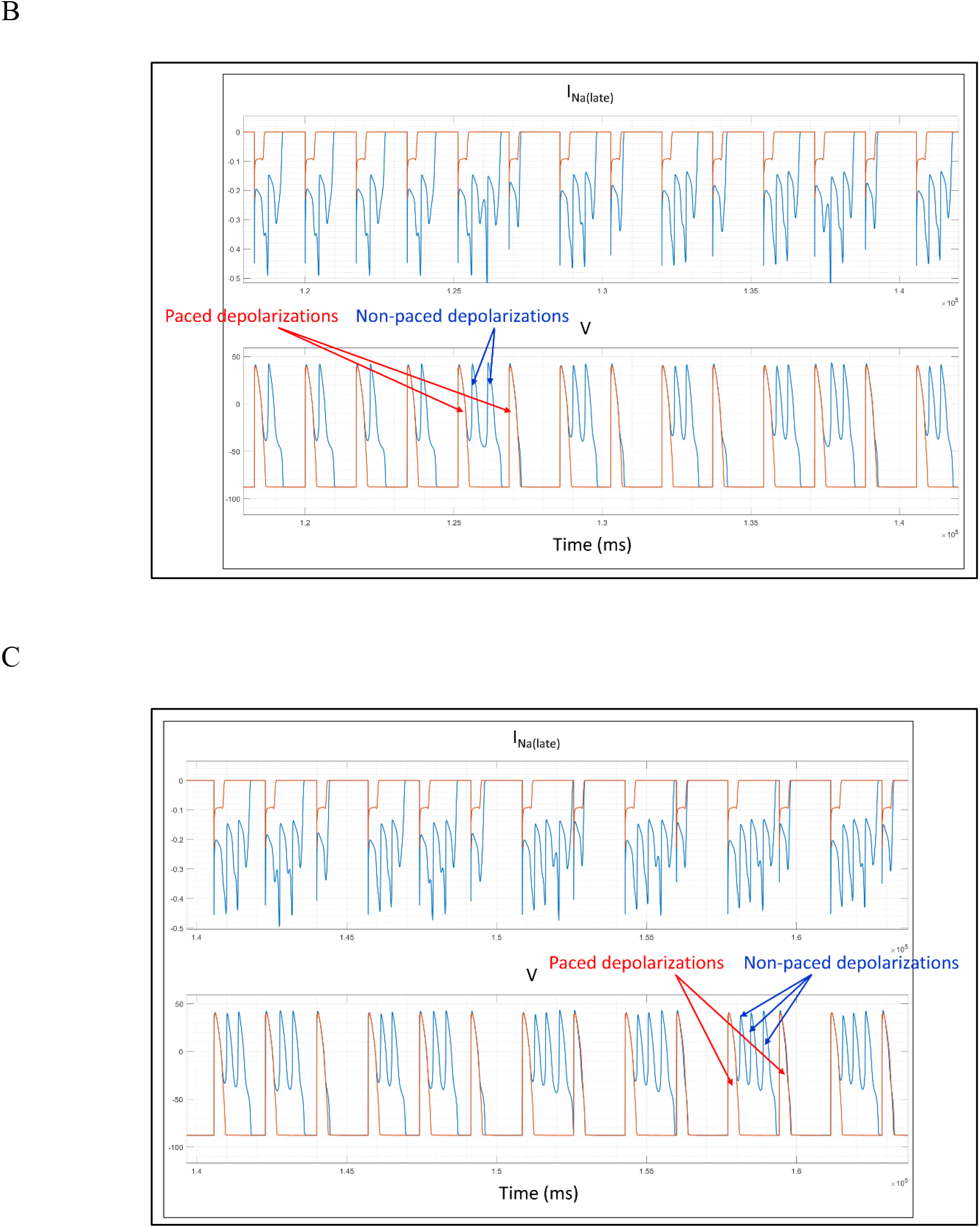
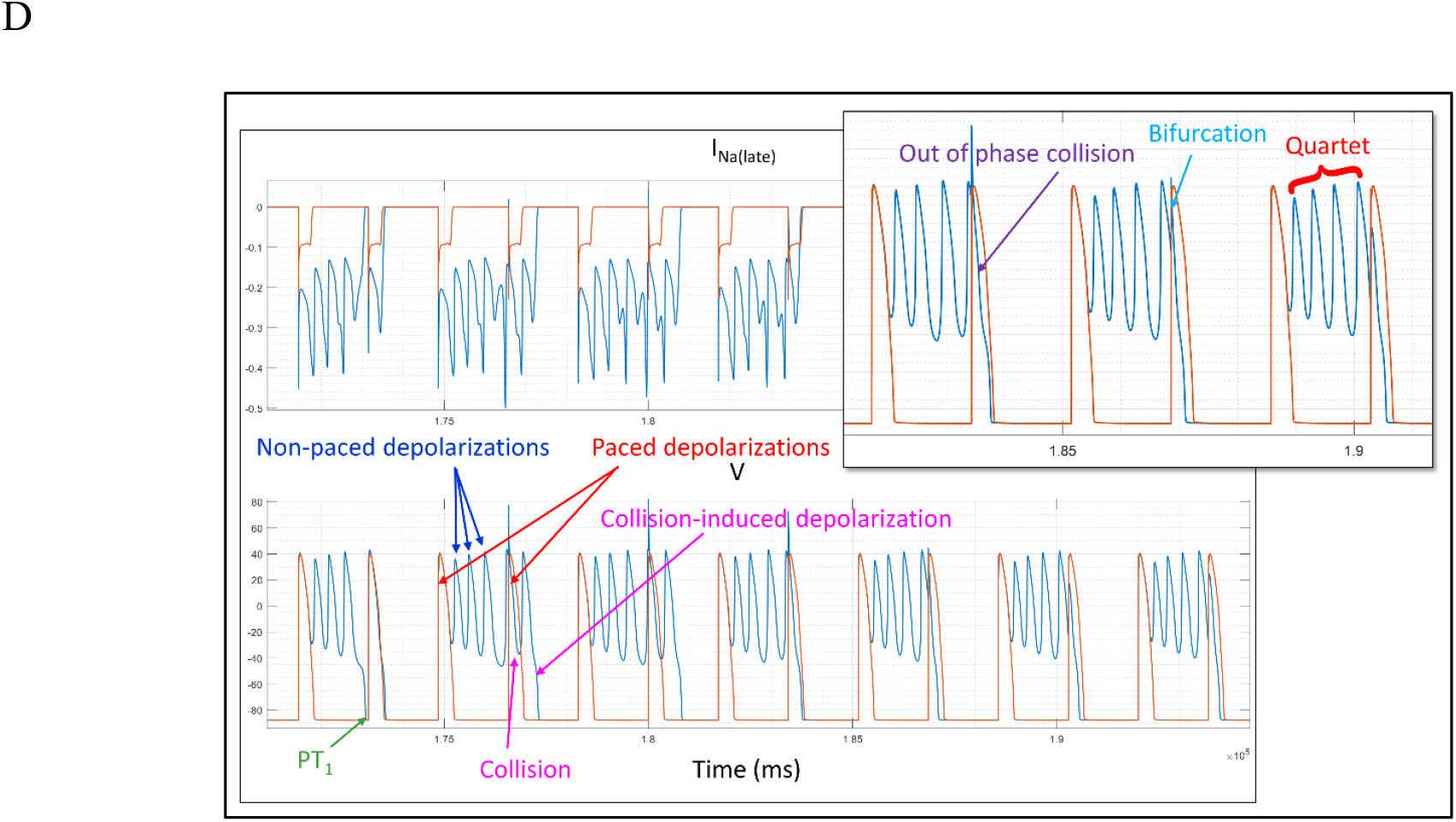
LQT3 *V_shift_* = −77.0 mV (2.81 mV above PT_1_) at CL = 1/35 min. Upper panels: I_Na(late)_ under perturbed and control conditions (blue and orange tracings, respectively). Lower panels: Δ*ψ_m_*(*t*) under perturbed and control conditions (blue and orange tracings, respectively). At this perturbation level, the intra-packet frequency of spontaneous depolarizations increases over time from singlets (B) to doublets (C), triplets (D), and quartets (E) sandwiched between pairs of consecutive paced APs. Quartets are formed by out-of-phase collisions, resulting in bifurcations of intra-packet waveforms, as noted in the inset.

**Figure 33.**
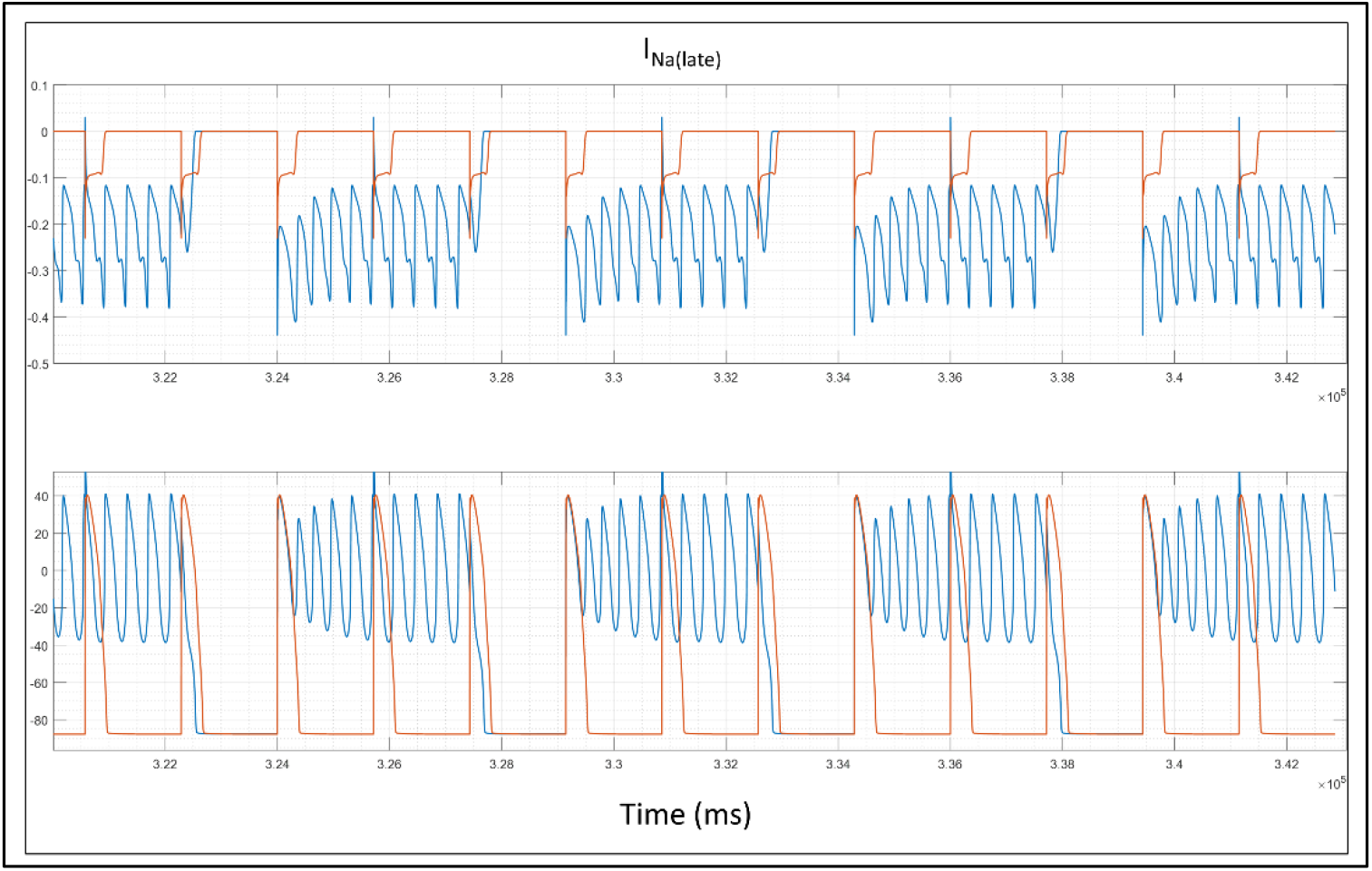
Same as Figure 29, except *V_shift_* = −79.0 mV (4.81 mV above PT_1_) at CL = 1/35 min. The intra-wave packet wavelength and frequency expand across three paced APs at this perturbation level, resulting in additional collisions and bifurcations.

**Figure 34.**
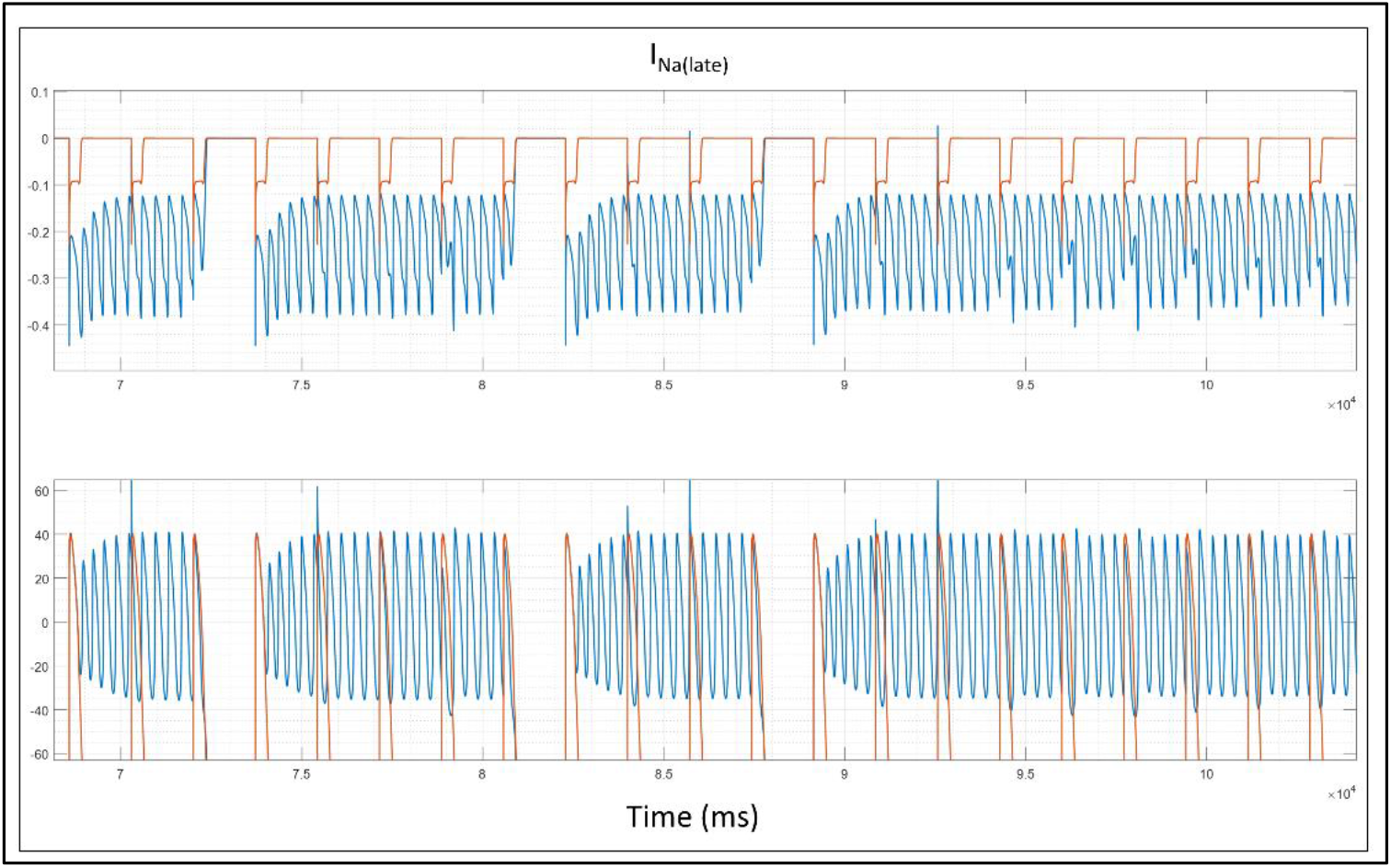
Same as Figure 29, except *V_shift_* = −83.0 mV (8.81 mV above PT_1_) at CL = 1/35 min. The intra-wave packet wavelength and frequency expand continuously across the paced APs at this perturbation level.

Atypical spontaneous ICa,L-driven depolarizations give way to pacing-triggered depolarizations at CL = 1/60 and 1/80 min. The T_1_ and T_2_ stages of this process are shown for hERG blockade in Figures 8 and 35.

**Figure 35.**
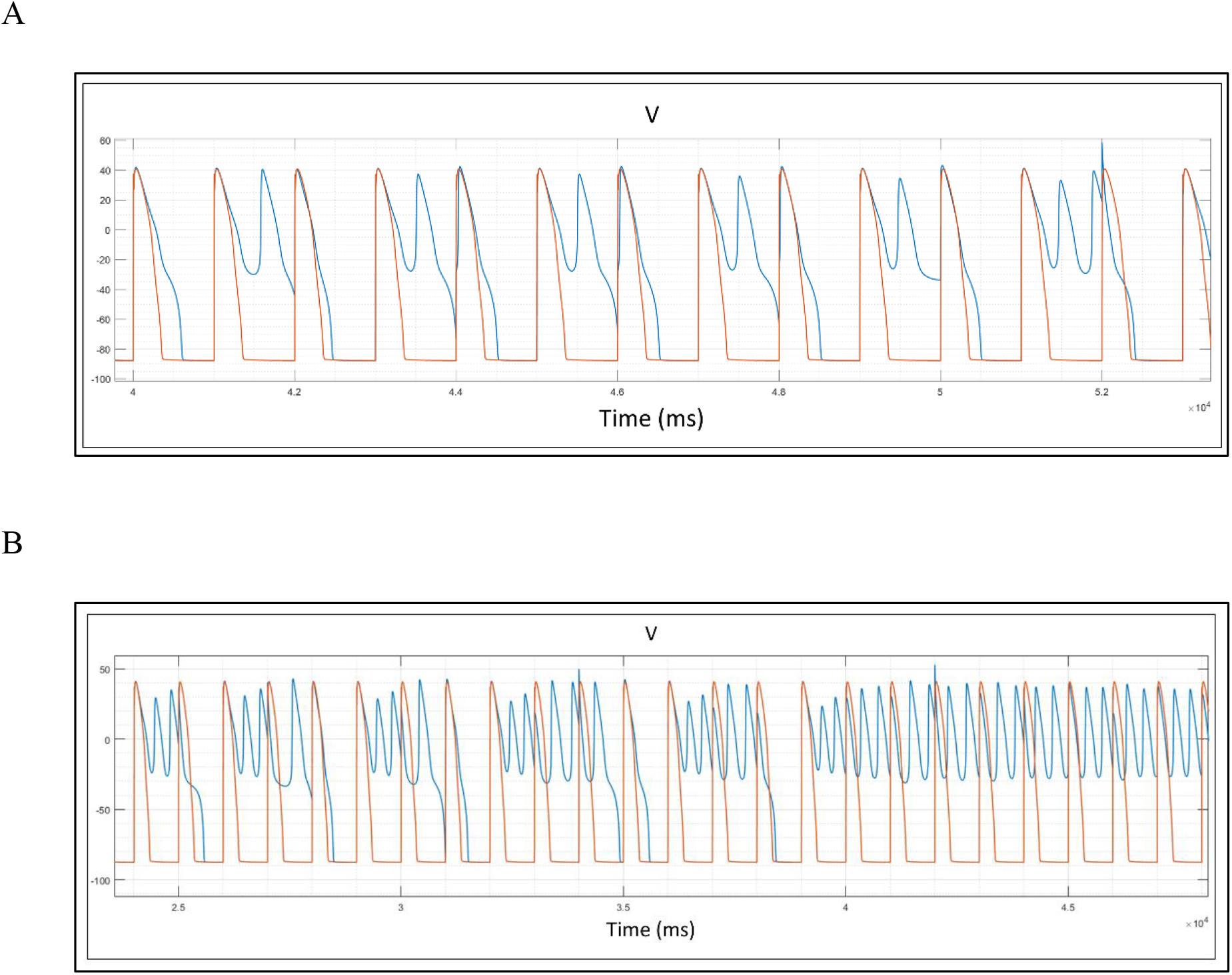
(A) The time-dependent progression of arrhythmic waveforms at CL = 1/60 min and hERG blocker concentration = 60.235 nM (k_on_ = 1e8, k_off_ = 2.0) (AP 1) to single pacing-triggered atypical I_Ca,L_-driven depolarizations (wave packets 2-5) to collision-induced AP dyads (wave packet 7). (B) Same as A, except hERG blocker concentration = 65.0 nM. The APs progress from a collision-induced bifurcation (wave packet 1), which evokes additional pacing-triggered atypical depolarizations (wave packets 2 and 3). This process further expands (wave packet 4), becoming continuous at wave packet 6.

## Discussion

### The implications of our findings for electrical homeostasis

We recently described a first principles physics-based theory (which we refer to as “Biodynamics” [1,21]) describing the general paradigm by which:

1. Cellular function (biochemical and electrical) is generated by molecular systems.
2. Molecular systems respond to perturbations.
3. Cellular dysfunction results from molecular dysfunction or excessive perturbation.

Here, we have used Biodynamics to explain arrhythmogenesis in, as well as the natural anti-arrhythmic/homeostatic properties of, ventricular cardiomyocytes based on perturbation-induced alterations in the dynamic net inward-outward current balance as a function of CL. Our results suggest that cellular homeostasis/arrhythmia resistance depends first and foremost on a Goldilocks zone of APD/CL across all CL (0.23, 0.36, and 0.48 under control conditions in our simulations at pacing CL = 35, 60, and 80 BPM, respectively). APDs shorten and lengthen with decreased and increased pacing CL, respectively. APD is governed by processes operating at both the intra- and extra-cycle levels (the latter of which are neglected in the ORd model). Intra-cycle processes include CaMK-II-mediated phosphorylation and phosphatase-mediated dephosphorylation of ion channels and RyR_2_ (which speed and slow inactivation, respectively), whereas extra-cycle processes (acting on timescales >> a single CL) include β2-mediated PIP_2_ levels and PKA-mediated phosphorylation. At a given CL (which depends strictly on pacing frequency), APD/CL depends directly on the degree to which the dynamic net inward-outward current balance tips in the inward versus outward direction, thereby lengthening and shortening the APD, respectively. The degree of net tipping, in turn, is governed by the rates of inwardly versus outwardly conducting ion channel activation versus inactivation/deactivation state transitions. APD per se, is governed largely by the Kir2.1 activation rate, which, in turn, depends on the rate at which Δ*ψ_m_*(*t*) repolarizes to < −50 mV (governed by *d*(Δ*ψ_m_*(*t*))/*dt* during AP phase 3).

Anomalous decrease of APD/CL due to severe bradycardia and/or APD shortening beyond nominal limits promotes I_Ca,L_ buildup to non-paced depolarizing levels. Conversely, anomalous increase of APD/CL due to severe tachycardia and/or APD expansion beyond nominal limits promotes collisions between successive paced APs (described by Fossa [22] as the loss of restitution time). Hondeghem et al. proposed that pro-arrhythmic versus non-pro-arrhythmic hERG blockade are distinguishable based on a set of criteria consisting of triangulation of the AP morphology, reverse use dependence of APD, instability, and dispersion of repolarization (“TRIaD”) [23]. Although our findings suggest that pro-arrhythmic propensity is governed solely by dynamic fractional ion channel occupancy (not necessarily limited to hERG), the TRIaD concept can be viewed as the phenotypic counterpart of our proposed pro-arrhythmia criteria, as follows:

1. Perturbation-induced changes in AP morphology (not necessarily triangulation). Our findings suggest that AP lengthening follows from decreased *d*(Δ*ψ_m_*(*t*))/*dt* in the shoulder region on approach to T_1_. APD prolongation is graded as a function of perturbation level, resulting in isomorphic (putatively anti-arrhythmic) and nonisomorphic (certainly pro-arrhythmic) variations at smaller versus larger prolongation levels, respectively. Our results suggest that therapeutic APD prolongation by class III anti-arrhythmic drugs necessarily falls within a Goldilocks zone between anti-arrhythmic increases in refractoriness versus pro-arrhythmic increases in APD/CL, and concomitant loss of restitution time. Our simulations clearly suggest that far greater levels of hERG blockade are tolerable than is widely assumed, consistent with the existence of a therapeutic index for class III hERG blocking anti-arrhythmic agents.
2. The susceptibility of Ca_v_1.2 to undergo spontaneous recovery from inactivation at slower HR/longer CL is consistent with reverse use dependence.
3. Instability, corresponding to oscillations in *d*(Δ*ψ_m_*(*t*))/*dt* and Ca_v_1.2 recovery within the PT_1_ regime.
4. Dispersion, corresponding to differential effects of perturbation on APDs in M, epi- and endocardial cells, manifesting as increased wavelength of the T wave.

## Conclusion

We used ORd/hERG Markov to simulate the progression of normal cardiac APs to the arrhythmic state under conditions of hERG blockade/loss of function and Na_v_1.5 gain of function, focusing on the underlying ion current anomalies driving the process. We describe the AP generation mechanism on the basis of Biodynamics theory (dynamic inward-outward current balance optimized for stimulus-response based electrical behavior and excitation-contraction coupling) and arrhythmogenesis (anomalous dynamic inward-outward current balance that promotes spontaneous depolarizations and AP-AP collisions). The major findings can be summarized as follows:

1. Arrhythmogenesis (irrespective of etiology) may be partitioned into graded pro-arrhythmic, pre-arrhythmic, and arrhythmic regimes as a function of perturbation severity and time, such that abrupt inter-regime transitions occur over tiny perturbation increments (somewhat analogous to buffered acid-base titration).
2. All phases of arrhythmogenesis depend on pacing CL. Longer CL/slower heart rates promote spontaneous atypical ICa,L-driven depolarizations (specifically due to anomalous recovery from the CDI and VDI states of Ca_v_1.2), whereas shorter CLs promote collisions between successive APs that result in pacing-induced I_Ca,L_-driven depolarizations. Arrhythmogenesis is thus governed by interactions between paced and non-paced APs (Figures 32–35). Arrhythmic AP wavelengths expand via collisions between spontaneous non-paced and paced depolarizations (noting that collision frequency grows with increasing wavelength above the pacing CL). This process culminates in continuous oscillations spanning across the diastolic interval of each cycle.
3. The AP generation system responds to increasing perturbation levels in a quasi-linear fashion throughout the pro-arrhythmic regime (Figure 24). However, non-linear progression to the arrhythmic state is tacitly assumed under the current hERG blocker safety assessment protocol. Our results reiterate our previous claim that significantly greater exposure is required to achieve pre-arrhythmic occupancy for non-trappable blockers exhibiting slower k_on_, versus trappable blockers or non-trappable blockers with faster k_on_ (at equivalent IC_50_) (Figure 23). Revision of the hERG safety margin may be warranted based on our simulated mechanism-based dose-response results (further addressed in [18]), which furthermore, could facilitate more straightforward consideration of benefit/risk criteria in the hERG safety equation.

